# Exploring Integrin α5β1 as a Potential Therapeutic Target for Pulmonary Arterial Hypertension: Insights from Comprehensive Multicenter Preclinical Studies

**DOI:** 10.1101/2024.05.27.596052

**Authors:** Sarah-Eve Lemay, Monica S. Montesinos, Yann Grobs, Tetsuro Yokokawa, Tsukasa Shimauchi, Charlotte Romanet, Mélanie Sauvaget, Sandra Breuils-Bonnet, Alice Bourgeois, Charlie Théberge, Andréanne Pelletier, Reem El Kabbout, Sandra Martineau, Keiko Yamamoto, Adrian S. Ray, Blaise Lippa, Bryan Goodwin, Fu-Yang Lin, Hua Wang, James E Dowling, Min Lu, Qi Qiao, T. Andrew McTeague, Terence I. Moy, François Potus, Steeve Provencher, Olivier Boucherat, Sébastien Bonnet

## Abstract

Pulmonary arterial hypertension (PAH) is characterized by obliterative vascular remodeling of the small pulmonary arteries (PA) and progressive increase in pulmonary vascular resistance (PVR) leading to right ventricular (RV) failure. Although several drugs are approved for the treatment of PAH, mortality remains high. Accumulating evidence supports a pathological function of integrins in vessel remodeling, which are gaining renewed interest as drug targets. However, their role in PAH remains largely unexplored. We found that the arginine-glycine-aspartate (RGD)-binding integrin α5β1 is upregulated in PA endothelial cells (PAEC) and PA smooth muscle cells (PASMC) from PAH patients and remodeled PAs from animal models. Blockade of the integrin α5β1 or depletion of the α5 subunit resulted in mitotic defects and inhibition of the pro-proliferative and apoptosis-resistant phenotype of PAH cells. Using a novel small molecule integrin inhibitor and neutralizing antibodies, we demonstrated that α5β1 integrin blockade attenuates pulmonary vascular remodeling and improves hemodynamics and RV function in multiple preclinical models. Our results provide converging evidence to consider α5β1 integrin inhibition as a promising therapy for pulmonary hypertension.

**One sentence summary:** The α5β1 integrin plays a crucial role in pulmonary vascular remodeling.

## INTRODUCTION

Pulmonary arterial hypertension (PAH) is a multifactorial and progressive disease characterized by the persistent elevation of the mean pulmonary artery (PA) pressure leading to death from right ventricular (RV) failure ^1, 2^. The pathogenic features leading to PAH include sustained vasoconstriction and extensive vascular remodeling and stiffening. These changes are primarily due to excessive proliferation and survival of PA-resident cells associated with extracellular matrix (ECM) accumulation and changes in mechanical properties. Although there are currently marketed therapies, aimed at restoring the vasoconstriction/vasodilation balance, delaying disease progression, and improving survival, they have only limited ability to reverse pulmonary vascular remodeling and do not offer a cure ^3, 4^. The recent FDA approval of sotatercept represents a significant step towards potentially targeting vascular remodeling in PAH and is expected to benefit many patients ^5, 6^. However, its clinical mechanism of action, particularly the extent of its anti-remodeling effects, is still under investigation and the long-term safety profile is unclear ^5, 7^. In this context, the identification of new druggable targets specifically involved in the pathological remodeling of the pulmonary vasculature and right ventricle represent a promising therapeutic direction.

The process of pulmonary vascular remodeling is complex, involving all three layers (intima, media, and adventitia) of the PA wall. Pulmonary artery smooth muscle cells (PASMC) and pulmonary artery endothelial cells (PAEC) are central to this process, actively contributing to intraluminal obliteration through their expansion ^2^. In addition to the accumulation of PA resident cells, the increased deposition of ECM-related proteins, such as collagens, tenascin-C, osteopontin, and fibronectin, is a significant determinant of vascular remodeling^8–10^. While the importance of ECM molecules in structural changes within the PA wall is well established, relatively little attention has been paid to integrins, a superfamily of cell surface receptors that link the ECM to cell function^11^. The interaction of integrins with their extracellular ligands not only provides a physical anchor for the cell but also functions as a mechanosensor, transmitting ECM cues to the cell by triggering an array of intracellular pathways, including tyrosine kinases (TK) such as focal adhesion kinase (FAK), and the proto-oncogene tyrosine kinase c-Src, all of which are known to be critical to PAH ^1, 12^. Although multiple preclinical efforts have been pursued to directly block TKs in PAH, translating these efforts to the clinic remains a challenge. Integrins, by regulating virtually all basic cellular processes, including cell adhesion, migration, survival, and cell cycle progression^11, 13, 14^, in part through TK regulation, represent an attractive therapeutic targets for PAH that have never been explored.

Integrins are heterodimeric cell surface receptors composed of one α and one β subunit, which can associate in various combinations to generate twenty-four unique heterodimers. Depending on their ligand recognition pattern, integrins are classified into several groups. Among them, Arg-Gly-Asp (RGD)-binding integrins are of particular interest in the context of PAH, as several studies have reported their implication in smooth muscle cell phenotype switching in vascular proliferative diseases ^15–17^. Despite the biological importance of integrins and the scientific interest they have generated over the past years, pinpointing the exact molecular mechanisms in which each integrin is involved remains a major challenge. This is partly due to the limited availability of specific RGD integrin-related tools (inhibitors/activators) for preclinical validation and the inherent complexity of integrin functions. With recent clinical successes in the development of several integrin inhibitors (small molecules or antibodies) for conditions such as dry eye, gastrointestinal inflammation, and multiple sclerosis^18^, research aimed at targeting RGD integrins has become a priority. Nonetheless, their potential role in treating cardiovascular diseases like PAH remains underexplored.

In this study, we explored a panel of small molecule and antibody-based integrin inhibitors. Our findings provide compelling evidence that blocking α5β1, which is highly expressed in PASMCs and PAECs from PAH patients, interferes with the behavioral facets of diseased cells. Furthermore, α5β1 improves cardiopulmonary structure and function in multiple animal and human preclinical models. Taken together, these results highlight the potential utility of α5β1 inhibitors for the management of pulmonary hypertensive diseases.

## RESULTS

### Integrin subunits profiling in tissues and cells from human PAH patients and animal models

To begin exploring the potential role of integrins in the pathophysiology of the disease, we first measured the expression of the 18 α subunits and 8 β subunits, which can assemble into 24 different heterodimers (Figure 1A), by NanoString PlexSet assay, using RNA derived from PAH-PASMCs. As shown in Figure 1B, the β1 subunit, encoded by the ITGB1 gene, was the most abundant, followed by α5, β5, α2 and α6 (encoded by the ITGA5, ITGB5, ITGA2, ITGA6 genes, respectively). Since several of these integrins recognize an arginine-glycine-aspartate (RGD) tripeptide sequence present in a variety of ECM components implicated in maladaptive remodeling in PAH^9, 10^, we next compared the expression of some RGD-binding integrin subunits in dissected PA (<1 mm in diameter) from control and PAH patients. Except for β3, all subunits examined (α5, αV, β1, and β5) were significantly upregulated in PAH patients (Figure 1C). In isolated PASMCs cultured on fibronectin-coated plates, only α5 was significantly increased in diseased cells. In contrast, β5 expression was significantly decreased (Figure 1D). In isolated PAECs, both α5 and β1 were significantly upregulated in diseased cells (Figure 1E), making elevated α5 expression a common feature of PA resident cells in the setting of PAH. Accordingly, the expression levels of α5 and αV integrin subunits were upregulated in remodeled dissected PA from rats exposed to Sugen/Hypoxia (Su/Hx) or monocrotaline (MCT) (Figure S1). Consistently, increased expression of the α5β1 heterodimer was found in PAH-PASMCs compared to control cells, as assessed by Meso Scale Discovery electrochemiluminescence assay (Figure 1F). Taken together, these findings suggest that α5β1 may have a critical role in the development and progression of PAH.

**Figure 1.**
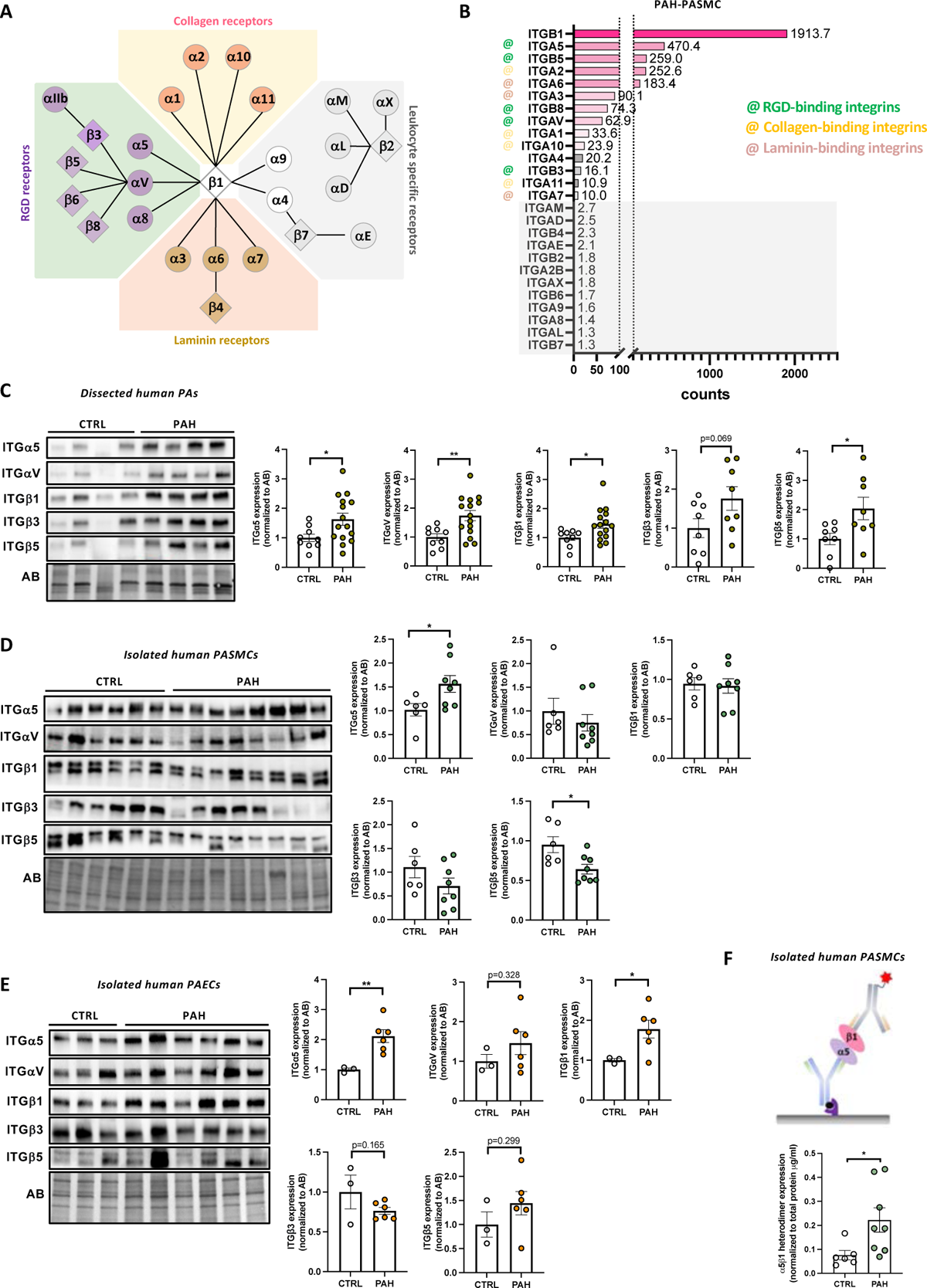
Integrin a5 subunit is upregulated in PASMCs and PAECs of PAH patients. (**A**) Organization and grouping of the integrin subunits in mammalian cells based on their matrix affinity. (**B**) Graph representing relative NanoString counts of various integrin subunits in PAH-PASMCs. (**C**) Representative Western blots and corresponding quantification of ITGa5, ITGaV, ITGpi, ITGP3, and ITGB5 in dissected pulmonary arteries from healthy donors and patients with PAH. Protein expression was normalized to amido black (AB) (n=8 or 15; *p<0.05, **p<0.01, unpaired Student’s t-test; data represent mean ± SEM). (**D**) Representative Western blots and corresponding quantification of ITGa5, ITGaV, ITGp1, ITGP3, and ITGP5 in PASMCs isolated from healthy donors and patients with PAH. Protein expression was normalized to amido black (AB) (n=6 or 8; *p<0.05, **p<0.01, unpaired Student’s t test for ITGa5, ITGpi, ITGP3 and ITGP5 expression or Mann-Whitney for ITGaV expression; data represent mean ± SEM). (**E**) Representative Western blots and corresponding quantifications of ITGa5, ITGaV, ITGp1, ITGP3, and ITGP5 in PAECs isolated from healthy donors and patients with PAH. Protein expression was normalized to amido black (AB) (n=3 or 6; *p<0.05, **p<0.01, unpaired Student’s t-test; data represent mean ± SEM). (**F**) a5pi protein levels as determined by Meso Scale Discovery custom electrochemiluminescence assay in control and PAH-PASMCs (n=6 or 8; *p<0.05, unpaired Student’s t test; data represent mean ± SEM).

### RGD-integrin inhibitor design and demonstration of potency and specificity

A drug discovery effort was undertaken to identify small molecule inhibitors of α5β1. We identified a small molecule RGD tripeptide mimetic, MRT1 (Figure 2A), which binds with high affinity to α5β1 in a fluorescence polarization (FP) assay (half-maximal effective concentration, IC_50,_ = 40 nM) and other RGD-binding β1 integrins such as αvβ1 and α8β1 (αvβ1 FP IC_50_ = 18 nM; α8β1 FP IC_50_ = 54 nM), and inhibits integrin activity in cell-based ligand binding assays with similar IC_50_ values (Figure 2C and Table S7). MRT1 is inactive against members of the collagen- and laminin-binding integrin subfamilies (Figure 2C and Table S7). The potency and selectivity of MRT1 combined with high cellular permeability in Madin-Darby canine kidney cells transfected with human MDR1 gene (MDCK-MDR1) (MDCK P_app_ = 7.86*10^-6^ cm/s) and the pharmacokinetic profile in rats (Table S8) supports the use of this novel reagent to examine α5β1/αvβ1 function in rat models of PAH^19^. Figure 2B shows the crystallographically determined binding pose of MRT1 with human α5β1. The tetrahydro-1,8-naphthyridine (THN) moiety of MRT1 makes critical salt bridge interactions with D227 of the α5-subunit, while the carboxyl group forms a strong metal coordination with the magnesium ion of the metal-ion-dependent adhesion site (MIDAS) in the β1-domain. MRT1 stabilizes the closed headpiece conformation by forming a hydrogen bond with a bridging water molecule (black arrow, Figure 2B)^20^. Additional potency and selectivity are conferred by interactions with the specificity-determining loop 2 (SDL2) ^21^.

**Figure 2.**
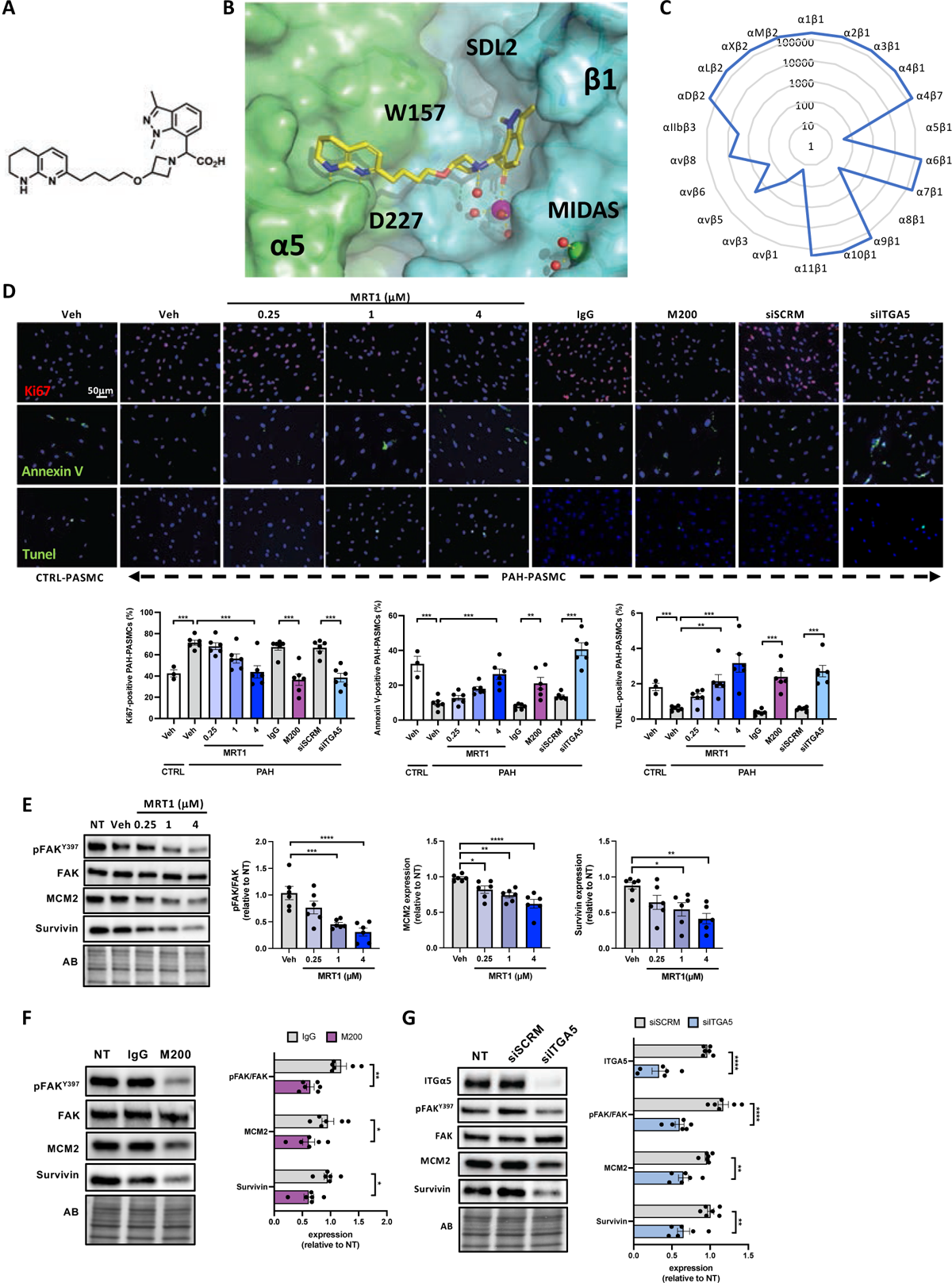
Inhibition of integrin a5pi reduces the pro-proliferative and apoptosis-resistant phenotype of PAH-PASMCs. (**A**) Structure of MRT1. (**B**) X-ray crystallographic structure of human a5pi headpiece in complex with MRT1 (yellow) determined at 1.7 A. The a5 subunit is shown in green and pi in cyan surfaces. The Mg^2+^ion is shown as a magenta sphere and water molecules are shown as red spheres. Hydrogen bonds with the ligand and coordination bonds between bound waters and the ligand are depicted as dashed lines; MIDAS: magnesium ion of the metal-ion-dependent adhesion site. (**C**) Radar plot of the inhibitory activity (in nM) of MRT1 against various integrins as determined through fluorescence polarization IC50 assays. (**D**) Representative fluorescent images of Ki67-labeled (red), Annexin V-labeled (green), and TUNEL-labeled (green) control and PAH-PASMCs exposed or not to escalating doses of MRT1, M200 (0.5 pg/ml) or siITGA5 (10 nM) for 48 hours. The corresponding quantifications are shown (n=3 or 6; *p<0.05, ***p<0.001, ****p<0.0001, one­way ANOVA followed by Tukey’s post hoc analysis; data represent mean ± SEM). Scale bars, 50 pm. (**E**) Representative Western blots, and corresponding quantifications of the ratio of phospho-FAK (Y397) to FAK, MCM2, and Survivin in PAH-PASMCs exposed or not to escalating concentrations of MRT1 for 48 hours. Protein expression was normalized to amido black (AB) (n=6; *p<0.05, **p<0.01, ***p<0.001, ****p<0.0001, one-way ANOVA followed by Dunnett’s post hoc analysis; data represent mean ± SEM). (**F**) Representative Western blots and corresponding quantifications of the ratio of phospho-FAK (Y397) to FAK, MCM2, and Survivin in PAH-PASMCs exposed or not to M200 (0.5pg/ml) for 48 hours (n=6; *p<0.05, **p<0.01, unpaired Student’s t test for MCM2 and Survivin expression and Mann-Whitney test for phospho-FAK (Y397) to FAK expression; data represent mean ± SEM). (**G**) Representative Western blots and corresponding quantifications of the ratio of phospho-FAK (Y397) to FAK, MCM2, and Survivin in PAH-PASMCs exposed or not to siITGA5 for 48 hours (n=6; **p<0.01, ****p<0.0001 unpaired Student’s t test; data represent mean ± SEM).

### MRT1 dose-dependently reduces PAH-PASMC and PAH-PAEC proliferation, resistance to apoptosis and migration without affecting healthy cells

We next investigated the effect of MRT1 on the abnormal behavior of PAH-PASMCs and PAH-PAECs. To this end, cells were plated on fibronectin and then exposed to escalating concentrations of MRT1 for 48 hours. As assessed by Ki67, Annexin V and TUNEL labeling, diseased cells were significantly more proliferative and resistant to apoptosis compared to their normal counterparts (Figure 2D and S2A). MRT1 treatment dose-dependently reduced the apoptosis resistance and proliferative capacity of PAH-PASMCs and PAH-PAECs, as evidenced by Annexin V, TUNEL and Ki67 labeling, respectively (Figure 2D and S2A). Western blotting yielded consistent results with all pro-proliferative and pro-survival markers tested (MCM2 and Survivin) being significantly diminished by MRT1 exposure (Figure 2E and S2B). Furthermore, using a scratch assay to evaluate the migratory ability of PAH-PASMCs, we found that MRT1 inhibition reduces the motility of diseased cells (Figure S3A). As expected, these effects were associated with a dose-dependent reduction in FAK phosphorylation (Y397) (Figure 2E and S2B). Note that MRT1 had no significant effect on control PASMC (Figure S3B).

### MRT1 effects on PAH-PASMC/PAH-PAEC proliferation, resistance to apoptosis and migration are mediated through the inhibition of α5β1

To determine the relative importance of each integrin heterodimer known to be modulated by MRT1 (α5β1, αvβ1 and α8β1) in PAH, we first exposed PAH-PASMCs to escalating doses of MRT6; a selective αvβ1 and α8β1 (IC_50_= 13.2 nM and IC_50_= 30.8 nM) integrin inhibitor (Figure S4A). As determined by Ki67 labeling and Western blots for proliferation markers (MCM2 and PCNA), the proliferative capacity of PAH-PASMCs was not affected by MRT6 (Figure S4, A and B). No change in the percentage of Annexin V-positive cells and surviving expression was observed after MRT6 exposure (Figure S4A). In addition, the abnormal phenotype of PAH-PAECs was not affected by MRT6 treatment (Figure S4C). We therefore hypothesized that the beneficial effects of MRT1 *in vitro* are mainly mediated by its inhibitory activity of α5β1 binding to fibronectin. This is supported by published data showing that cells expressing α5β1 integrin are more resistant to apoptosis than the same cells engineered to express αvβ1^22^.

To further validate the role of α5β1 in PAH-PASMC and PAH-PAEC proliferation and apoptosis, cells were exposed to M200 (volociximab), a clinically tested, high-affinity, blocking antibody against human α5β1^23^, Figure 2 and S2). Assessment of cell proliferation and apoptosis by Ki67, Annexin V and TUNEL labeling showed that blockade of α5β1 integrin significantly reduced proliferation and survival of PAH-PASMCs and PAH-PAECs (Figure 2D and S2C). These findings were substantiated by immunoblot analysis for proliferation/survival markers (Figure 2F and S2D). To complement our findings, a molecular approach was performed by depleting the α5 subunit using siRNA. Compared to control cells transfected with a scrambled siRNA, cells receiving the ITGA5-targeting siRNA exhibited >60% reduction in α5 protein levels (Figure 2G and S2F), which was accompanied by a significant reduction in disease cell proliferation and resistance to apoptosis (Figure S2D and S2E). Exposure to M200 and knockdown of ITGA5 also resulted in reduced motility of PAH-PASMCs (Figure S3A). As seen for MRT1, these effects were associated with a reduction in the activation of FAK network (Figure S2, D and F).

### Inhibition of α5β1 integrin decreases a FOXM1-dependent pro-mitotic network

To gain mechanistic insights into how α5β1 integrin blockade elicits therapeutic effects, we performed RNA-Seq transcriptomic analysis on human PAH-PASMCs exposed to MRT1 or M200 for 48 hours. Using a cut-off for |fold change| >1.4 and an adjusted p-value < 0.05, we identified 194 and 216 genes that were differentially expressed (DEG) in MRT1 *vs.* vehicle and M200 *vs.* vehicle, respectively (Figure 3A). We next carried out functional annotation analysis using ShinyGO to investigate the biological relevance of downregulated genes. Consistent with our *in vitro* and *in vivo* observations, we found that downregulated genes upon MRT1 or M200 exposure were involved in the regulation of chromosome segregation/mitotic spindle organization (Figure 3B). Gene set enrichment analysis (GSEA) revealed that among the genes downregulated by MRT1 and M200, those regulating G2/M checkpoint and mitotic spindle were significantly enriched (Figure 3C). These include budding uninhibited by benzimidazole 1 (BUB1), NIMA-related kinase 2 (NEK2), and Centromere Protein M (CENPM). By combining these two analyses, we generated a high-confidence list of 87 genes that were consistently altered in both comparisons and exhibited the same direction of change (44 upregulated and 43 downregulated) (Figure 3, D and E). To support our transcriptomic findings highlighting a requirement of α5β1 for proper progression through mitosis, we enriched PAH-PASMCs in G2/M and treated them with α5β1 inhibitors. MRT1- or siITGA5-treated cells exhibited centrosome abnormalities, including failure of centrosome separation and centrosome fragmentation, resulting in mono- and multipolar mitotic spindles (Figure 3F). The experiment did not include M200-treated cells to mitigate potential challenges associated with high background and non-specific labeling. This precaution was taken to avoid any potential cross-reactivity issues between secondary antibodies and the M200 antibody.

**Figure 3.**
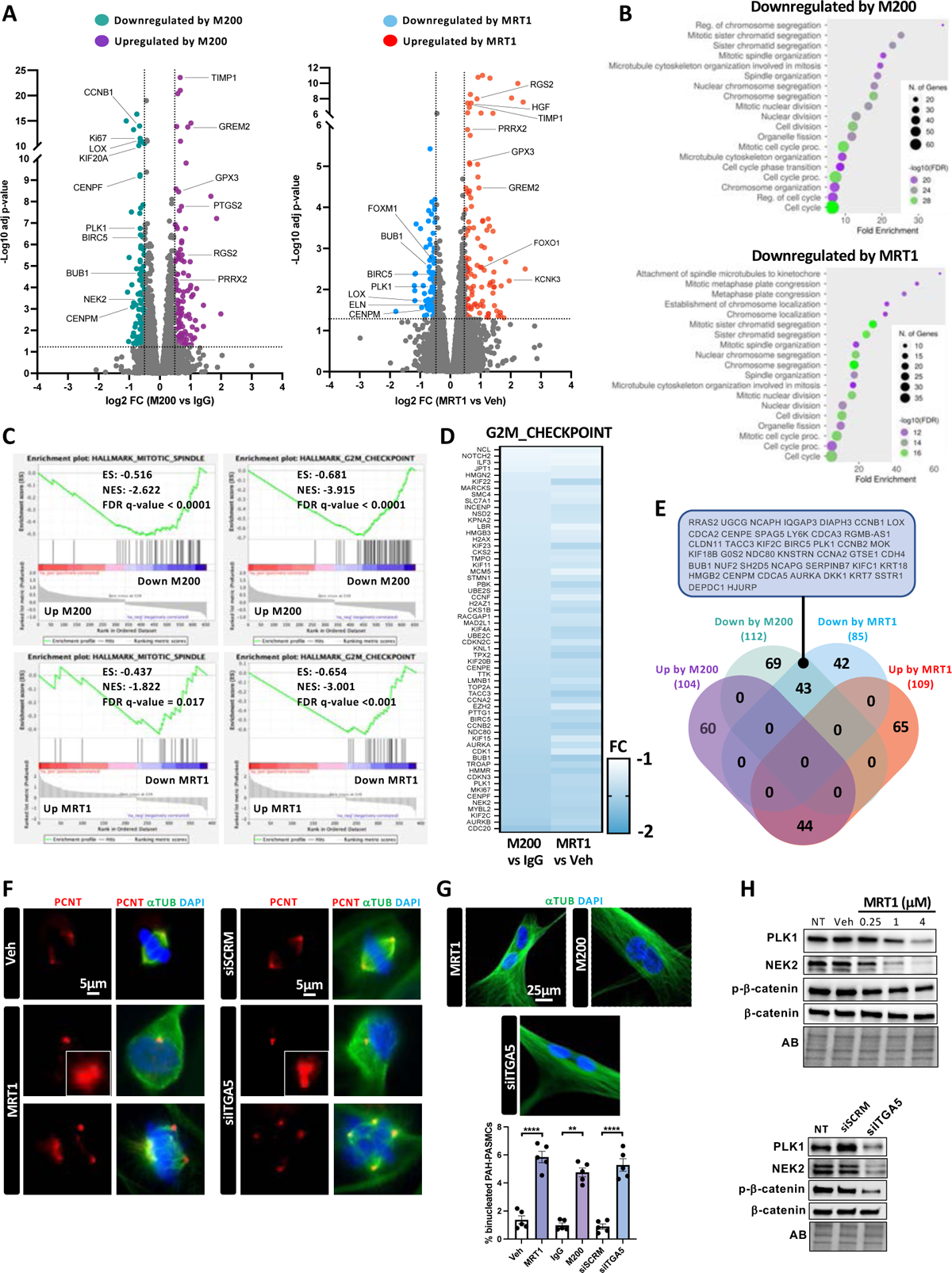
Integrin a501 regulates a mitotic gene network. (**A**) Volcano plots of differentially expressed genes in response to M200 or MRT1 treatment for 48 hours across different PAH-PASMC cell lines (n=4), with genes considered as differentially expressed (|FC| > 1.4, adjusted p-value <0.05) colored in purple/red (up-regulated by a5p1 inhibition) or turquoise/blue (down-regulated by a5p1 inhibition). Genes of interest are highlighted. (**B**) Enrichment analysis performed by ShinyGo software v0.77 showing the top GO biological processes (BP) in both down- and up-regulated genes following M200 or MRT1 exposure in PAH-PASMCs. (**C**) Gene set enrichment analysis (GSEA) charts showing the enrichment of genes related to mitotic spindle and G2/M checkpoint. A negative Normalized Enrichment Score (NES) value indicates the downregulation in a5p1-inhibited PAH-PASMCs. (**D**) Heatmap displays genes related to G2/M checkpoint that are altered (adjusted P<0.05) by M200. The fold change of the same genes in MRT1-treated PAH-PASMCs is shown. (**E**) Venn diagram of shared up- and down-regulated transcripts between the indicated groups. (**F**) Representative images of the structure of the mitotic spindle in synchronized PAH-PASMCs treated or not with MRT1 (4 pM, 10 hours) or siITGA5 (transfected 24 hours prior to the double thymidine block initiation). Red, green and blue in the image represent pericentrin, a-tubulin and DNA, respectively. Scale bar: 5 pm. (**G**) Representative images and frequency of binucleated cells after treatment with MRT1 (4 pM, 10 hours), M200 (0,5 pg/ml, 10 hours) or siITGA5 (transfected 24 hours prior to the double thymidine block initiation). (n=5; ****p<0.0001, unpaired Student’s t test for MRT1 and siITGA5 and Mann-Whitney test for M200; data represent mean ± SEM). Scale bar: 25 pm. (**H**) Representative Western blots of PLK1, NEK2, phospho-P-catenin and P-catenin in PAH-PASMCs exposed or not to escalating doses of MRT1 for 48 hours or siITGA5 for 48 hours.

Consistently, decreased expression of the mitotic marker pHH3 and increased binucleation were observed in α5β1-inhibited PAH-PASMCs (Figures S5A and 3G). Furthermore, expression levels of Polo-like kinase 1 (PLK1), NIMA-related kinase 2 (NEK2), and phosphorylation of its downstream target β-catenin; a mitotic cascade documented to regulate centrosome duplication and separation^24^ and increased in PAH-PASMCs (Figure S5B, ^25–27^), were found to be reduced in cells exposed to either MRT1, M200 or siITGA5 (Figures 3H and S5, C and D). Similar findings were observed in PAH-PAECs subjected to MRT1, M200 or siITGA5 treatments (Figure S6). This supports the view that α5β1 controls centrosome disjunction/splitting and the execution of mitosis.

To further dismantle the mechanism of action of α5β1 we performed an EnrichR predictive analysis of the 43 genes commonly downregulated by MRT1 and M200 and identified FOXM1, which is increased in PAH-PASMCs, as the first transcription factor governing the expression of these genes (Figure 4A and B). This finding was confirmed at the mRNA levels by treating PAH-PASMCs with FOXM1 siRNA. We showed that 31 genes among the 43 downregulated by MRT1/M200 were also significantly decreased upon FOXM1 inhibition (Figure 4C). At the protein level, we demonstrated that MRT1, M200 and siITGA5 significantly repress FOXM1 expression (Figure 4, D to F). Finally, the mechanism accounting for FOXM1 activation by α5β1 is mediated, in part, by FAK activation as FAK inhibition by PF-573228 significantly represses FOXM1 expression (Figure 4G). This finding is in line with previously reported results in cancer studies^28, 29^.

**Figure 4.**
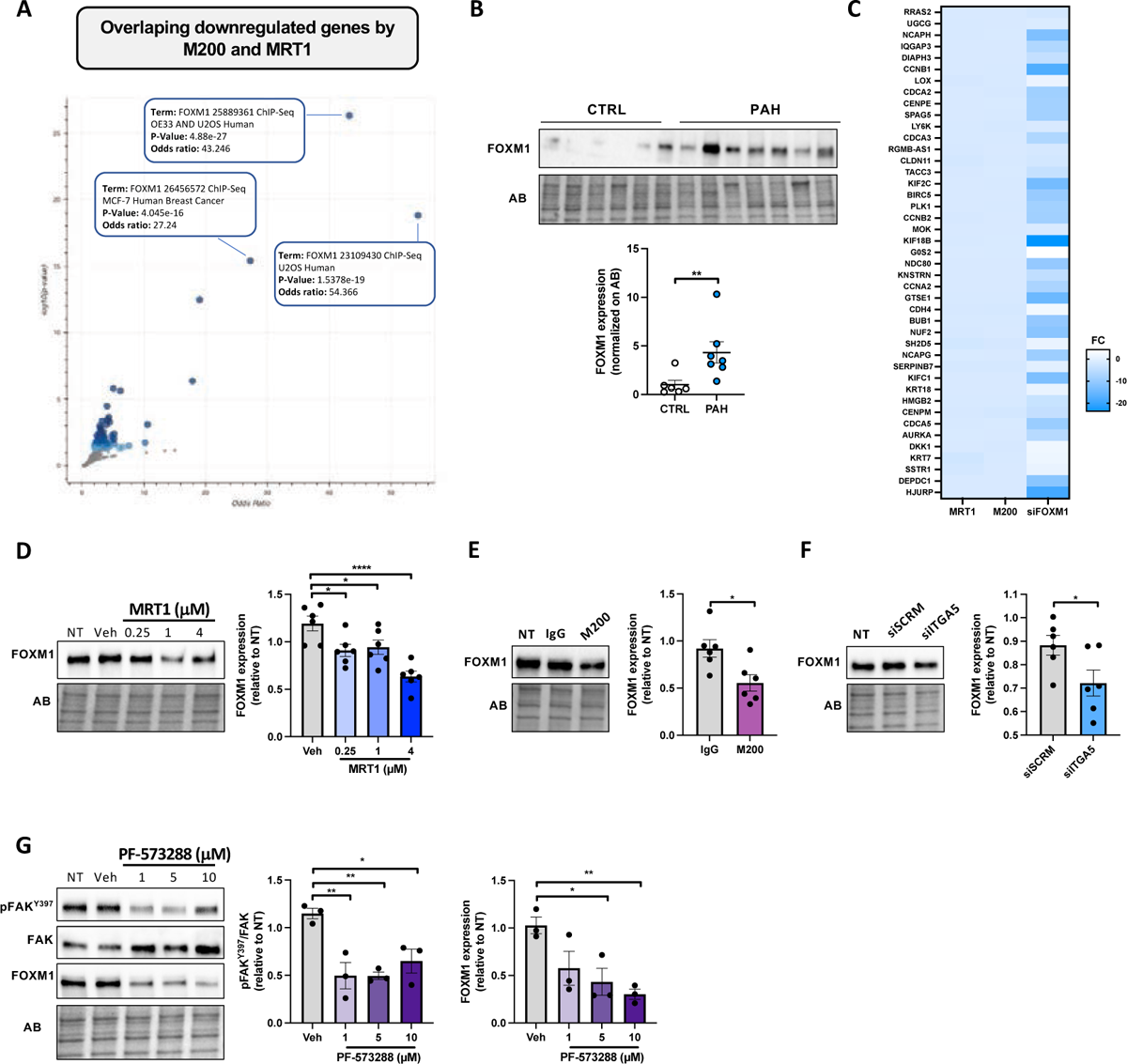
The downregulated genes upon a501 inhibition are regulated by FOXM1. (**A**) Volcano plot illustrating terms from the encode and ChEA consensus transcription factors from the ChIP-X gene set. Each term is represented as a point, positioned according to its odds ratio (position on the x-axis) and -log10 (p-value) (position on the y-axis), reflecting results from the enrichment analysis of the gene set submitted. Stronger enrichment significance of the term within the input gene set is represented by larger and darker-colored point. (**B**) Western blot and corresponding quantification of FOXM1 in PASMCs isolated from healthy donors and patients with PAH. Protein expression was normalized to amido black (AB) (n=6 or 7; **p<0.01, unpaired Student’s t-test; data represent mean ± SEM). (**C**) Heatmap displaying the overlapping downregulated genes by M200 and MRT1. The fold change of the same genes in siFOXM1-treated PAH-PASMCs is shown. (**D**) Representative Western blot, and corresponding quantification of FOXM1 in PAH-PASMCs exposed or not to escalating concentrations of MRT1 for 48 hours. Protein expression was normalized to amido black (AB) (n=6; *p<0.05, ****p<0.0001, one-way ANOVA followed by Dunnett’s post hoc analysis; data represent mean ± SEM). (**E**) Representative Western blot, and corresponding quantification of FOXM1 in PAH-PASMCs exposed or not to M200 (0.5pg/ml) for 48 hours (n=6; *p<0.05, unpaired Student’s t-test; data represent mean ± SEM). (**F**) Representative Western blot, and corresponding quantification of FOXM1 in PAH-PASMCs exposed or not to siITGA5 for 48 hours (n=6; *p<0.05 unpaired Student’s t-test data represent mean ± SEM). (**G**) Representative Western blot, and corresponding quantification of the ratio of phospho-FAK (Y397) to FAK and FOXM1 in PAH-PASMCs exposed or not to PF-573288 at indicated doses for 48 hours (n=3; *p<0.05, **p<0.01; one­way ANOVA followed by Dunnett’s post hoc analysis; data represent mean ± SEM).

### MRT1 reverses pulmonary vascular remodeling and improves hemodynamics *in vivo*

Having demonstrated that MRT1 corrects the pro-proliferative and apoptosis-resistant phenotype of PAH-PASMCs and PAH-PAECs *in vitro*, using PAH clinical trials principle (randomization, blinding’s comparison to standard of care (macitentan/tadalafil) and in combination to standard of care) we next investigated the therapeutic effect of MRT1 in the multiple PAH rodents models, which is known to develop progressive obliterative and complex lesions reminiscent of human PAH^19^.

As described in Figure 5A, adult male rats were injected with SU5416 (Sugen, Su) and then subjected to chronic hypoxia for three weeks. Prior to initiation of MRT1 treatment on Day 21, animals were evaluated by transthoracic echocardiography (echo) to confirm the presence of the disease in all animals subjected to Su/Hx (Figure S7A). Once PAH was established after 3 weeks of hypoxia, the animals were returned to normoxic conditions and randomly assigned to four groups. Rats were treated orally by gavage with either vehicle, macitentan+tadalafil (PAH standard-of-care drugs that function mainly as vasodilators), MRT1 alone, or MRT1+macitentan+tadalafil for an additional two weeks. A control group (no PAH) was also monitored. As assessed by echo, MRT1 treatment improved conventional parameters of RV function (cardiac output (CO), stroke volume (SV), tricuspid annular plane systolic excursion (TAPSE), S-wave, and RV fractional area change (FAC)) when compared to vehicle-treated Su/Hx animals (Figures 5B and S7B). The improvement in hemodynamics and RV function was also confirmed by closed-chest right heart catheterization (RHC) with a significant lowering of RV systolic pressure (RVSP) and mean PA pressure (mPAP) associated with an improvement in SV and CO and a reduction in total pulmonary vascular resistance (TPR) (Figure 5C). Medial wall thickness of small PAs (<75 μm in diameter) was next quantified. As predicted by the hemodynamic data, treatment with MRT1 resulted in a significant reduction in vascular remodeling (Figure 5D). Consistent with this, fewer proliferating (PCNA-positive) and more apoptotic (cleaved caspase 3-positive) PASMCs were detected in MRT1-treated Su/Hx animals (Figure 5D). Immunofluorescence analysis revealed that phosphorylated FAK levels were significantly reduced in distal PAs of MRT1-treated Su/Hx rats compared to vehicle-treated Su/Hx animals, confirming the efficacy of MRT1 to inhibit RGD-integrins function *in vivo* (Figure S7C). Beyond its beneficial effects on the pulmonary vasculature, inhibition of RGD-integrins significantly attenuated Su/Hx-induced RV mass, cardiomyocyte (CM) hypertrophy and RV fibrosis, as assessed by the Fulton index, quantification of CMs cross-sectional area (CSA), and Masson’s trichrome staining, respectively (Figure S7, D and E). Moreover, we found that MRT1 had similar therapeutic effects to a combination of macitentan and tadalafil, and a superior effect when combined with macitentan and tadalafil (Figure 5, B to D). The circulating concentration of MRT1 in the plasma was measured at sacrifice (results presented in Table S9) confirming expected target coverage. Importantly, MRT1 treatment was well tolerated with no treatment-related adverse findings on body weight, systemic pressures, or changes in plasma levels of kidney and liver injury markers (Figure S8). The effects of MRT1 on hemodynamic and histological endpoints in the Sugen/hypoxia model were independently confirmed by a contract research organization (Supplemental Figure S9).

**Figure 5.**
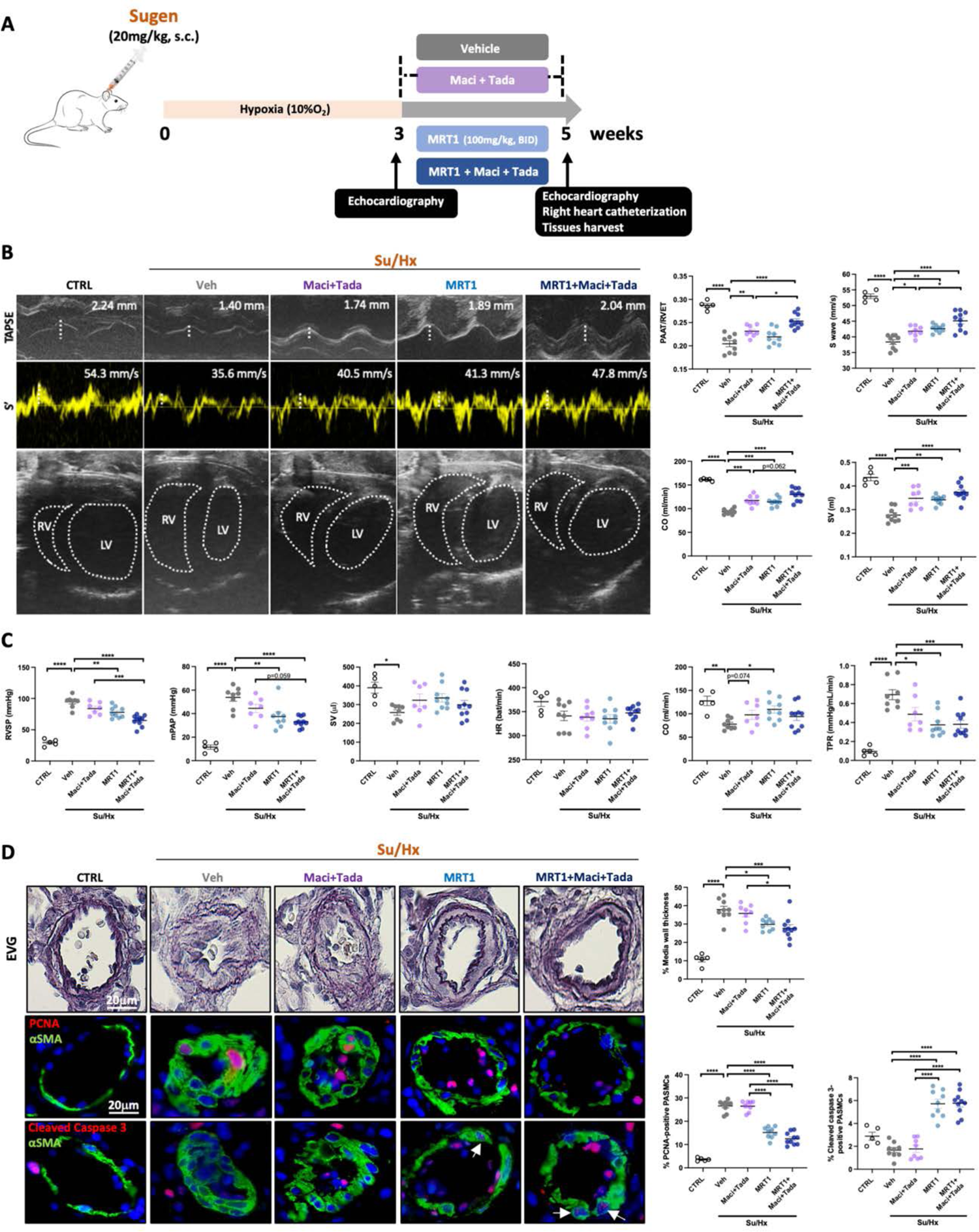
Treatment with RGD-integrins inhibitor alone or combined with macitentan and tadalafil attenuates established PAH in Su/Hx rats. (**A**) Study design using the Sugen/Hypoxia (Su/Hx) rat model. (**B**) Representative echocardiographic images of tricuspid annular plane systolic excursion (TAPSE), S wave (S’), and morphological change of the RV and LV in control and Su/Hx rats treated with vehicle, macitentan+tadalafil, MRT1 or MRT1+macientant+tadalafil. The quantifications of PAAT/ET, S wave, CO, and SV are shown (n=5 to 10; *p<0.05, **p<0.01, ***p<0.001, ****p<0.0001, one-way ANOVA followed by Tukey’s post hoc analysis; data represent mean ± SEM). (**C**) Effect of integrin a5p1/avp1 inhibition on RVSP, mPAP, SV, HR, CO and TPR, as assessed by right heart catheterization (n=5 to 10; *p<0.05, **p<0.01, ***p<0.001, ****p<0.0001, one-way ANOVA followed by Tukey’s post hoc analysis for RVSP, mPAP, SV, HR, CO and Kruskal-Wallis’ test followed by Dunn’s post hoc analysis for TPR; data represent mean ± SEM). (**D**) Representative images of distal PAs stained with Elastica van Gieson (EVG) or labeled with PCNA (proliferative marker, red) or cleaved caspase 3 (apoptosis marker, red). Alpha smooth muscle actin (aSMA, green) was used to detect smooth muscle cells. The quantifications of medial wall thickness, PCNA-positive and cleaved caspase 3-positive smooth muscle cells are presented on the right. (n=5 to 10; *p<0.05, **p<0.01, ***p<0.001, ****p<0.0001, one-way ANOVA followed by Tukey’s post hoc analysis; data represent mean ± SEM). Scale bars, 20 pm.

Because no animal model fully recapitulates the features of human disease^19^ and in accordance with recent recommendations for optimal preclinical studies in PAH^30^, we decided to test the therapeutic effects of MRT1 in a second drug-induced animal model, namely, the monocrotaline (MCT) rat model (Figure S10A). Hemodynamic parameters before treatment are presented in Supplementary Figure S11A. As shown by echo and RHC, we found that treatment with either MRT1 alone or MRT1+macitentan+tadalafil significantly improved pulmonary hemodynamics (RVSP and mPAP) and RV function (TAPSE, S wave, CO, SV, RVFAC) (Figure S10, B and C and Figure S11B). Medial wall thickness of small PAs, FAK phosphorylation, Fulton index, CM hypertrophy and RV fibrosis were also significantly reduced by MRT1 administration (Figure S10D and Figure S11, C to F). Taken together, our results indicate that RGD-integrins drive pulmonary and RV remodeling processes in the setting of PAH. Finally, to confirm the anti-remodeling effect of MRT1, we compared its efficacy to imatinib an accepted anti-remodeling agent that has shown its efficacy in multiple PAH preclinical models^31, 32^. MRT1 has similar hemodynamic and anti-remodeling effects than imatinib as shown in Figure S12.

### Antibody blockade of α5β1 ameliorated PAH in rodents

To further support the fact that MRT1 effects are mediated by α5β1 inhibition, we demonstrated that administration of MRT6 failed to improve Fulton index, pulmonary hemodynamics, and RV function in rats (Figure S13), further refuting the implication of αvβ1/α8β1 in PAH physiopathology. We next generated a rat blocking antibody against rat α5β1. The mAb4 clone (sequence shown in Figure 6A) proved to be a highly potent blocker, exhibiting strong affinity for rat α5β1 (dissociation constant, K_d_ = 11.5 nM by surface plasmon resonance, IC_50_ = 3.75 nM in cell-based assay and IC_50_ = 6.3 nM in solid-phase assay, Table S10) and an acceptable pharmacokinetic profile in rats (Table S11). To confirm that our generated rat antibody has similar properties to the human (M200) and mice (339.1) commercially available α5β1 antibody, we performed negative-stain electron microscopy that showed mAb4 binds the β-propeller domain of α5, as the purified integrin ectodomain assumed its native closed conformation (Figure 6A)^33^. Next, we evaluated its effects in the Su/Hx rat model (Figure 6B). Echo parameters recorded before the start of treatment are presented in Supplemental Figure S14A. Compared with the IgG isotype control-treated Su/Hx rats, RV hypertrophy was significantly reduced when rats were treated with mAb4 (Figure 6C). mAb4 administration significantly increased PAAT/RVET ratio, RVFAC, CO, SV, TAPSE and S wave when compared to IgG isotype control Su/Hx rats (Figure 6D and S14 B, C). Measurements by RHC showed that RVSP and mPAP were significantly decreased by mAb4 (Figure 6E). As a result, mAb4 significantly reduced TPR as compared to IgG isotype control rats (Figure 6E). Western blot and immunofluorescence analysis revealed that phospho-FAK expression levels were significantly reduced in mAb4-treated Su/Hx rats compared to IgG isotype control Su/Hx animals, confirming the efficacy of mAb4 to block α5β1 integrin engagement *in vivo* (Figures 6G and S14D). Morphometric analysis of small PAs showed that Su/Hx injection resulted in severe medial wall thickening, which was markedly reduced by mAb4 (Figure 6F). Consistent with this finding, the proportion of PCNA-positive proliferating PASMCs was reduced by mAb4, which was associated with a concomitant increase in the proportion of apoptotic cells (Figure 6F). Histological and molecular analysis revealed that RV cardiomyocyte hypertrophy and fibrosis were reduced in the mAb4-treated Su/Hx group compared with the vehicle group (Figure S14, E and F). Reduced expression of *Col1α1*, *Col3α1*, *Ltbp2* (three pro-fibrotic markers) and *Nppb* (cardiac stress marker) was also observed in the RV of mAb4-treated rats compared to IgG isotype control animals (Figure 6H). As for MRT1, treatment with mAb4 did not result in any significant toxicity assessed by liver and kidney function, body weight and systemic pressure (Figure S8). Collectively, these results provide compelling evidence that targeting the α5β1 integrin reverses existing pulmonary hypertension and RV dysfunction.

**Figure 6.**
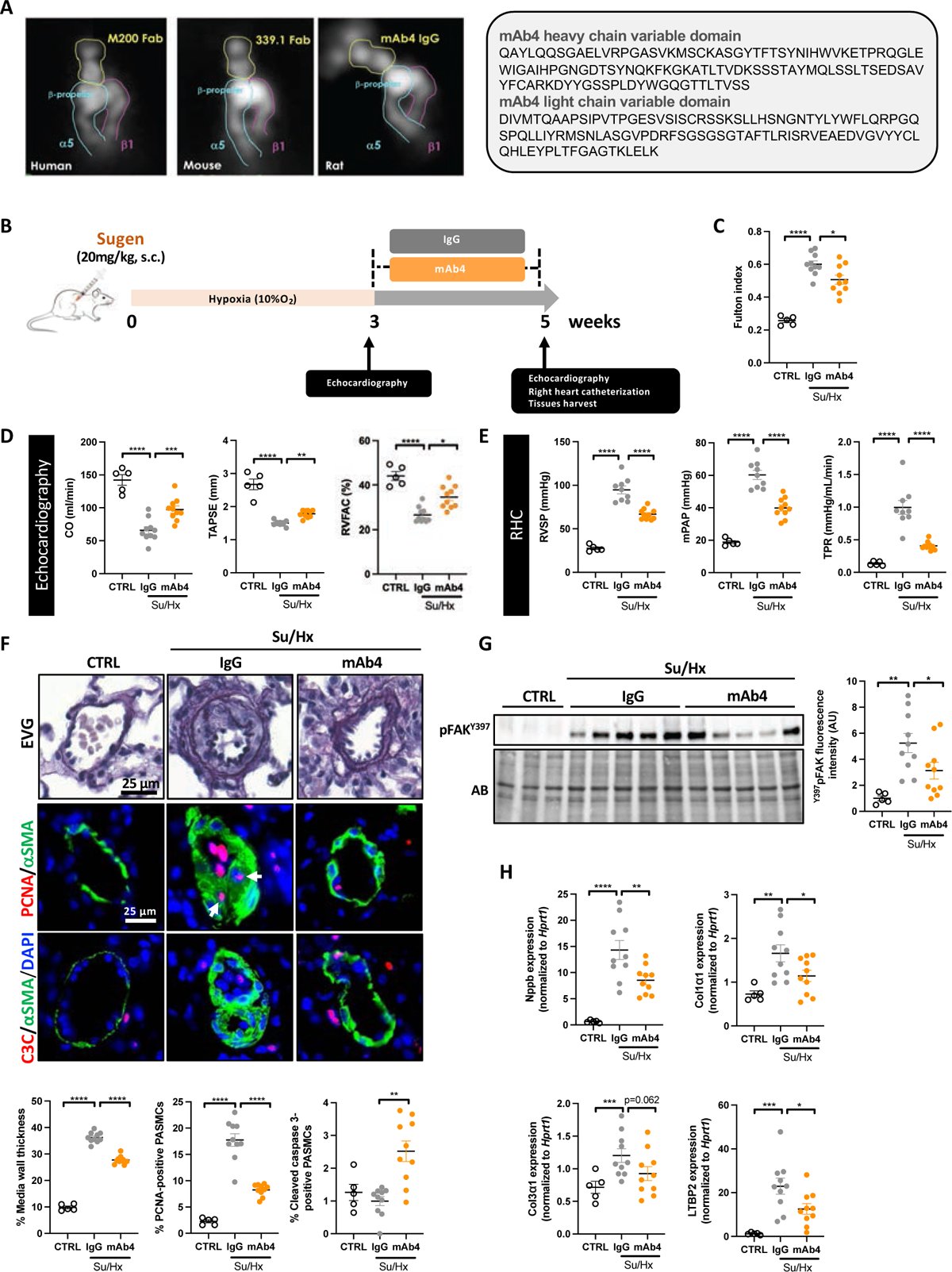
A function-blocking anti-rat integrin a5pi antibody improves established PAH in Su/Hx-challenged rats. (**A**) Representative 2D classes of the three a5-specific blocking antibodies used in the study, M200 (as Fab), 339.1 (Fab) and mAb4 (IgG) in complex with recombinant a501 ectodomain proteins of their corresponding species and variable domain sequences of heavy and light chains. The yellow lines highlight the Fab fragments, cyan for the a5 and magenta for 01 subunit. The complex structures clearly indicate that the functional epitope of all three antibodies is located in the P-propeller domain of the a5 subunit. The P-propeller domain has been shown to interact with the Fn9 domain of fibronectin ^61^ and hence explaining the blocking activity of these antibodies. (**B**) Study design using the Sugen/Hypoxia (Su/Hx) rat model. (**C**) Assessment of RV hypertrophy by Fulton index (n=5 to 10; *p<0.05, ****p<0.0001, one-way ANOVA followed by Dunnett’s post hoc analysis; data represent mean ± SEM). (**D**) CO, TAPSE, and RVFAC measured using echocardiography in control and Su/Hx rats treated with IgG or mAb4 (n=5 to 10; *p<0.05, **p<0.01, ***p<0.001, ****p<0.0001, one-way ANOVA followed by Dunnett’s; data represent mean ± SEM). (**E**) Effect of integrin a5pi inhibition on RVSP, mPAP, and TPR, as assessed by right heart catheterization (n=5 to 10; ****p<0.0001, one-way ANOVA followed by Dunnett’s post hoc analysis; data represent mean ± SEM). (**F**) Representative images and corresponding quantifications of distal PAs stained with Elastica van Gieson (EVG) or labeled with PCNA (proliferative marker, red) or cleaved caspase 3 (apoptosis marker, red) in control, Su/Hx+Veh and Su/Hx+mAb4 rats. PASMCs were labeled with alpha-smooth muscle actin (aSMA, green). (n=5 to 10; **p<0.01, ****p<0.0001, one-way ANOVA followed by Dunnett’s post hoc analysis for EVG and PCNA, Kruskal-Wallis’ test followed by Dunn’s post hoc analysis for cleaved caspase 3; data represent mean ± SEM). Scale bars, 25 gm. (**G**) Representative Western blot and corresponding densitometric analyses of phospho-FAK (Y397) in dissected PAs from control and Su/Hx rats treated with either mAb4 or IgG control. Protein expression was normalized to amido black (AB) (n=5 to 10; ***p<0.001, one-way ANOVA followed by Dunnett’s post hoc analysis; data represent mean ± SEM). (**H**) *Nppb*, *Col1a1*, *Col3a1* and *Ltbp2* transcripts in control, Su/Hx+Veh and Su/Hx+mAb4 rats (n=5 to 10; *p<0.05, **p<0.01, ***p<0.001, one-way ANOVA followed by Dunnett’s post hoc analysis; data represent mean ± SEM)

To further emphasize the importance of α5β1 in MRT1 response in PAH, all these findings were confirmed in Su/Hx mice using a commercially available and in vivo validated blocking anti-mouse α5β1 antibody called 339.1^34^ (Figure S15, Figure S16). 339.1 and MRT1 produced comparable efficacy on pulmonary hemodynamics, PAs vascular remodeling and RV hypertrophy and fibrosis. This indicates that the therapeutic efficacy of MRT1 is mediated through the inhibition of α5β1 (Figure S15, Figure S16).

### MRT1 directly improves right ventricular function in the pulmonary artery banding rat model

RV function is the predominant determinant of survival in patients with PAH^35^. Published data documenting the expression of RGD-binding integrins in the hypertrophied heart^36, 37^ together with the results presented above suggest that inhibition of RGD-integrins may have no adverse effect on RV remodeling or even direct cardioprotective effects. To test our hypothesis, we used a surgical rat model of pulmonary artery banding (PAB) to induce RV afterload/dysfunction without increasing pulmonary vascular resistance. Only male rats were used to reduce the experimental variability. Three weeks after PAB, rats were randomized to the vehicle or MRT1 groups for a treatment period of seven weeks (Figure 7A). As shown in Figure 7B, there were no significant differences in the degree of constriction between vehicle and MRT1-treated PAB rats noted at the end of the treatment period. Longitudinal echocardiographic assessments revealed a gradual decline in RV function in vehicle-treated PAB rats as compared to sham-operated rats, as indicated by a progressive decrease in the values of SV, CO, TAPSE, S-wave and RV FAC. In parallel, a change in the opposite direction was observed for the RV end-diastolic diameter (RVEDD). In contrast, treatment with MRT1 significantly improved all these parameters (Figure 7, C and D), which was supported by RHC measurement (Figure 7E). Inhibition of RGD-integrins also resulted in a significant reduction in CMs CSA and a marked tendency to decrease RV fibrosis (Figure 7F).

**Figure 7.**
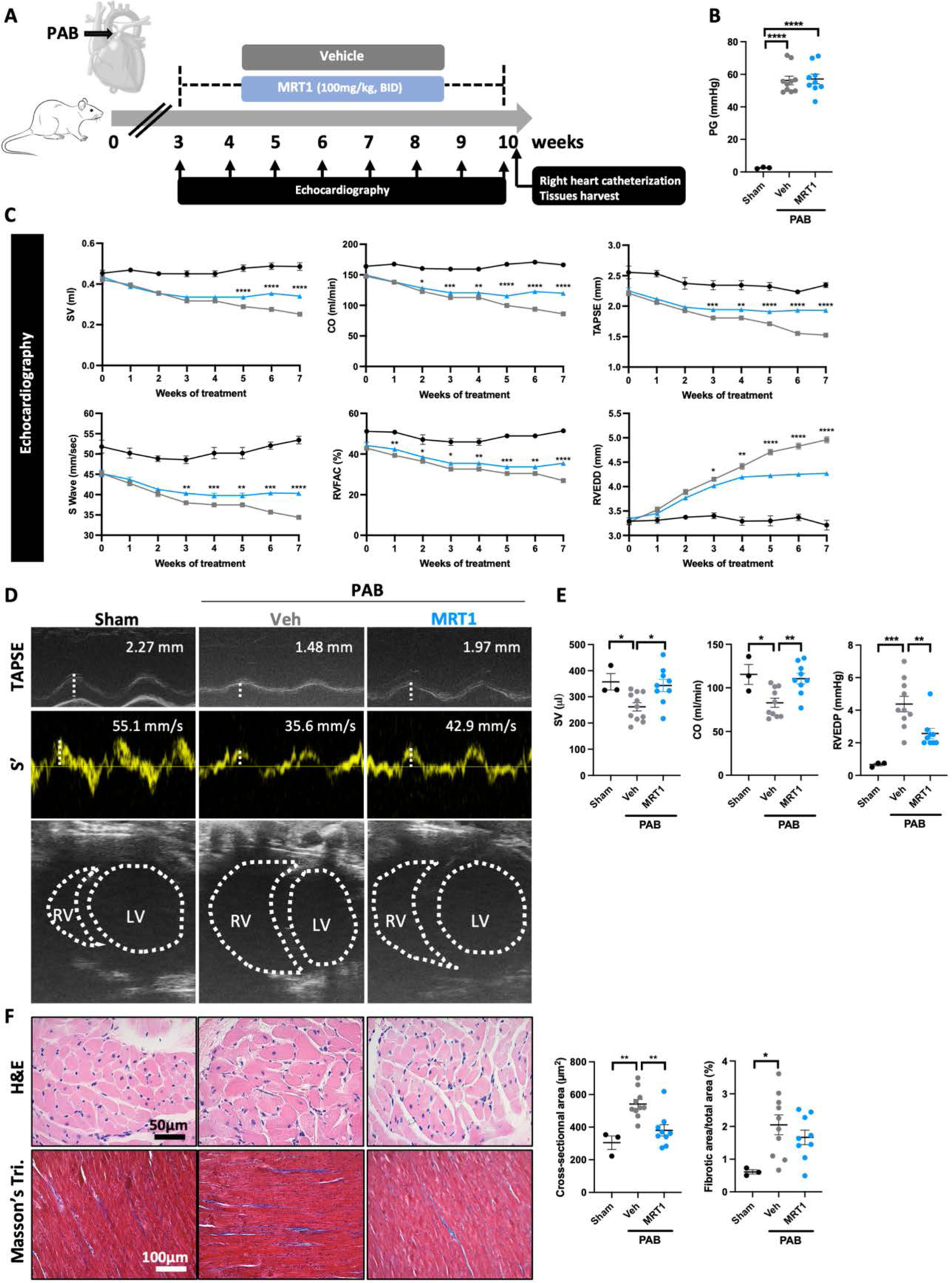
Inhibition of RGD-integrins elicits cardioprotective effects in rats subjected to PAB. (**A**) Schematic representation of the experimental protocol for induction and therapeutic intervention in a PAB-induced RV dysfunction model in rats. (**B**) Pressure gradient (PG) at PAB site measured by echocardiography at the end of the protocol (n=9 to 10; ****p<0.0001, Mann-Whitney test; data represent mean ± SEM). (**C**) Stroke volume (SV), cardiac output (CO), tricuspid annular plane systolic excursion (TAPSE), S wave, right ventricular fractional area change (RVFAC), and right ventricular end-diastolic diameter (RVEDD) measured by echocardiography in sham-operated and PAB-subjected rats every week after treatment or not with MRT1 (n=3 to 10; *p<0.05, **p<0.01, ***p<0.001, ****p<0.0001, linear mixed model; data represent mean ± SEM). (**D**) Representative echocardiographic images of TAPSE, S wave (S’) and morphological change of the RV and LV in control and PAB rats treated with MRT1 or vehicle at 7 weeks after the initiation of the treatment. (**E**) SV, CO, and RV end-diastolic pressure (RVEDP) measured by right heart catheterization at the protocol in control and PAB rats treated with MRT1 or vehicle (n=3-10/group) (n=3 to 10; *p<0.05, **p<0.01, ***p<0.001, one-way ANOVA followed by Dunnett’s post hoc analysis for CO and Kruskal-Wallis’ test followed by Dunn’s post hoc analysis for SV and RVEDP; data represent mean ± SEM). (**F**) Representative images of RV sections stained with Hematoxylin and eosin (H&E) or Masson’s Trichrome in control and PAB rats treated with MRT1 or its vehicle. The measurement of cardiomyocyte cross-sectional area (CSA) and the quantification of the percentage of collagen area are presented on the right. (n=3 to 10; *p<0.05, **p<0.01, one-way ANOVA followed by Dunnett’s post hoc analysis for H&E and Kruskal-Wallis’ test followed by Dunn’s post hoc analysis for Masson’s trichrome; data represent mean ± SEM). Scale bars: 50 μm (H&E) and 100 μm (Masson’s Trichrome).

### Inhibition of integrin α5β1 improves vascular remodeling in human ex vivo organotypic model precision-cut lung slices

On the strength of these *in vitro* and *in vivo* results, we investigated the therapeutic efficacy of α5β1 inhibitors *ex vivo* using human precision-cut lung slices (PCLS), a model that offers the advantage of preserving cell-cell and cell-matrix interactions. We first modeled the development of PH by exposing PCLS derived from control donors to a growth factor cocktail (GFC). Co-treatment with MRT1 or M200 mitigated GFC-induced medial wall thickness of small PAs and proliferation of PASMCs (Figure 8A). Pulmonary hypertension (PH) lungs were obtained from 5 patients (3 PAH and 2 idiopathic pulmonary fibrosis-associated PH). Treatment with α5β1 inhibitors significantly reduced medial wall thickness and PASMC proliferation (Figure 8B). Taken together, these data suggest that α5β1 inhibitors may represent new tools in our therapeutic armamentarium against PAH.

**Figure 8.**
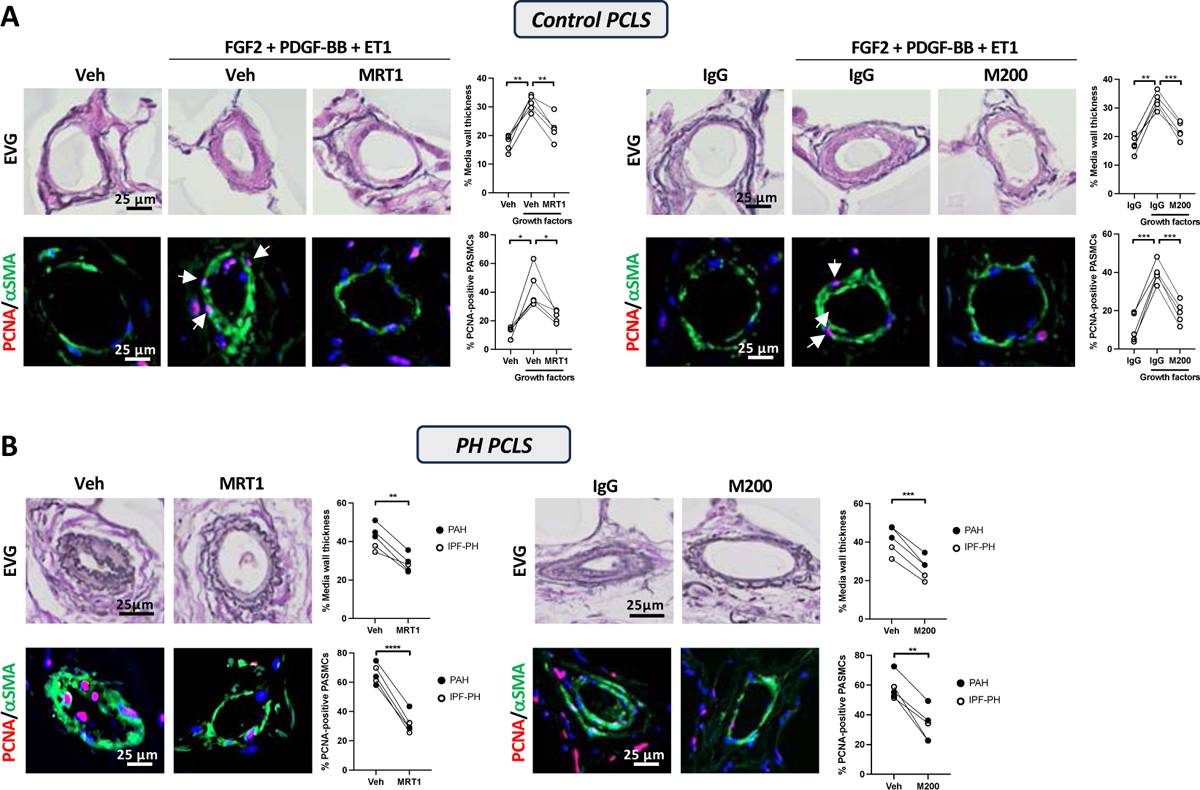
Inhibition of integrin a5 subunit reduces vascular remodeling in human precision­cut lung slices (PCLS). (**A**) Representative images of distal PAs stained with Elastica van Gieson (EVG) or labeled with PCNA in PCLS prepared from control patients after exposure or not to a growth factor cocktail in presence or not to MRT1 (4 pM) or M200 (1 pg/ml) for 10 days. PASMCs were labeled with alpha-smooth muscle actin (aSMA, green). The quantification of vascular remodeling and PASMCs positive for PCNA are shown. Each point represents a patient and corresponds to the average of! 13-15 arteries per slide (n=5; *p<0.05, **p<0.01, ***p<0.001, Repeated measures one-way ANOVA followed by Dunnett’s post hoc analysis). Scale bars, 25 pm. (**B**) Representative images of distal PAs stained with EVG or labeled with PCNA in PCLS from patients with pulmonary hypertension (PH; PAH or PH secondary to idiopathic pulmonary fibrosis (IPF-PH)) after exposure to MRT1 (4 pM) or M200 (1 gg/ml) for 10 days. Each point represents a patient and corresponds to the average of 13-15 arteries per slide (n=5; *p<0.05, **p<0.01, Paired Student’s t-test). Scale bars, 25 pm.

## DISCUSSION

PAH is a multifactorial and complex disease in which PASMCs and PAECs expansion play a dominant role in vascular obliteration. In this study, we identified the α5β1 integrin as a key orchestrator of PAH-PASMCs and PAH-PAECs behavior. Indeed, using a small molecule integrin inhibitor and neutralizing antibody, together with employing diverse preclinical models and adopting a multi-batch design for drug testing, alongside comparison with standard of care and external validation, we present compelling evidence supporting the potential utility of α5β1 integrin inhibition as a promising therapeutic approach for patients with PAH.

While several RGD-binding integrins have attracted considerable interest as therapeutic targets in pulmonary fibrosis and asthma ^38–41^ relatively little attention has been paid to PAH, although downstream signaling mediators of integrins, such as the TKs FAK/c-Src complex, Akt, ERK and ILK have been documented to contribute to pulmonary vascular remodeling^12, 27, 42^.

Due to the important role of integrins in regulating a plethora of cellular processes that are key in the pathophysiology of PAH, selective inhibition of integrins is the ideal approach to increase the therapeutic success. Among the integrin subunit genes surveyed, we found that PAH-PASMCs and PAH-PAECs have higher levels of α5 and correspondingly increased expression of α5β1 compared to control cells. *In vitro* experiments demonstrated that blockade of α5β1 as well as knockdown of α5 significantly reduced the proliferative capacity of disease cells. These results are consistent with published data showing that α5β1 is required for ECM-induced proliferation of normal PA adventitial fibroblasts and PASMC s^43^. Although the promoting effects of α5β1 on cell survival and proliferation likely rely on the activation of a variety of converging signaling pathways, we found that inhibition of α5β1 impaired FAK signaling and the activation of the transcription factor FOXM1 resulting in a mitosis defect as evidenced by mitotic spindle abnormalities and binucleation. In this regard, several studies have shown that integrins are key regulators of mitotic events in directing the spindle parallel to the adhesion plane. Indeed, Reverte and colleagues documented that decreased integrin β1 activity allows entry into mitosis but inhibits microtubule assembly from the centrosome, leading to a cytokinesis defect by preventing the formation of a normal bipolar spindle ^44^. Consistent with this, integrin-mediated adhesion via FAK signaling has been shown to promote centrosome separation in early mitosis^45^. All these effects on mitosis are supported by our findings showing an α5β1-dependent activation of transcription factor FOXM1 in PAH-PASMC. FOXM1 is a master regulator of mitosis implicated in PAH ^46–50^ and activated by the TKs regulated by α5β1 i.e. FAK, c-Src and Akt ^29, 50–52^.

In addition to demonstrating that α5β1 supports the pro-proliferative and apoptosis-resistant phenotype of PAH-PASMCs and PAH-PAECs *in vitro*, we have convincingly and repeatedly shown that small molecule RGD-integrin inhibitor (i) confers therapeutic benefits in multiple preclinical animal models of the disease and (ii) enhances the effects of current standards of care for PAH. These effects are likely driven by α5β1 blockade, supported by the replication of beneficial outcomes upon inhibition using species-specific blocking antibodies. In contrast, blocking both αvβ1 and α8β1 with MRT6 had no discernible impact, highlighting the significant role of α5β1 in pathological vascular remodeling in PAH. Although not directly explored in our study, previous research has shown that inhibition of α5β1 can reduce airway smooth muscle contraction ^41^. Therefore, beyond its impact on vascular remodeling, α5β1 inhibition in the pulmonary circulation might also mitigate PAH by promoting vasodilation. However, our findings suggest that α5β1 inhibition offers more than just a vasodilatory effect. This is supported by the observation that α5β1 inhibition provided additional benefits beyond the combination of macitentan and tadalafil, two major vasodilatory drugs used in PAH, and provided similar efficacy to imatinib further confirming its anti-remodeling effects.

The pulmonary circulation and the RV form an integrated functional unit in PAH, and documentation of the direct beneficial or deleterious effects of a new compound in the stressed RV is often lacking. This is even more important as the available literature is limited and mostly focused on β1^53, 54^, which combines with multiple alpha subunits, making it difficult, if not impossible, to extrapolate its role and confront previous data. Nevertheless, our results showing that MRT1 treatment has no cardiotoxic effect and even confers protection against PAB-induced progressive deterioration of RV function add further weight to the fact that α5β1 targeting holds great promise. This is in line with published data showing that fibronectin contributes to pathological cardiac remodeling^55, 56^ and that α5β1 is upregulated in the neointima following arterial injury ^57^. However, further in-depth and targeted investigation into the role of α5β1 in cardiac function is warranted in future research endeavors.

Some limitations of our study should be noted. Based on the multiple functions fulfilled by integrins, it is expected that α5β1, in addition to promoting cell survival and division, mediates multiple aspects of small PAs remodeling. In this regard, integrin α5 has been reported to stimulate proinflammatory gene expression in vascular cells^58, 59^, a hallmark and driving force of pulmonary vascular remodeling^60^. Finally, although our data suggest that integrin α5β1 is required for FOXM1 activation and for centrosome separation and proper execution of mitosis, additional candidate effectors/signaling pathways that are involved in α5β1 integrin regulation of PAH cell behavior remain to be characterized.

In conclusion, this reveals a hitherto unappreciated role of α5β1 in pathological lung remodeling in the context of PAH and lends credence to α5β1 antagonists for clinical translation.

## MATERIALS AND METHODS

The full experimental details are presented in the supplemental materials.

### Human tissues

Experimental procedures using human tissues or cells conformed to the principles outlined in the Declaration of Helsinki and were performed with the approval of Laval University and the Institut Universitaire de Cardiologie et de Pneumologie de Québec (IUCPQ) Biosafety and Ethics Committees (CER #20773 and #22390). The characteristics of patients are described in Supplemental Tables S1 to S3.

### Animal models of pulmonary hypertension (PH)

All animal studies were approved by the Animal Ethics Committee of Université Laval (#2023-1317 and 2022-1049) and in accordance with the guidelines of the Canadian Council on Animal Care.

### Statistical analysis

Statistical analyses were conducted using GraphPad Prism software (version 9.3.1). To evaluate the normality of the data, the Shapiro-Wilk test was employed. The statistical significance of differences between two groups was determined using either the paired or unpaired Student’s t test or the corresponding non-parametric alternative, depending on the data’s distribution. For analyses involving multiple comparisons, one-way or two-way ANOVA or the Kruskal-Wallis’ analysis were applied, with subsequent post-hoc testing using Dunn’s, Dunnett’s or Tukey’s methods as appropriate. For datasets involving paired measures, a repeated-measures ANOVA was performed, followed by post-hoc analysis with Dunnett’s test. Longitudinal comparisons among groups were assessed using mixed linear models. All findings are reported as the mean ± standard error of the mean (SEM). A p-value of less than 0.05 was considered statistically significant. Significance is indicated as * p<0.05, **p<0.01, ***p<0.001 and ****p<0.0001.

## Acknowledgements

We thank the IUCPQ Biobank of the Quebec Respiratory Health Research Network as well as the department of cytology and pathology from the IUCPQ for providing access to tissue and clinical data. We extend our gratitude to Elizabeth H. Konopka, Hailey Rapisardi, Inese Smukste, Jessica Pondish, Meghan Monroy, Minh Hai Nguyen, Parmita Saxena, Sujan Lama, Laura Cappellucci, Kevin McManus and Woojin Kim for their invaluable contribution and collaboration.

## Author contributions

Conceptualization: SEL, MSM, ASR, ML, SP, OB, SB

Methodology: SEL, YG, TY, TS, CR, MS, SM, SBB, AB, CT, AP, REK, FP

Investigation: SEL, YG, TY, TS, CR, MS, SM, SBB, AB, CT, AP, REK, SM, KY, FYL, HW, JED, EHK, IS, JP, MM, MHN, SL, LC, KM, WK

Visualization: SEL, SP, OB, SB Funding acquisition: SB Supervision: SP, OB, SB

Writing – original draft: SEL, SP, OB, SB

Writing – review & editing: SEL, MSM, ML, ASR, BL, BG, FYL, TIM, SP, OB, SB

## Data and material availability

All data associated with this study are present in the paper or Supplementary Materials. RNA sequencing data have been uploaded to the NCBI Gene Expression Omnibus (GEO) repository (……….). De-identified human/patient data are listed in Supplementary tables.

## SUPPLEMENTAL MATERIALS AND METHODS

### Human lung tissue acquisition

Experimental procedures using human tissues or cells conformed to the principles outlined in the Declaration of Helsinki and were performed with the approval of Laval University and the Institut Universitaire de Cardiologie et de Pneumologie de Québec (IUCPQ) Biosafety and Ethics Committees (CER #20773 and #22390). Tissues were obtained from patients who had previously given signed informed consent. PAH and control tissues were obtained from Respiratory Health Network tissue bank (www.rsr-qc.ca). PAH tissues were from lung explants from transplants or warm autopsies. PAH diagnosis was previously confirmed by right heart catheterization. Healthy lung tissues (controls) were obtained during lung resection for tumors. Lung samples were taken at distance from the tumor and demonstrated normal lung parenchyma. The characteristics of patients are described in Supplemental Tables S1 to S3. The pulmonary arteries smaller than 1 mm in outer diameter were isolated from both PAH and non-PAH patients and PASMCs and PAECs were enzymatically isolated as previously described ^1^.

### Ex vivo models of precision-cut lung slices

Human lung biopsies were obtained from control, PAH, and IPF patients. Lung tissues were perfused with 3% (m/V) low-melting agarose (Sigma-Aldrich) through visible vessels and bronchi. Lung biopsies (1cmx1cm) were then sectioned in 400µm width slices (Compresstome® VF-310-0Z Vibrating Microtome (Precisionary, USA). Lung slices were sequentially placed in a 24-well plate in 1 ml of medium. PAH and IPF lung slices were cultured in DMEM supplemented with 10% fetal bovine serum (FBS), 100 units/ml penicillin, 0.1 mg/ml streptomycin, 0.25 mg/ml amphotericin B and 0.03 mg/ml Gentamicin in presence of MRT1 (4µM) and M200 (1µg/ml) or their respective vehicles. Control lung slices were cultured in DMEM supplemented with 1% FBS, 100 units/ml penicillin, 0.1 mg/ml streptomycin, 0.25 mg/ml amphotericin B and 0.03 mg/ml Gentamicin. To reproduce vascular remodeling observed in PAH lung sections, control lung sections were exposed to PDGF-BB (30 ng/ml), FGF2 (30 ng/ml) and endothelin-1 (100 ng/ml) in presence of MRT1 and M200 or their respective vehicle. The culture medium was refreshed every 48 hours. On day 10, slices were fixed in 4% formaldehyde, embedded in paraffin, and processed for histological measurements. The characteristics of patients included in this study are described in Table S4.

### Animal models

All animal studies were approved by the Animal Ethics Committee of Université Laval (#2023-1317 and 2022-1049) and in accordance with the guidelines of the Canadian Council on Animal Care. The animals were housed in a 12-hour light-dark cycle with access to food and water *ad libitum*.

Three rodent models of PH were used in the present study. They included a rat model of Sugen-hypoxia (Su/Hx) induced PH, a rat model of monocrotaline (MCT)-induced PH and a mouse model of Su/Hx-induced PH. For rat models, adult male Sprague-Dawley rats (6–8 weeks, 200-250g in body weight) were purchased from Charles River laboratories. Rats were allowed 1 week to adjust to the new environment. In Su/Hx model, rats were injected subcutaneously with the vascular endothelial growth factor (VEGF) receptor inhibitor SU5416 (20 mg/kg, APExBIO) and were then exposed to hypoxia (10% O_2_) for 21 days. Then, these rats returned to normoxia for 2 additional weeks. Control rats were maintained in normoxic conditions. For the MCT rat model, one dose of MCT (60 mg/kg body weight, Sigma, St. Louis, MO, USA) was subcutaneously injected to rats, while control rats received an equal volume of saline. Regarding PH induction in mice, adult C57Bl/6J mice were weekly injected with SU5416 at 20 mg/kg once a week for 3 weeks. After the first injection of sugen, mice were exposed to hypoxia for 3 weeks followed by a return to normoxia for an additional 2 weeks.

MCT or Su/Hx rats with established PAH (3 weeks post-SU5416 injection or two weeks post MCT injection) were randomly assigned to receive MRT1 (100 mg/kg by oral gavage twice daily), MRT6 (10 mg/kg by oral gavage twice daily), mAb4 (10 mg/kg, given subcutaneously thrice a week), macitentan (1 mg/kg by oral gavage once daily, MedChemExpress), tadalafil (10 mg/kg by oral gavage once daily, MedChemExpress), imatinib (50 mg/kg by oral gavage given once daily), indicated drugs combination or corresponding vehicles for an additional period of 2 weeks. Su/Hx mice at the end of the 3-week hypoxic exposure were randomly divided into 6 groups that were treated with either MRT1 (100 mg/kg by oral gavage twice daily), 339.1 (10 mg/kg, given subcutaneously thrice a week, ATUM Bio), macitentan (15 mg/kg by oral gavage twice daily, MedChemExpress), 339.1+macitentan, or corresponding vehicles for an additional period of 2 weeks.

To interrogate RV effects of therapies independent of pulmonary vascular changes, the pulmonary artery banding (PAB) rat model was used. Adult male Sprague-Dawley rats (150-200 g in body weight) were randomly subjected to PAB or sham surgery. Briefly, rats were anesthetized with isoflurane and buprenorphine and then intubated. Following local anesthesia with lidocaine, a median sternotomy was performed, and the PA was dissected free from the aorta and left atrium. The main PA was banded with non-absorbable prolene sutures (6-0), tied tight against a 19-gauge needle, which was removed quickly. Sham-operated rats underwent the same procedure but without tying the pulmonary trunk. Three weeks post-PAB, rats were randomly assigned to receive MRT1 (100 mg/kg) or vehicle (PBS) by oral gavage twice per day for a period of 7 weeks. Rat plasmatic levels of alanine aminotransferase (ALT1), aspartate aminotransferase (AST), urea, and creatinine were dosed by the clinical biochemistry laboratory of our hospital using a Siemens Dimension Vista 1500 system.

For all animal experiments, only males were used to minimize potential hormonal effects and to reduce the experimental variability. No formal sample size and power calculations were conducted, because the primary goal of this study was to explore the effect of an intervention (α5β1 and αvβ1 inhibition) *in vivo* for the first time.

### Echocardiographic and hemodynamic measurements

Before treatment initiation and at the end of treatment, animals were anesthetized with 3%–4% isoflurane and maintained with 2% during procedures. Body temperature was maintained throughout imaging with the use of a heated platform and a heat lamp. Transthoracic echocardiography was performed in rodents using a Vevo 3100 (Visualsonics) imaging system. Pulmonary hemodynamics were assessed through images acquired from parasternal short-axis and long-axis views, as previously described^2^ and according to current guidelines^3^. Next, hemodynamic parameters, including RV systolic pressure (RVSP) and mean PA pressure (mPAP) were measured blindly by closed chest right heart catheterization (RHC) (SciSence catheters). Briefly, the right jugular vein was cannulated, and a polyethylene pressure catheter connected to a pressure transducer (SciSence catheters) was advanced into the RV and the PA. After hemodynamics had been recorded, the animals were sacrificed, and their tissues were harvested.

### Measurement of RV hypertrophy

After sacrifice, the heart was excised and RV and the left ventricle plus septum (LV+S) were carefully dissected before subsequent analysis. The extent of RV hypertrophy was determined by dividing the weight of the right ventricle by the sum of left ventricle and septum [RV/(LV+S)].

### Protein crystallization and structure determination

Recombinant human ⍺5β1 protein production and crystallization were performed based on protocols from a previous publication^4^. In brief, the four-domain α5β1 integrin protein was produced in Expi293 GnTI^-/-^ cells (Thermo-Fisher). Complex crystal structure was obtained by soaking the *apo* crystal overnight with 1 mM MRT1 in 100 mM Tris pH 7.1, 150 mM NaCl, 1 mM MgCl2, 1 mM CaCl2, 25% P3350, 10% P400. Diffraction data was collected at NSLSII and processed by XDS. The structure was solved by molecular replacement, and model building and refinement were done with Phenix^5^ and Coot^6^ (Table S12). PyMOLwas used for generating Figure 2B.

### Fluorescence Polarization (FP) assays

FP assays were used to measure compound activity through binding competition with purified integrin protein and a fluorescent ligand probe. For α5β1, αvβ1, αvβ3, and αIIbβ1, the disulfide bridged cyclized fluorescein-labeled peptide ACRGDGWCG was used as the probe^4^. For α1β1, α2β1, α3β1, α6β1, α7β1, α9β1, α10β1, and α11β1, the TAMRA-labeled peptide compound from R&D Systems, catalog # 6048, was used as the probe. For α4β1, the fluorescein-labeled peptide compound from R&D Systems, catalog # 4577, was used as the probe. For α4β7, the disulfide bridged cyclized peptide CRSDTLCGEK-fluorescein was used as the probe^7^. For α8β1, the fluorescein-labeled peptide PQKPRGDVFIPRQPTNDLFEIFEIER was used as the probe^8^. For αDβ2, αLβ2, αMβ2, and αXβ2, the fluorescein-labeled compound 1 was used as the probe^9^. For αvβ6 and αvβ8, the fluorescein-labeled peptide compound GRGDLGRL was used as the probe^10^. Compounds were tested in FP assays containing 20 mM HEPES buffer at pH 7.3, 150 mM sodium chloride, 2 mM manganese chloride, 0.1 mM calcium chloride, 0.01% Triton X-100, 2% dimethyl sulfoxide, 3 nM of the fluorescent probe, and the purified extracellular domain of the integrin protein. The integrin protein concentration was the following: 14 nM α1β1, 7 nM α2β1, 14 nM α3β1, 2 nM α4β1, 7 nM α4β7, 2 nM α5β1, 15 nM α6β1, 40 nM α7β1, 6 nM α8β1, 16 nM α9β1, 17 nM α10β1, 30 nM α11β1, 7 nM αVβ1, 3 nM αVβ3, 15 nM αVβ5, 1.0 nM αVβ6, 20 nM αVβ8, 250 nM αIIbβ3, 2 nM αDβ2, 4 nM αLβ2, 6 αMβ2, and 7 nM αXβ2. Assays were performed in 384-well plates in a volume of 20 µL per well, and the integrin protein was pre-incubated with the test compounds for 15 minutes at 22⁰C before the fluorescein-labeled peptide was added. After the fluorescein-labeled peptide was added, the assay was incubated at 22⁰C for 1 hour and fluorescence polarization was measured. IC_50_ values were determined by nonlinear regression, 4-parameter curve fitting.

### MDCK Permeability Assay

The permeability assay used Madin-Darby canine kidney (MDCK) cells transfected with human MDR1 gene (MDCK-MDR1) encoding for P-glycoprotein (hP-gp, ABCB1). A test article (5 μM), atenolol (low permeability control, 5 μM), metoprolol (high permeability control, 5 μM), or quinidine (P-gp substrate, 5 μM) in the Hanks’ Balanced Salt Solution (HBSS) transport buffer (pH 7.4; containing 0.4% DMSO) were placed in either the apical side (A) (0.4 mL) or the basolateral side (B) (0.8 mL) of the Transwell plates as the donor chamber, respectively. The compound dosing solutions were centrifuged at 4000 rpm for 5 min before loading to donor chambers. The transport buffer was placed in the corresponding B side (0.8 mL) for A-to-B direction or A side (0.4 mL) for B-to-A direction as the receiver chamber, respectively. Lucifer Yellow (LY, 5 μM) and P-gp inhibitor elacridar (10 μM) were added to the apical side of all Transwells. The plates were then incubated for 90 min at 37°C in an atmosphere of 5% CO_2_ and 90% humidity. At the end of the incubation, 6 μL of the medium in each donor chamber, and 60 μL of the medium in each receiver chamber were taken for Liquid chromatography–tandem mass spectrometry (LC-MS/MS) analysis. Donor samples were diluted with 54 μL of the assay buffer, and then 60 μL of acetonitrile with internal standard (IS) were added to the donor and receiver chamber samples. Then all the samples were analyzed by LC-MS/MS. Lucifer Yellow permeability was used to determine monolayer integrity for each Transwell.

### Generation of rat anti-α5β1 antibody and specificity/affinity measurement

Antibodies against rat α5β1 were raised via immunizing SJL mice with rat α5β1 integrin ectodomain. The rat α5β1 ectodomain protein was expressed recombinantly in expi293 cells and purified using Ni-NTA, Strep-Tactin XT, and subsequently size-exclusion chromatography. Hybridoma method was conducted to generate multiple clones demonstrating specific binding to rat α5β1. Screening of hybridoma supernatants was conducted through binding ELISA (enzyme-linked immunosorbent assay) against rat α5β1 and rat αvβ1 to identify clones binding to rat α5 subunit selectively. The clones showed specific binding to rat α5β1 and then underwent additional screens in both protein-based and cell-based fibronectin-blocking assays. The top functional clones were subcloned and sequenced. Clone mAb4 was identified as the top candidate based on its high affinity and high potency in cell-based assays. Subsequently, mAb4 was recombinantly expressed on the rat IgG1 backbone with a N297G mutation using CHO cells for *in vivo* studies. The endotoxin level of the recombinantly expressed mAb4 and 339.1 antibodies are below 5 EU/mg.

### Negative Staining Electron Microscopy

Negative stain EM results were obtained by NanoImaging Services, Inc. Rat α5β1 ectodomain protein was mixed with antibody or Fab in a 1:5 molar ratio and incubated for 5 minutes at room temperature, and then diluted to final concentration 8-30 μg/ml with a buffer contains 10mM HEPES pH 7.3, 150mM NaCl, 1mM MgCl2 and 1mM CaCl2. A 3μl drop of sample suspension is applied to an EM grid and stained with 1% uranyl formate. 1000-1500 micrographs were collected using a Thermo Fisher Scientific (Hillsboro, Oregon) Glacios Cryo Transmission Electron Microscope operated at 200kV and a nominal magnification of 92,000. Data processing was performed using cryoSPARC3.3^11^.

### Preclinical pharmacology and pharmacokinetic (PK) properties

The pharmacokinetic profile of MRT1 was evaluated in male Sprague Dawley rats (6-8 weeks old). Water was available to all animals *ad libitum.* All animals from PO studies were fasted overnight and fed 4 hours post-dose. MRT1 was formulated as a solution in 20 mM citric acid buffer and dosed via oral gavage. Approximately 150 µL of whole blood was collected from all animals via submandibular vein or tail vein at pre-dose, 0.083, 0.25, 0.5, 1, 2, 4, 8 and 24 hr post-dosing into the test tubes containing anticoagulant potassium ethylenediaminetetraacetate (K_2_EDTA). Blood samples were centrifuged at 2,000 g for 5 minutes at 4°C to obtain plasma samples. All samples were stored at approximately −70°C until analysis. Plasma samples were mixed with acetonitrile containing internal standard and centrifuged at 5,800 rpm for 10 minutes. The supernatant was used for analysis. The concentration of MRT1 in plasma samples was determined using liquid chromatography with tandem mass spectrometry (LC-MS/MS) based methods.

The pharmacokinetic profile of mAb4 was evaluated in male Sprague Dawley rats (6-8 weeks old). Water and food were available to all animals *ad libitum*. mAb4 was formulated as a solution in PBS for SC dosing. Approximately 125 µL of whole blood was collected from all animals via jugular vein at 1, 4, 8, 24, 48, 72, 96, 168 and 336 hr post-dosing. Blood was collected into 300 μL red cap serum tubes (Sarstedt-20.1308.100) and kept at ambient temperature until fully clotted (for up to 30 minutes) and then centrifuged at 1300 g for 15 minutes in a refrigerated centrifuge. The resultant serum was transferred to polypropylene tubes, immediately frozen on dry ice, and stored at −80 °C until analysis using Meso Scale Discovery (MSD) based method.

### Cell-based Ligand Binding Assays (LBA)

Assays against human integrins were performed on K562 cells that natively express α5β1 or on HEK293 cells that stably express αvβ1, αvβ6 or αvβ8^12^. Assays against rat α5β1 were performed on α5-deficient CHO B2 cells^13^ that stably express rat α5 and rat β1. The cells were barcoded with 2 dyes to allow multiplexing during flow cytometry analysis^14^. Cells were washed in 50 mM HEPES pH 7.3 and 150 mM sodium chloride, and resuspended in 0.05% e780 Fixable Viability Dye in the assay buffer described below. Into wells of 96-well plates, 50,000 cells per well and inhibitor samples were incubated for 15 minutes at room temperature. Purified fibronectin (Sigma F2006) or proTGF-β1 protein fluorescently labeled with Dylight 650 (Abcam catalog # ab201803) were added to a concentration of 25 nM or 40 nM, respectively, and incubated for 45 minutes at room temperature. The assays were performed in the following assay buffers: 1) 95% human sera, 50 mM HEPES pH 7.3, 2) 95% human sera, 50 mM HEPES pH 7.3, 2 mM manganese chloride, 3) 95% rat sera, 50 mM HEPES pH 7.3, 2 mM manganese chloride, 4) 50 mM HEPES pH 7.3, 150 mM sodium chloride, 1% bovine sera albumin, 10 mM glucose, 0.7 mM magnesium chloride, 0.7 mM calcium chloride, 5) 50 mM HEPES pH 7.3, 150 mM sodium chloride, 1% bovine sera albumin, 10 mM glucose, 2 mM manganese chloride, 0.1 mM calcium chloride. 25 µl of 20% fixation buffer (BioLegend catalog # 420801) diluted in 50 mM HEPES pH 7.3 and 150 mM sodium chloride was added and cells were fixed for 30 minutes. 260 µl of wash buffer (50 mM Tris pH 7.5, 150 mM sodium chloride, 1% bovine serum albumin, and 1 mM EDTA) was added to the wells and then up to 6 assay plates were combined into a deep well plate. The deep well plate was centrifuged for 5 minutes at 500 x gravity, the supernatant was removed, and resuspended in wash buffer. Samples were analyzed using flow cytometry to quantify levels of Daylight 650. IC_50_ values were determined by nonlinear regression, 4-parameter curve fitting.

### Solid Phase Assays

Wells of 384-well plates were coated with 2 µg/mL mouse fibronectin (Abcam ab92784) or rat fibronectin (Sigma F0635) for 16 hours at 4°C, washed with washing buffer of PBS with 0.05% Tween 20, and blocked with 1% BSA in PBS for 2 hours at 37°C. After washing, antibody samples and 2 nM His-tagged mouse α5β1 protein or 8 nM His-tagged rat α5β1 protein were added in a solution of 50 mM HEPES pH 7.3, 150 mM sodium chloride, 1 mM magnesium chloride, 1 mM calcium chloride, and 0.05% Tween20 in a volume of 20 µL and incubated at room temperature for 1 hr. The wells were washed and the amount of bound His-tagged protein was detected using Anti His 6X detection antibody-HRP conjugated (Abcam Ab1187) and ELISA reagents (R&D Systems DY998 and DY994) as directed by the manufacturers.

### Cell Adhesion Assays

CHO B2 cells expressing rat α5β1 or mouse C2C12 cells, which natively express mouse α5β1, were incubated with the antibody samples for 15 minutes at room temperature in assay buffer (50 mM HEPES pH 7.3, 150 mM sodium chloride, 1% BSA, 10 mM glucose, 1 mM magnesium chloride, 1 mM calcium chloride) for the sera-free assays or in 50% rat sera with 50% assay buffer. In 96-well plates, wells were coated with 0.625 µg/mL GST-fibronectin (residues 168-368) protein and blocked with 1% BSA. The cells and antibodies were transferred to the wells and incubated for 1 hour at room temperature. Non-adherent cells were removed by inverted centrifugation in the BlueWasher (Blue Cat Bio) and adherent cells were quantified using CelTiter-Glo (Promega) as directed by the manufacturer.

### Cell culture and treatments

Human PASMCs and PAECs were isolated from dissected PA (<1mm) from control (n=9) and PAH (n=12) patients. Immunofluorescent staining was performed to determine the purity of the isolated PASMCs by using α-SMA and calponin 1 antibodies. The isolation of PAECs was achieved using magnetic beads coated with CD31 antibody. To ensure their integrity, an endothelial cell tube formation angiogenesis assay was conducted. PASMCs and PAECS were grown in smooth muscle cell basal medium supplemented with smooth muscle cell growth kit (Cell Applications) and in endothelial basal medium supplemented with Endothelial Cell Growth Medium-2 BulletKit (Lonza), respectively. Cells were maintained at 37°C in a 5% CO_2_ air humidified incubator. PAH-PASMCs were used at passages 5-10 and PAH-PAECs were used at passages 4-7. Cells were confirmed to be negative for mycoplasma. For all *in vitro* experiments, cell culture plates were first coated with fibronectin (1µg/ml) for 1 hour at 37°C, then washed once with PBS before seeding human PASMCs or PAECs. To explore the function of α5β1/αvβ1 integrins in PAH-PASMCs and PAH-PAECs, cells were treated with MRT1, M200 (Absolute antibody), MRT6 and PF-573288 (APExBIO) at indicated doses for 48 hours. Alternatively, cells were transfected in serum-free medium with a siRNA targeting ITGA5 (L-008003-00-0010, Dharmacon) or siFOXM1 ((#sc-270048, Santa Cruz Biotechnologies) at a final concentration of 10 nM or 50 nM, respectively with Lipofectamine RNAiMAX (Thermo Fisher Scientific) according to the manufacturer’s instructions. After transfection, cells were cultured in serum-containing medium for 48 hours for ITGα5 protein knockdown.

### Assessment of cell proliferation, migration and resistance to apoptosis

Proliferation and survival of PASMCs and PAECs were assessed after 48 hours post-drug exposure or siRNA transfection. To assess the resistance to apoptosis, cells were cultured in serum-free medium in presence or not of indicated drugs for 48 hours. After that, the percentage of apoptotic cells was quantified by Annexin V labeling (ApoAlert AnnexinV-FITC Apoptosis Kit, Takara Bio) and TUNEL (TdT-mediated dUTP Nick-End Labeling assay, Promega) according to the manufacturer’s instructions. For the assessment of cell proliferation, cells were cultured in 10% FBS-supplemented medium in presence or not of indicated drugs for 48 hours and were labeled with Ki67. Cell proliferation and apoptosis rates were calculated as the ratio of the mean numbers of Ki67-, Annexin V- and TUNEL-positive cells to DAPI-stained cells, respectively.

For the wound healing assay, PAH-PASMCs were seeded in 6-well plates and grown to confluence. A vertical and horizontal cross-shaped scratch was made with a sterile p200 pipette tip. The cells were washed twice with PBS to remove detached cells and then refilled with a standard media containing a low serum concentration to suppress proliferation. Cell movement into the wound area was examined and recorded using an EVOS *xl* digital inverted microscope (Fisher) at 0, 6, 9 and 12 hours. The wound areas (A) were measured using the Wound Healing Size Tool plugin^15^ on ImageJ v.2.1.0/1.53c software (NIH, Bethesda, MD, USA). The migration of PAH-PASMCs toward the wounds was expressed as the percentage of wound closure: % of wound closure = [(A_t=0h_ – A_t=Δh_)/A_t=0h_] × 100.

### Double thymidine block

Initially, PAH-PASMCs and PAH-PAECs were treated with 2 mM thymidine (Sigma-Aldrich) for 18 hours, followed by a 9-hour release period, and then subjected to a second 2 mM thymidine treatment for 17 hours. Subsequently, cells were released for an additional 10 hours in the presence of either MRT1, M200 or their corresponding vehicles. To evaluate the impact of siRNA against ITGA5 (siITGA5) on the process of mitosis, cells underwent transfection with siITGA5 or siSRCM 24 hours prior to the initiation of the double thymidine block procedure.

### Immunofluorescence on cells

Cells grown on coverslips were fixed for 15 minutes with 4% PFA. After fixation, coverslips were washed with PBS, followed by permeabilization with 0.1% Triton X-100 in PBS for 10 minutes. After permeabilization, cells were washed three times with 0.05% Tween20 in PBS (PBST) and were blocked with 10% goat serum in PBS for one hour. After blocking, cells were incubated with indicated primary antibodies (Table S5) diluted in 2% goat serum overnight at 4°C. Subsequently, cells were washed three times with PBS and incubated with appropriate fluorescent-dye conjugated secondary antibodies (Table S5) for 1 h at room temperature. After washing, coverslips were mounted on glass slides using DAPI (4′,6-diamidino-2-phenylindole) Fluoromount G. Antigen labeling was visualized using a digital slide scanner (Axio Scan Z1, Zeiss) or an Axio Observer microscope (Zeiss). Images were captured using the ZEN software system (Zeiss).

### NanoString Analysis Platform and integrins Profiling

Gene expression was investigated using the NanoString Technology (NanoString Technologies, Seattle, WA, USA). 100 ng of total RNA (20ng/μl) was used from isolated PAH-PASMCs (n=5). A custom-made panel containing all 26 human integrin subunits was used, including the corresponding housekeeping genes and internal positive controls, following manufacturer instructions. Normalization was performed using the geometric mean of four housekeeping genes (ACTB, GAPDH, POLR2D and PPIA) that were not affected by the experimental conditions. Gene expression data underwent imaging quality control and normalization checks before analysis. Count detection limit was determined using threshold based on negative controls. Data analysis was performed using the nSolver™ package (version 4.0)

### Quantification of α5β1 integrin level

Protein levels of α5 and β1 as heterodimers were quantified by the MesoScale Discovery custom electrochemiluminescence assay using the following biotinylated antibodies (Anti-Human Integrin Alpha 5 (CD49e), 13-0496-80, Thermo Fisher Scientific; Anti-Human Integrin Beta 1 (CD29), NBP2-36561, Novus Biologicals). Samples were prepared and analyzed by Morphic following the manufacturer’s instructions.

### RNA sequencing and analysis

RNA was isolated from cells using the *Direct-zol RNA MicroPrep* Kit (*Zymo Research*) according to the manufacturer’s instructions. DNase I (RNeasy MinElute Cleanup kit; QIAGEN) was used to remove genomic DNA contamination, while RNA integrity and quantity were assessed using a NanoDrop spectrophotometer (Thermo-Fisher). RNA sequencing was performed by the CHU de Québec-Université Laval Research Center genomics platform. Briefly, samples were processed using NEBNext Ultra II directional RNA library prep kit (New England Biolabs), with library preparations sequenced on the Novaseq 6000 Sequencing System (Illumina) to generate 150 base-pair paired end reads. FASTQ files from the Novaseq instrument were examined for sequencing quality and adapters contamination using MultiQC v1.13^16^. Reads with low quality were removed prior to further analyses and adapters sequences were removed from the reads using Trimmomatic v0.36^17^. Alignment of trimmed reads to the human genome (hg38) was performed using the STAR aligner software v2.7.10^18^ and the data was converted to gene-level expression using RSEM v1.3.1^19^. Gene counts from the datasets were then used to assess differential expression using DESEQ2 v1.32.0^20^. Transcripts that had low or no counts were filtered out prior to the differential expression analysis. We incorporated the cell line as a covariate in the DESEQ2 model to mitigate its influence on differential expression results. This approach enables us to differentiate true biological variation from the impact of the cell lines, thereby enhancing the precision of our gene expression analysis. High-throughput sequencing data used in this study have been deposited in NCBI’s Gene Expression Omnibus and are accessible through GEO Series accession number GSEXXXXX. An absolute fold change of ≥1.4 and adjusted p-values ≤0.05 were used to define genes whose expression was significantly altered.

### Bioinformatic Analysis

Gene set enrichment analysis (GSEA, v4.1.0) was performed on differentially expressed genes (DEG) preranked by gene-expression-based log2 fold change between treatment and vehicle^21^. Hallmark gene sets were applied to GSEA. The results with FDR below 0.05 were considered significantly differential pathways. In addition, DEGs were subjected to Gene Ontology (GO) enrichment analyses using ShinyGO (v0.76)^22^.

### Histological, immunofluorescence, and morphometric analysis

Tissues were fixed in 4% paraformaldehyde, embedded in paraffin, and serially sectioned at 5 µm. Tissue sections were deparaffinized with xylene, rehydrated through a graded ethanol series, and processed for Elastica van Gieson (EVG) or immunofluorescence staining as described below. For immunofluorescence staining, tissue sections were immersed in 10 mM sodium citrate buffer (pH 6.0) and heated in a microwaveable pressure cooker for 15 min. After antigen retrieval, sections were blocked with normal goat serum (10%) in PBS for 2h at room temperature. The sections were then incubated with the indicated primary antibodies (Table S5) in a humidified chamber at 4°C overnight. After washing in PBS, sections were incubated with appropriate fluorescent-dye conjugated secondary antibodies (Table S5) for 1 h at room temperature. Sections were mounted in DAPI (4′,6-diamidino-2-phenylindol) Fluoromount G and coverslipped for imaging. Sections were viewed microscopically using a digital slice scanner (Axio Scan Z1, Zeiss) or an Axio Observer microscope (Zeiss), and images were acquired using Zen system (Zeiss).

For the assessment of pulmonary vascular remodeling, EVG-stained lung tissue sections were used. External diameter and internal diameter of at least 15 PAs (vessels with double elastic lamina) per animal were randomly determined and recorded by an independent investigator blinded to the treatment regimen. This analysis was performed for the small PAs with external diameters inferior to 75 μm. Medial wall thickness was expressed as follows: Percent of wall thickness = [(external diameter–internal diameter)/external diameter] × 100. To assess proliferation and apoptosis within small PAs, lung tissue sections were stained for PCNA (proliferation marker) or cleaved caspase-3 (apoptosis marker). Alpha smooth muscle actin (αSMA) was used to labeled PASMCs. The percentage of PCNA- or cleaved caspase-3-positive PASMCs in distal PAs was calculated by examining 15 randomly selected distal PAs from each animal. Species- and isotype-matched IgG was used as a negative control instead of the primary antibodies. Masson trichrome stain was used to assess the degree of fibrosis in cardiac sections. Fibrosis was quantified on digitized images, on which, blue-stained tissue areas were expressed as percentage of the total surface area. Hematoxylin-eosin staining was utilized to evaluate hypertrophy in cardiomyocytes. The cross-sectional area of the cardiomyocytes was determined by manually outlining each cell. The results are expressed as the average area measured across 50-100 cardiomyocytes for each specimen.

### Western blotting

Total proteins were extracted from tissues or cells using a 2% Chaps protein extraction buffer supplemented with a protease/phosphatase-inhibitor cocktails (Roche). Protein concentrations were determined with the Bradford assay. Equal amounts of protein (15 μg/lane) were subjected to Sodium dodecyl sulfate-polyacrylamide gel electrophoresis. After electrophoresis, the separated proteins were transferred onto polyvinylidene difluoride (PVDF) or nitrocellulose membranes using a liquid blotting system (Bio-Rad). Membranes were then incubated for 1 hour at room temperature in a blocking buffer (0.1% Tween 20 in TBS (TBST)) containing 5% nonfat dry milk powder followed by an overnight incubation at 4°C with indicated primary antibodies (Table S5) overnight. After washing three times with TBST, membranes were incubated with appropriate horseradish peroxidase–conjugated secondary antibodies for 2 hours at room temperature (Table S5). Antibodies were revealed using ECL reagents (Perkin–Elmer) and signals were visualized using the imaging Chemidoc MP system (Bio-Rad Laboratories). Protein expression was quantified using the Image lab software (Bio-Rad Laboratories) and normalized to Amido black (AB).

### Quantitative Real-Time PCR

Total RNA was extracted using Trizol (Invitrogen), according to the supplier’s indications. The concentration and quality of RNA were then assessed using a NanoDrop 2000 instrument (NanoDrop 2000, Thermo Scientific, USA). Synthesis of cDNA was obtained using the qScript Flex cDNA Synthesis Kit (Quanta Bio). Quantitative PCR was performed on a QuantStudio 7 Flex real-time PCR system (Applied Biosystems) using SsoAdvanced Universal SYBR Green Supermix (Bio-Rad Laboratories). The exclusive amplification of the expected PCR product was confirmed by melting curve analysis and gel electrophoresis. The relative quantification was carried out using hypoxanthine phosphoribosyl transferase 1 (HPRT1) as reference gene. The primers sequences used are listed in Table S6. The 2^−ΔΔCt^ method was applied to assess the relative expression of all RNAs. All reactions were performed in triplicate.

### Statistical analysis

Statistical analyses were conducted using GraphPad Prism software (version 9.3.1). To evaluate the normality of the data, the Shapiro-Wilk test was employed. The statistical significance of differences between two groups was determined using either the paired or unpaired Student’s t test or the corresponding non-parametric alternative, depending on the data’s distribution. For analyses involving multiple comparisons, one-way or two-way ANOVA or the Kruskal-Wallis’ analysis were applied, with subsequent post-hoc testing using Dunn’s, Dunnett’s or Tukey’s methods as appropriate. For datasets involving paired measures, a repeated-measures ANOVA was performed, followed by post-hoc analysis with Dunnett’s test. Longitudinal comparisons among groups were assessed using mixed linear models. All findings are reported as the mean ± standard error of the mean (SEM). A p-value of less than 0.05 was considered statistically significant.

## SUPPLEMENTARY FIGURES

**Figure S1.**
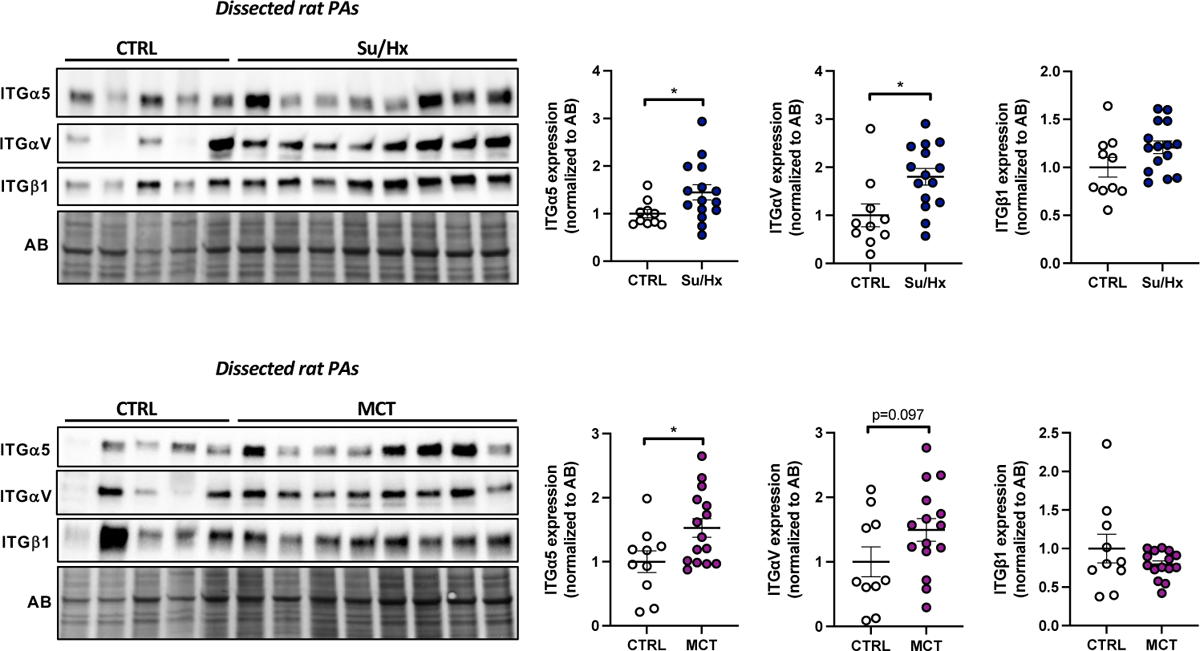
**(related to** Figure 1**). ITGa5 is upregulated across** *in vivo* **models of PAH.** Representative Western blots and corresponding quantifications of ITGa5, ITGaV, and ITGpi in dissected PA from rats exposed to either Sugen/Hypoxia (Su/Hx) or monocrotaline (MCT). Protein expression was normalized to amido black (AB) (n=10 to 15; *p<0.05, Mann-Whitney test in the Su/Hx rat model and unpaired Student’s t test in the MCT rat model; data represent mean ± SEM).

**Figure S2.**
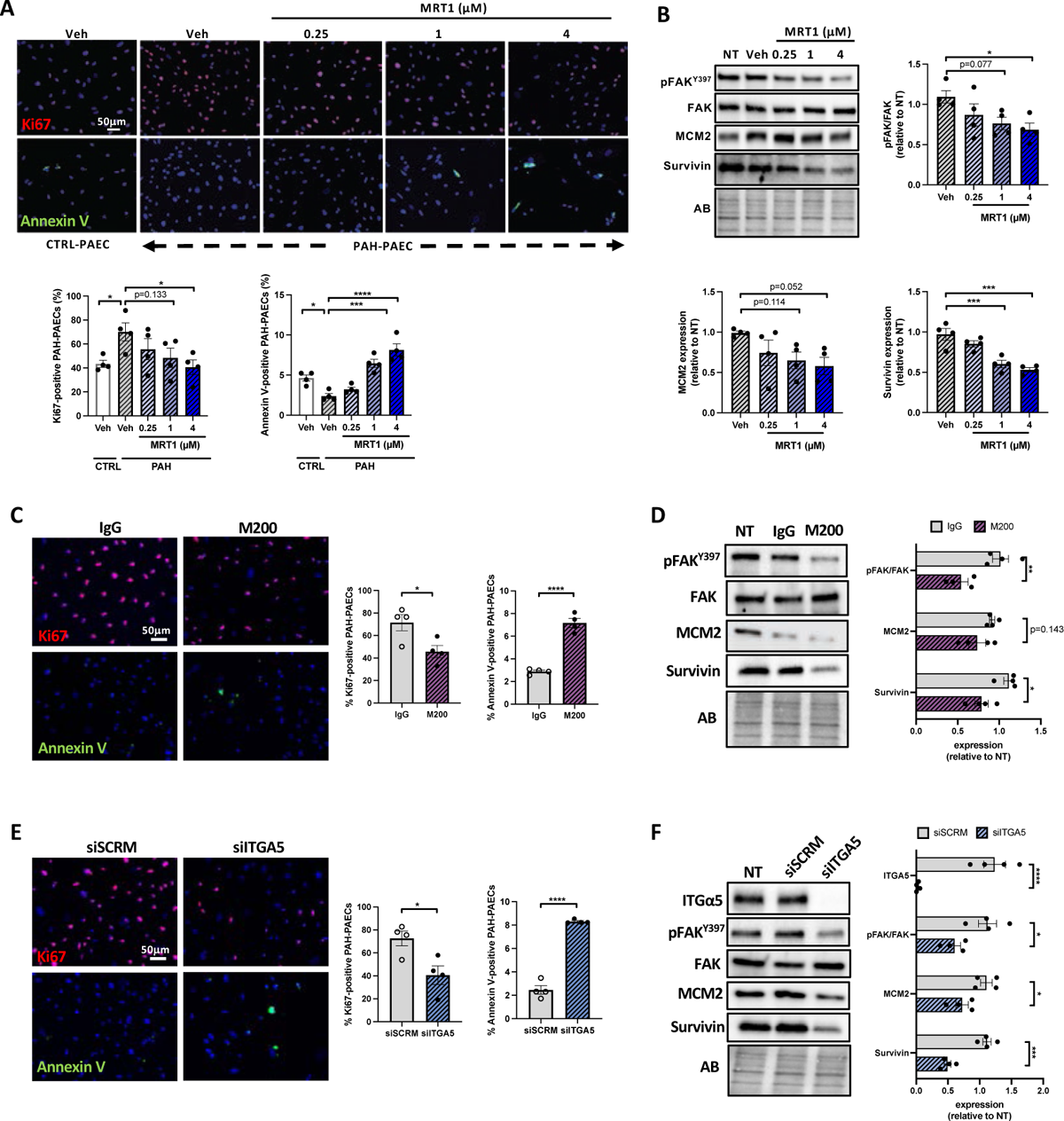
Inhibition of integrin a501 reduces the pro-proliferative and apoptosis-resistant phenotype of PAH-PASMCs and PAH-PAECs. (**A**) Representative fluorescent images of Ki67-labeled (red) and Annexin V-labeled (green) control and PAH-PAECs exposed or not to escalating doses of MRT1 for 48 hours. The corresponding quantifications are shown (n=4; *p<0.05, ***p<0.001, ****p<0.0001, one-way ANOVA followed by Dunnett’s post hoc analysis; data represent mean ± SEM). Scale bars, 50 pm. (**B**) Representative Western blots, and corresponding quantifications of the ratio of phospho-FAK (Y397) to FAK, MCM2, and Survivin in PAH-PAECs exposed or not to escalating concentrations of MRT1 for 48 hours. Protein expression was normalized to amido black (AB) (n= 4; *p<0.05, ***p<0.001, ****p<0.0001, one-way ANOVA followed by Dunnett’s post hoc analysis; data represent mean ± SEM). (**C**) Representative fluorescent images, and corresponding quantifications of Ki67-labeled (red) and Annexin V-labeled (green) PAH-PAECs exposed or not to M200 for 48 hours. (n=4; *p<0.05, ****p<0.0001, unpaired Student’s t test; data represent mean ± SEM). Scale bar: 50 pm. (**D**) Representative Western blots and corresponding quantifications of the ratio of phospho-FAK (Y397) to FAK, MCM2, and Survivin in PAH-PAECs exposed or not to M200 (0.5μg/ml) for 48 hours (n=4; *p<0.05, **p<0.01, unpaired Student’s t test for phospho-FAK (Y397) to FAK and MCM2 expression and Mann-Whitney test for Survivin expression; data represent mean ± SEM). (**E**) Representative fluorescent images, and corresponding quantifications of Ki67-labeled (red) and Annexin V-labeled (green) PAH-PAECs exposed or not to siITGA5 for 48 hours. (n=4; *p<0.05, ****p<0.0001, unpaired Student’s t test; data represent mean ± SEM). Scale bar: 50 μm. (**F**) Representative Western blots and corresponding quantifications of the ratio of phospho-FAK (Y397) to FAK, MCM2, and Survivin in PAH-PAECs exposed or not to siITGA5 for 48 hours (n=4; *p<0.05, ***p<0.001, ****p<0.0001 unpaired Student’s t test; data represent mean ± SEM).

**Figure S3.**
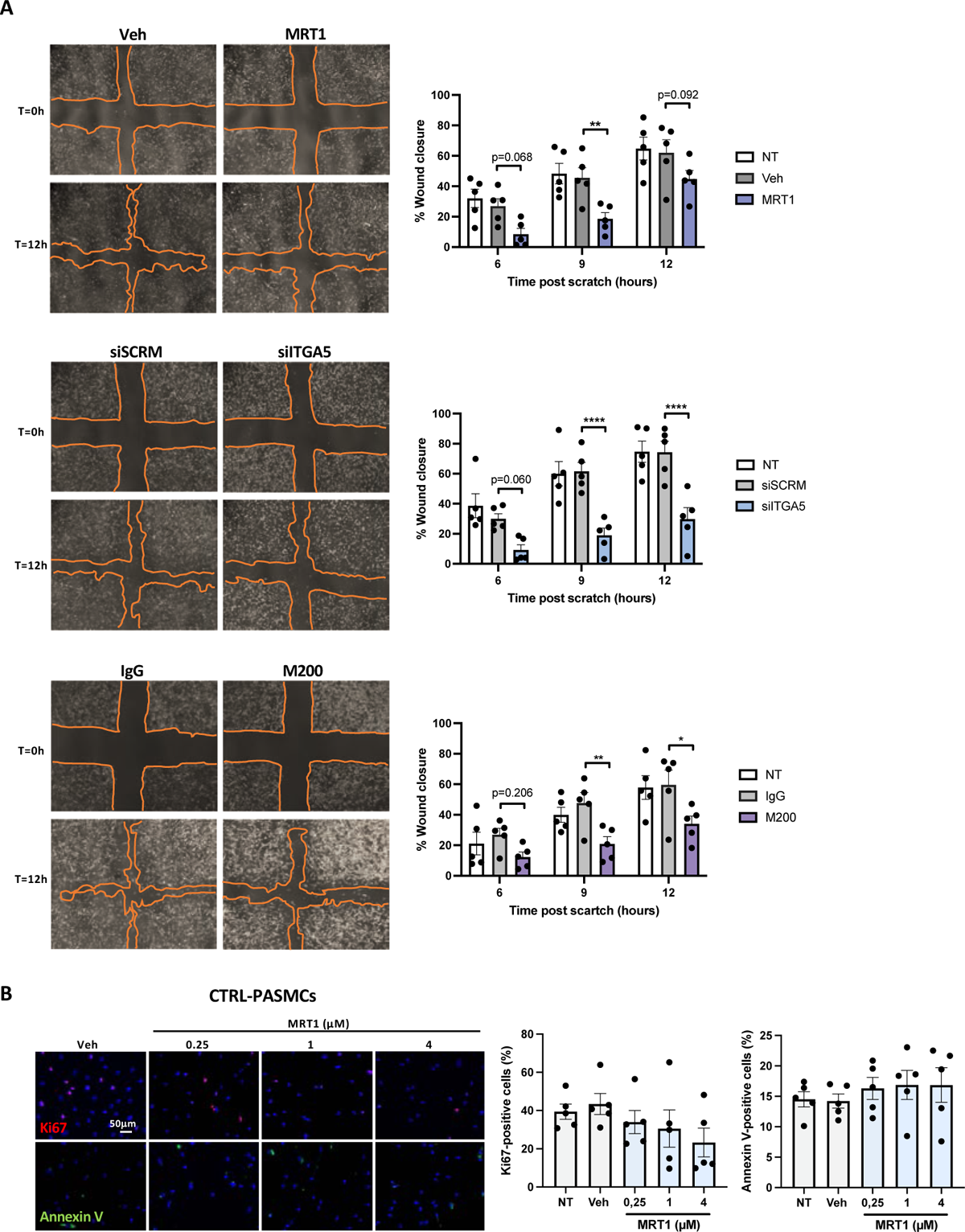
Inhibition of integrin a5pi decreases PAH-PASMCs migration and effect of MRT1 on CTRL-PASMCs. (**A**) Representative images from *in vitro* scratch wound assays demonstrating that CTRL-PASMC migration into the cell-free area (outlined) is significantly delayed in the presence of MRT1, siITGA5 or M200 when compared to their respective controls. (n=5; *p<0.05, **p<0.01, ****p<0.000, Two-way ANOVA followed by Tukey’s post hoc analysis; data represent mean ± SEM). (**B**) Representative fluorescent images of Ki67-labeled (red), Annexin V-labeled (green), and *TUNEL*-labeled (green) control and PAH-PASMCs exposed or not to escalating doses of MRT1 for 48 hours. The corresponding quantifications are shown (n=6; one-way ANOVA followed by Dunnett’s post hoc analysis; data represent mean ± SEM). Scale bars, 50 pm.

**Figure S4.**
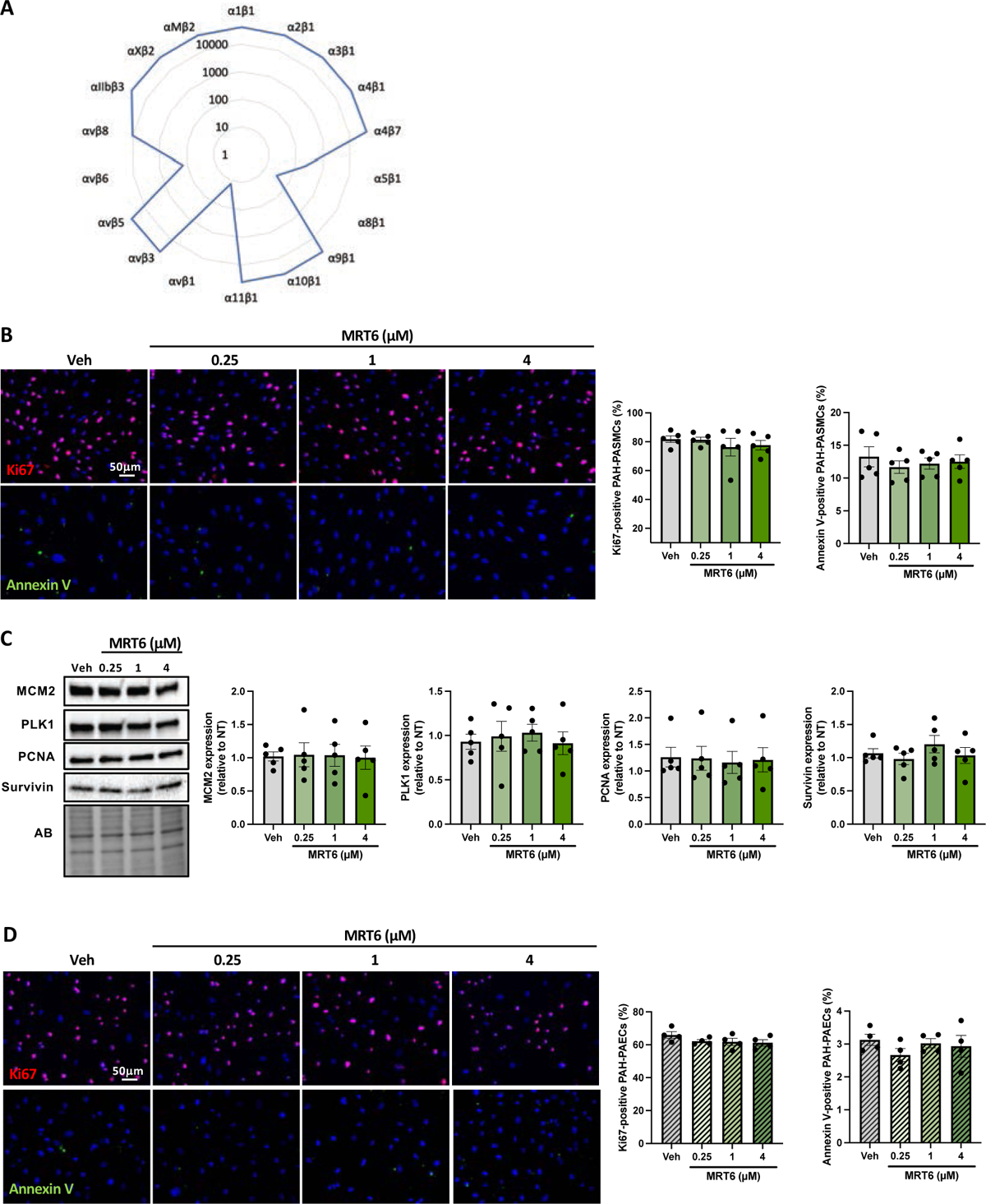
Inhibition of avpi/a8pi integrins using MRT6 had no impact on PAH-PASMC and PAH-PAEC behavior. (**A**) Radar plot of the inhibitory activity (in nM) of MRT6 against various integrins as determined through fluorescence polarization IC50 assays. (**B**) Representative fluorescent images of Ki67-labeled (red) and Annexin V-labeled (green) PAH-PASMCs exposed or not to escalating doses of MRT6 for 48 hours. The corresponding quantifications are shown (n=5; one­way ANOVA followed by Dunnett’s post hoc analysis; data represent mean ± SEM). Scale bar: 50 pm. (**C**) Representative Western blots, and corresponding quantifications of MCM2, PLK1, PCNA and Survivin in PAH-PASMCs exposed or not to escalating concentrations of MRT1 for 48 hours. Protein expression was normalized to amido black (AB) (n=5; one-way ANOVA followed by Dunnett’s post hoc analysis for MCM2 and PLK1 expression and Kruskal-Wallis’ test followed by Dunn’s post hoc analysis for PCNA and Survivin expression; data represent mean ± SEM). (**D**) Representative fluorescent images of Ki67-labeled (red) and Annexin V-labeled (green) PAH-PAECs exposed or not to escalating doses of MRT6 for 48 hours. The corresponding quantifications are shown (n=4; one-way ANOVA followed by Dunnett’s post hoc analysis; data represent mean ± SEM). Scale bar: 50 pm.

**Figure S5.**
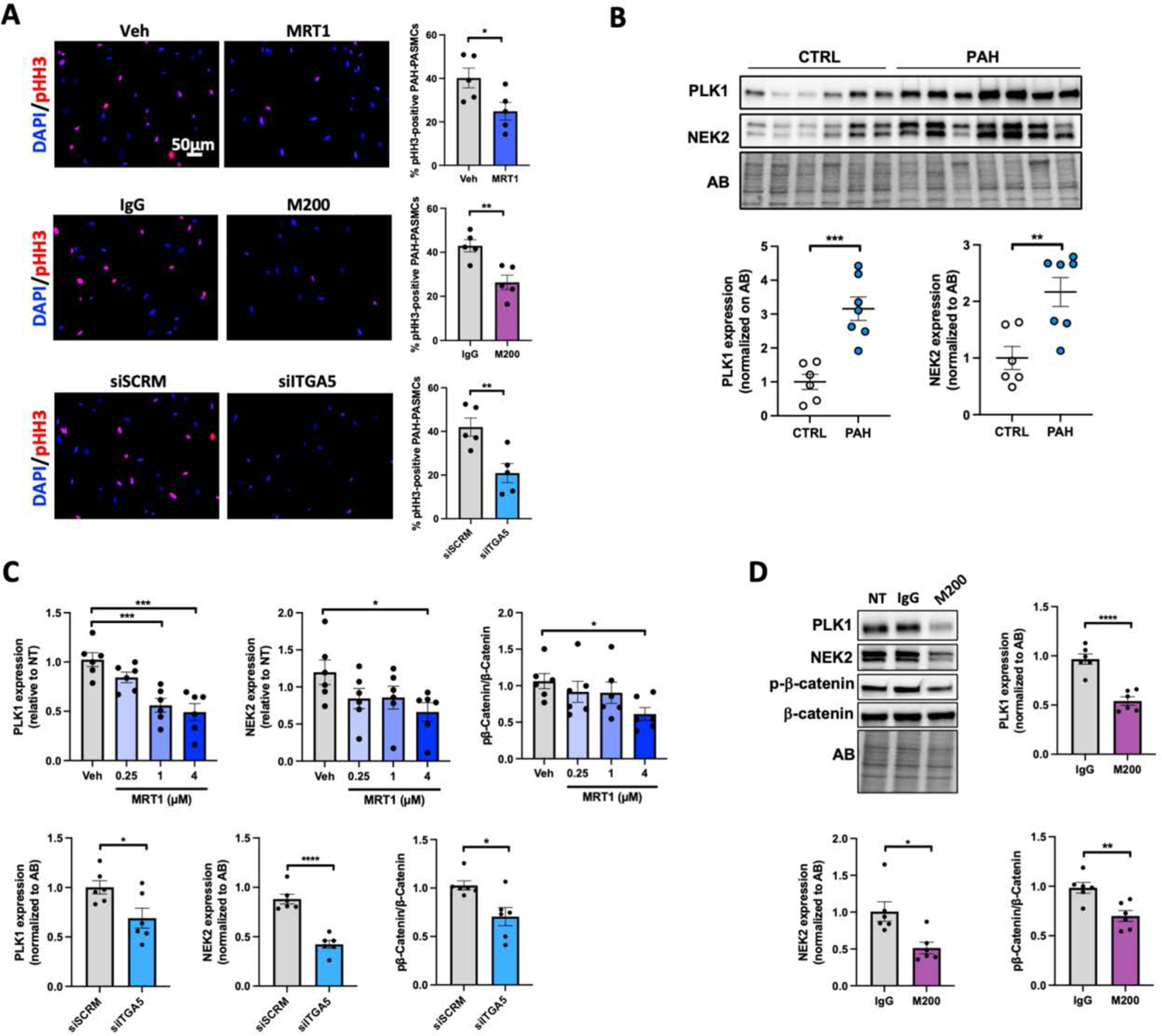
(**related to** Figure 3). Blockade of the integrin a5p1 or depletion of the a5 subunit results in mitotic defects in PAH-PASMCs. (**A**) Representative fluorescent images, and corresponding quantifications of pHH3-labeled PAH-PASMCs exposed or not to MRT1 (4 μM, 10 hours), M200 (0,5 pg/ml, 10 hours) or siITGA5 (transfected 24 hours prior to the double thymidine block initiation) (n=5; *p<0.05, **p<0.01, unpaired Student’s t-test; data represent mean ± SEM). Scale bar: 50 μm. (**B**) Western blots and corresponding quantifications of PLK1 and NEK2 in PASMCs isolated from healthy donors and patients with PAH. Protein expression was normalized to amido black (AB) (n=6 or 7; **p<0.01, ***p<0.001, unpaired Student’s t-test; data represent mean ± SEM). (**C**) Graphs related to Western blots presented in Figure 8H showing the quantification of PLK1, NEK2, and of the ratio of phospho-P-catenin to P-catenin in PAH-PASMCs exposed or not to escalating doses of MRT1 or siITGA5 for 48 hours. (n=6; *p<0.05, ***p<0.001, ****p<0.0001, MRT1: one-way ANOVA followed by Dunnett’s post hoc analysis, siITGA5: unpaired Student’s t test for PLK1 and NEK2 expression and Mann-Whitney test for the ratio of phospho-P-catenin to P-catenin; data represent mean ± SEM). (**D**) Representative Western blots and corresponding quantifications of PLK1, NEK2 and the ratio of phospho-P-catenin to P-catenin in PAH-PASMCs exposed or not to M200 (0.5μg/ml) for 48 hours. Protein expression was normalized to amido black (AB) (n=6; *p<0.05, **p<0.01, ***p<0.001, unpaired Student’s t test for PLK1 expression and the ratio of phospho-P-catenin to P-catenin and Mann-Whitney test for NEK2 expression; data represent mean ± SEM).

**Figure S6.**
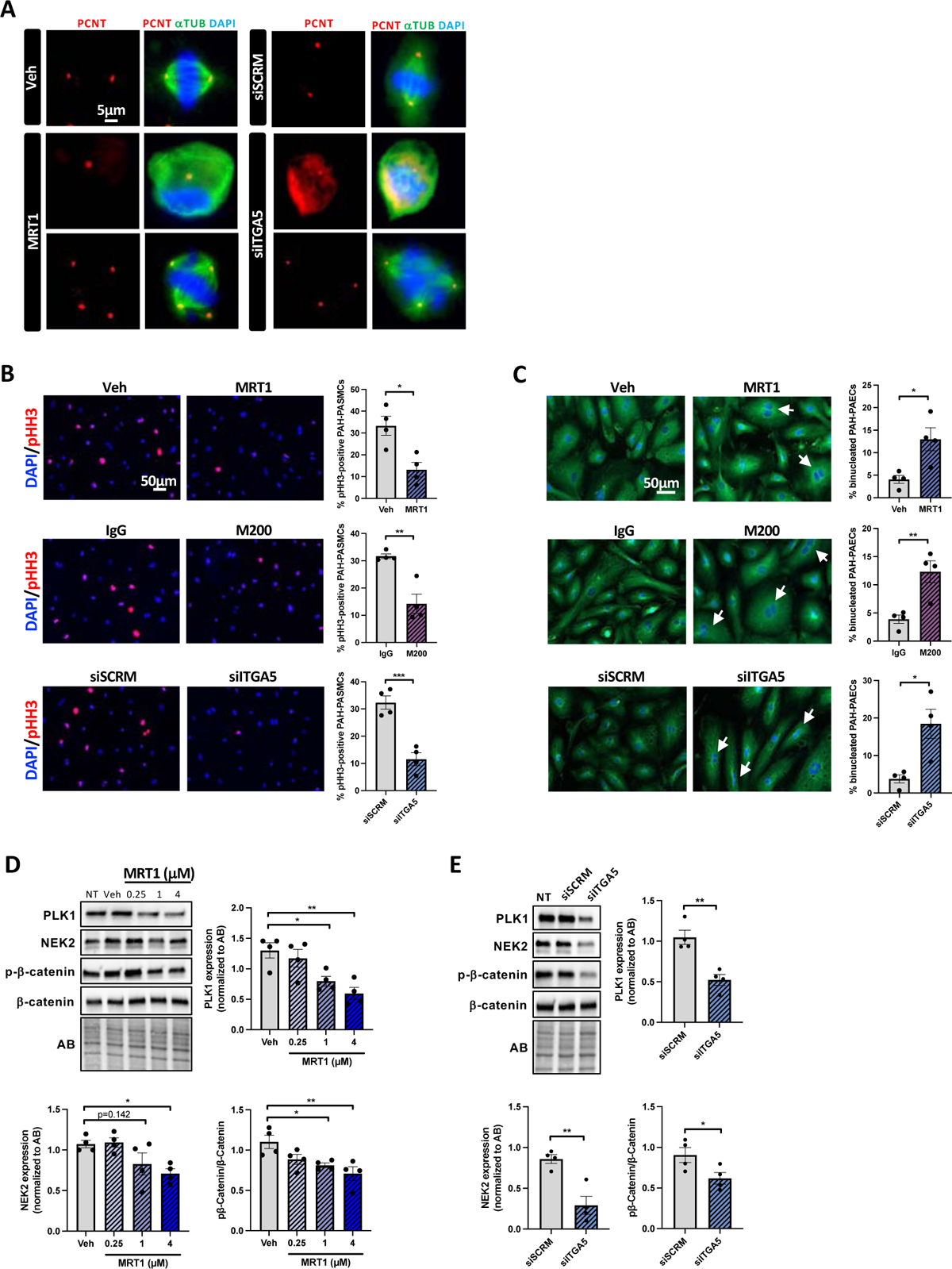
Blockade of the integrin a501 or depletion of the a5 subunit impairs mitosis in PAH-PAECs (Related to figure 3). (**A**) Representative images of the structure of the mitotic spindle in synchronized PAH-PAECs treated or not with MRT1 (4 pM, 10 hours) or siITGA5 (transfected 24 hours prior to the double thymidine block initiation). Red, green and blue in the image represent pericentrin, a-tubulin and DNA, respectively. Scale bar: 5 pm. (**B**) Representative fluorescent images, and corresponding quantifications of pHH3-labeled PAH-PAECs exposed or not to MRT1 (4 pM, 10 hours), M200 (0,5 μg/ml, 10 hours) or siITGA5 (transfected 24 hours prior to the double thymidine block initiation) (n=4; *p<0.05, **p<0.01, ***p<0.01, unpaired Student’s t test; data represent mean ± SEM). Scale bar: 50 μm. (**C**) Representative images and frequency of binucleated PAH-PAECs after treatment with MRT1 (4 μM, 10 hours), M200 (0,5 μg/ml, 10 hours) or siITGA5 (transfected 24 hours prior to the double thymidine block initiation) (n=4; *p<0.05, **p<0.01, unpaired Student’s t test; data represent mean ± SEM). Scale bar: 50 μm. (**D**) Representative Western blots and corresponding quantifications of PLK1, NEK2, and the ratio of phospho—P—catenin to P—catenin in PAH-PAECs exposed or not to escalating doses of MRT1 for 48 hours. Protein expression was normalized to amido black (AB) (n=4; *p<0.05, **p<0.01, one-way ANOVA followed by Dunnett’s post hoc analysis; data represent mean ± SEM). (**E**) Representative Western blots and corresponding quantifications of PLK1, NEK2, and the ratio of phospho-P-catenin to P-catenin in PAH-PAECs transfected or not to with siITGA5 for 48 hours. Protein expression was normalized to amido black (AB) (n=4; *p<0.05, **p<0.01, unpaired Student’s t-test; data represent mean ± SEM).

**Figure S7.**
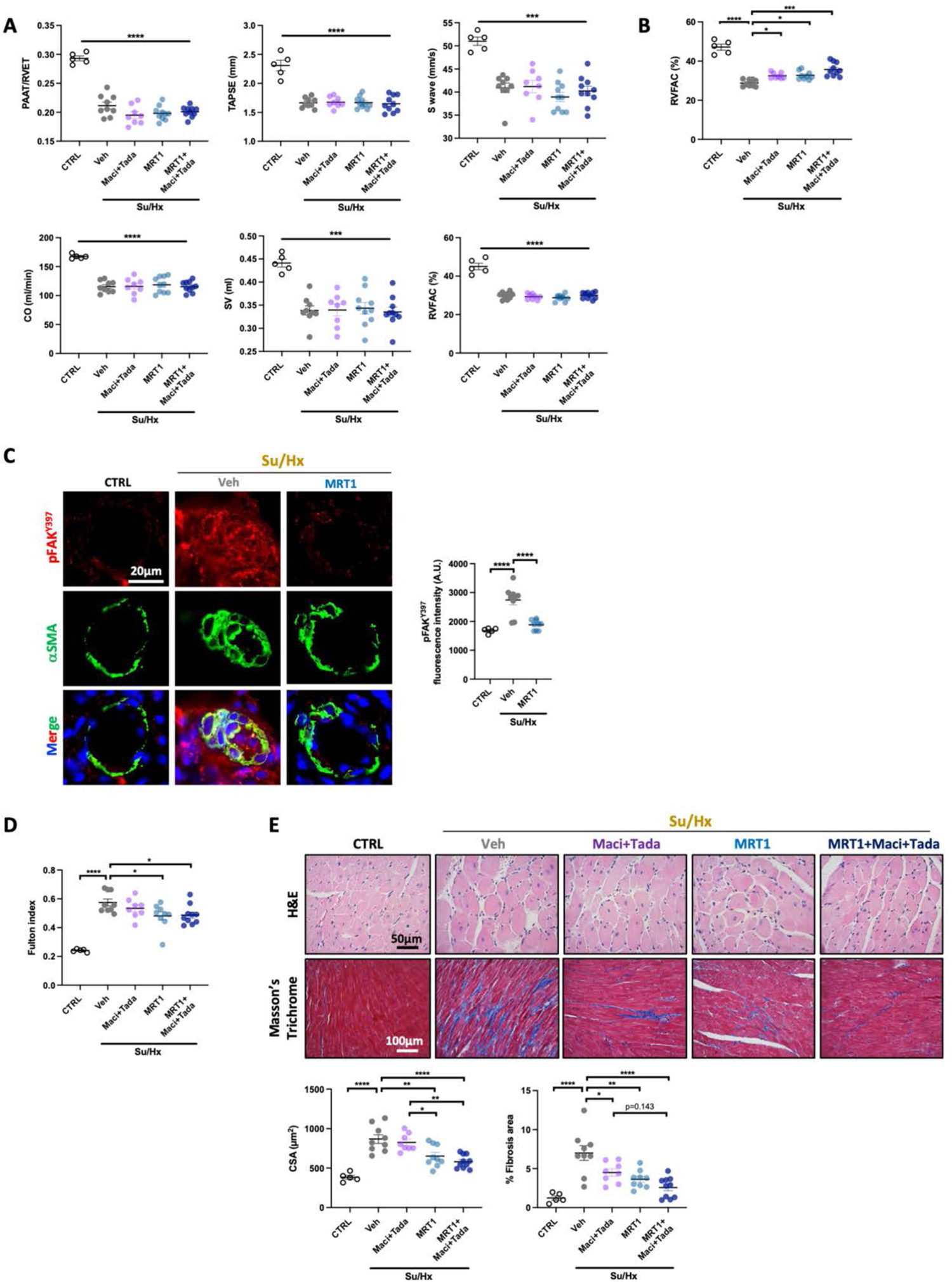
**(related to** Figure 5). Treatment with a small molecule inhibitor targeting RGD-integrins alone or combined with macitentan and tadalafil attenuates established PAH in Su/Hx rats. (A) PAAT/ET, TAPSE, S wave, CO, SV, and RVFAC measured using echocardiography in control and Su/Hx rats treated with vehicle, macitentan+tadalafil, MRT1 or MRT1+macientant+tadalafil before treatment initiation (n=5 to 10; ****p<0.0001, one-way ANOVA followed by Dunnett’s post-hoc analysis; data represent mean ± SEM). (**B**) RVFAC measured using echocardiography in control and Su/Hx rats treated with vehicle, macitentan+tadalafil, MRT1 or MRT1+macientant+tadalafil (n=5 to 10; *p<0.05, ***p<0.001, ****p<0.0001, Kruskal-Wallis’ test followed by Dunn’s post hoc analysis data represent mean ± SEM). (**C**) Representative images, and corresponding quantification of phosphor-FAK (Y397, red) expression in control and Su/Hx rats treated with vehicle or MRT1. PASMCs were labeled with alpha-smooth muscle actin (aSMA, green) (n=5 to 9; ****p<0.0001, one-way ANOVA followed by Dunnett’s post hoc analysis; data represent mean ± SEM). Scale bars: 20 μm. (**D**) Assessment of RV hypertrophy by Fulton index (n=5 to 10; *p<0.05, ****p<0.0001, one-way ANOVA followed by Tukey’s post hoc analysis; data represent mean ± SEM). (**E**) Representative images of RV sections stained with Hematoxylin and eosin (H&E) or Masson’s Trichrome in control and Su/Hx rats treated with vehicle, macitentan+tadalafil, MRT1 or combination of MRT1, macitentan, and tadalafil. The measurement of cardiomyocyte cross-sectional area (CSA) and the quantification of the percentage of collagen area are presented below (n=5 to 10; *p<0.05, **p<0.01, ****p<0.0001, one-way ANOVA followed by Tukey’s post hoc analysis; data represent mean ± SEM). Scale bars: 50 μm (H&E) and 100 μm (Masson’s Trichrome).

**Figure S8.**
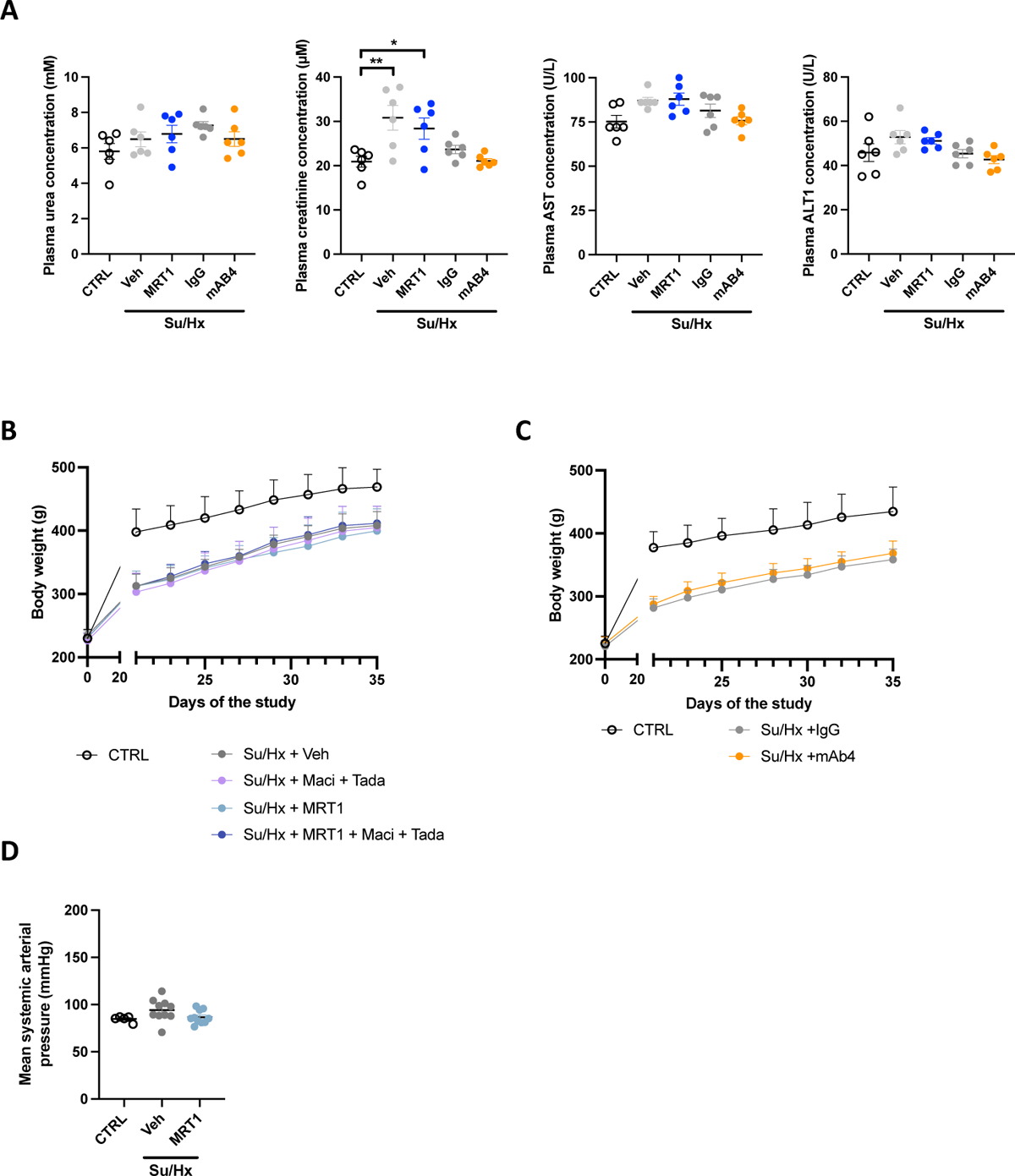
Effects of MRT1 and mAb4 on liver and kidney function, body weight and systemic pressure. **(A)** Plasmatic urea, creatinine, AST, and ALT levels measured at the end of experimental studies (n=6; *p<0.05, **p<0.01, one-way ANOVA followed by Tukey’s post hoc analysis; data rep **(B)** L) an vehicle, macitentan+tadalafil, MRT1 or combination of MRT1, macitentan, and tadalafil **(C)** red during the experimental protocol in control (CTRL) and Sugen/Hypoxia (Su/Hx) rats receiving either mA **). (D)** rol (CTRL) and Sugen/Hypoxia (Su/Hx) rats receiving either vehicle or MRT1 (n=5-10 per groups, data represent mean ± SEM).

**Figure S9.**
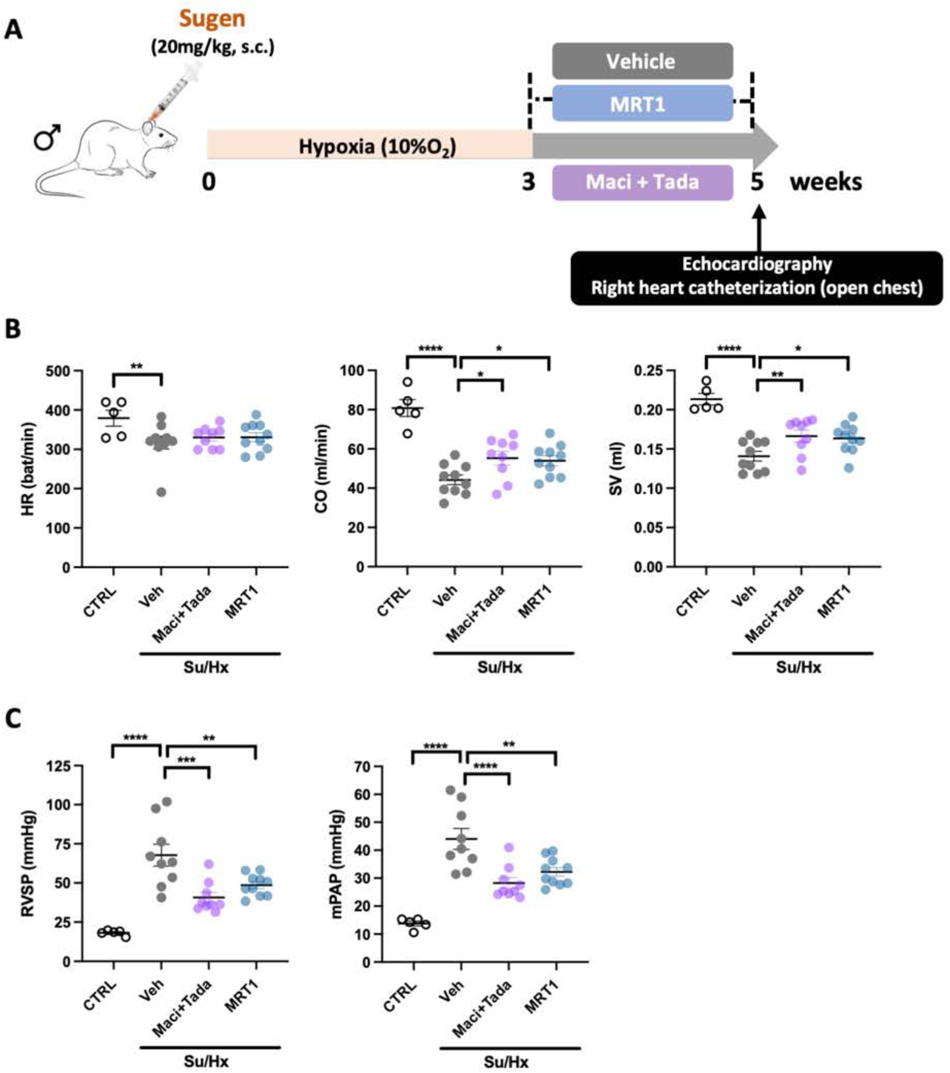
**(related to** Figure 5). External validation of the therapeutic benefits of MRT1 in the Su/Hx rat model. (**A**) Study design using the Sugen/Hypoxia (Su/Hx) rat model. (**B**) HR, CO, and SV measured by echocardiography in control and Su/Hx rats treated with vehicle, macitentan+tadalafil or MRT1 (100 mg/kg twice daily) (n=5 to 10; **p<0.01, **p<0.01, ****p<0.0001, one-way ANOVA followed by Dunnett’s post hoc analysis; data represent mean ± SEM). (**F**) Effect of MRT1 on RVSP and mPAP, as assessed by right heart catheterization with an open chest approach (n=5 to 10; *p<0.05, **p<0.01, ***p<0.001, ****p<0.0001, one-way ANOVA followed by Dunnett’s post hoc analysis; data represent mean ± SEM).

**Figure S10.**
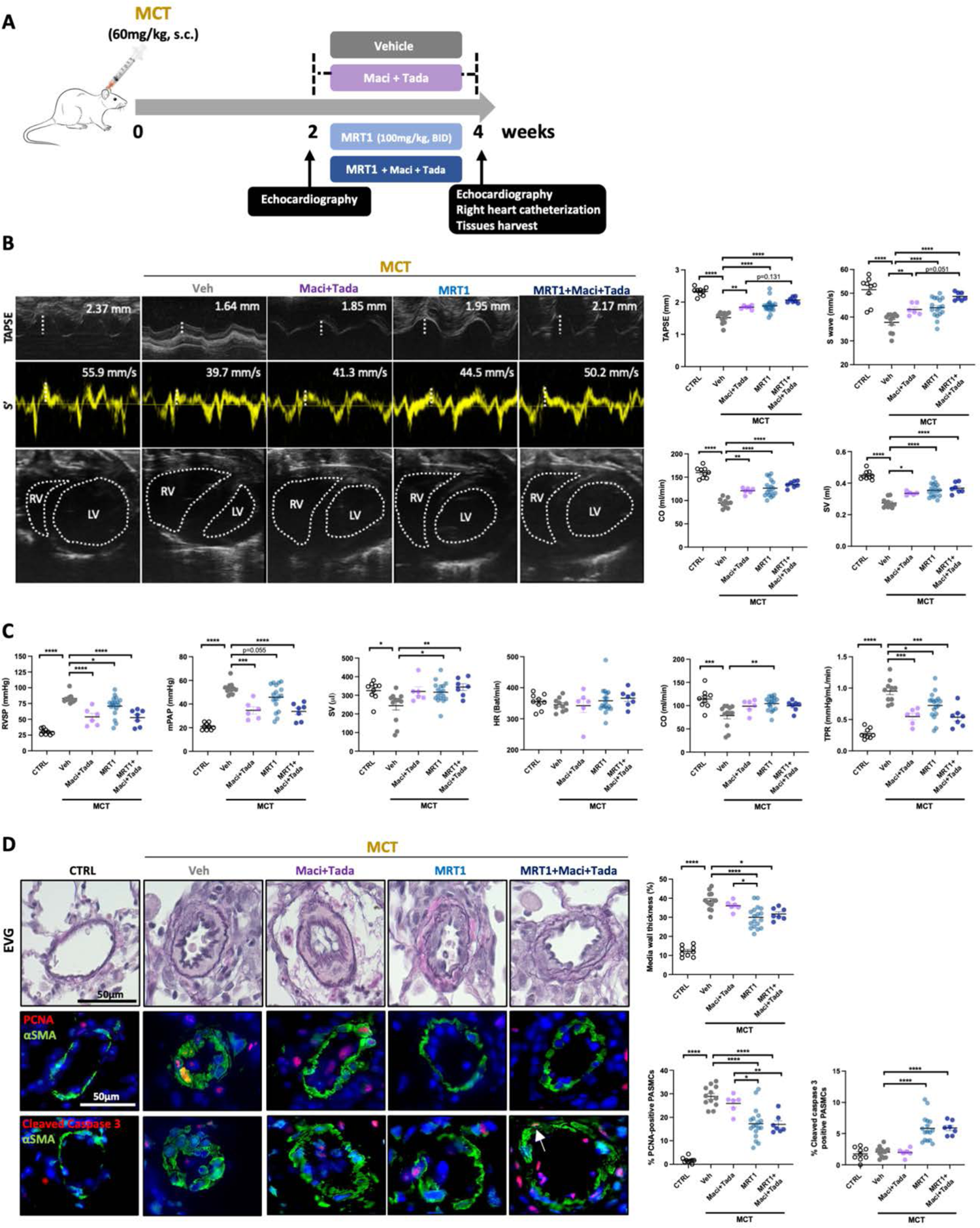
MRT1 alone or combined with standard-of-care drugs improves structural and functional changes in the MCT rat model of PAH. (**A**) Study design using the monocrotaline (MCT) rat model. (**B**) Representative echocardiographic images of TAPSE, S wave (S’), and morphological change of the RV and LV in control and MCT rats treated with vehicle, macitentan+tadalafil, MRT1 or MRT1+macientant+tadalafil. The Quantifications of TAPSE, S wave, CO, and SV are shown (n=9 to 17; *p<0.05, **p<0.01, ***p<0.001, ****p<0.0001, one-way ANOVA followed by Tukey’s post hoc analysis; data represent mean ± SEM). (**C**) Effect of RGD-integrins inhibition on RVSP, mPAP, SV, HR, CO and TPR, as assessed by right heart catheterization (n=9 to 17; *p<0.05, **p<0.01, ***p<0.001, and ****p<0.0001, one-way ANOVA followed by Dunnett’s post hoc analysis for RVSP, mPAP, HR, CO and TPR and Kruskal-Wallis’ test followed by Dunn’s post hoc analysis for SV; data represent mean ± SEM). (**D**) Representative images of distal PAs stained with Elastica van Gieson (EVG) or labeled with PCNA (proliferative marker, red) or cleaved caspase 3 (apoptosis marker, red). Alpha-smooth muscle actin (aSMA, green) was used to detect smooth muscle cells. The quantifications of medial wall thickness, PCNA-positive and cleaved caspase 3-positive smooth muscle cells are presented on the right (n=9 to 18; *p<0.05, **p<0.01, ****p<0.0001, one-way ANOVA followed by Tukey’s post hoc analysis; data represent mean ± SEM). Scale bars, 50 pm.

**Figure S11.**
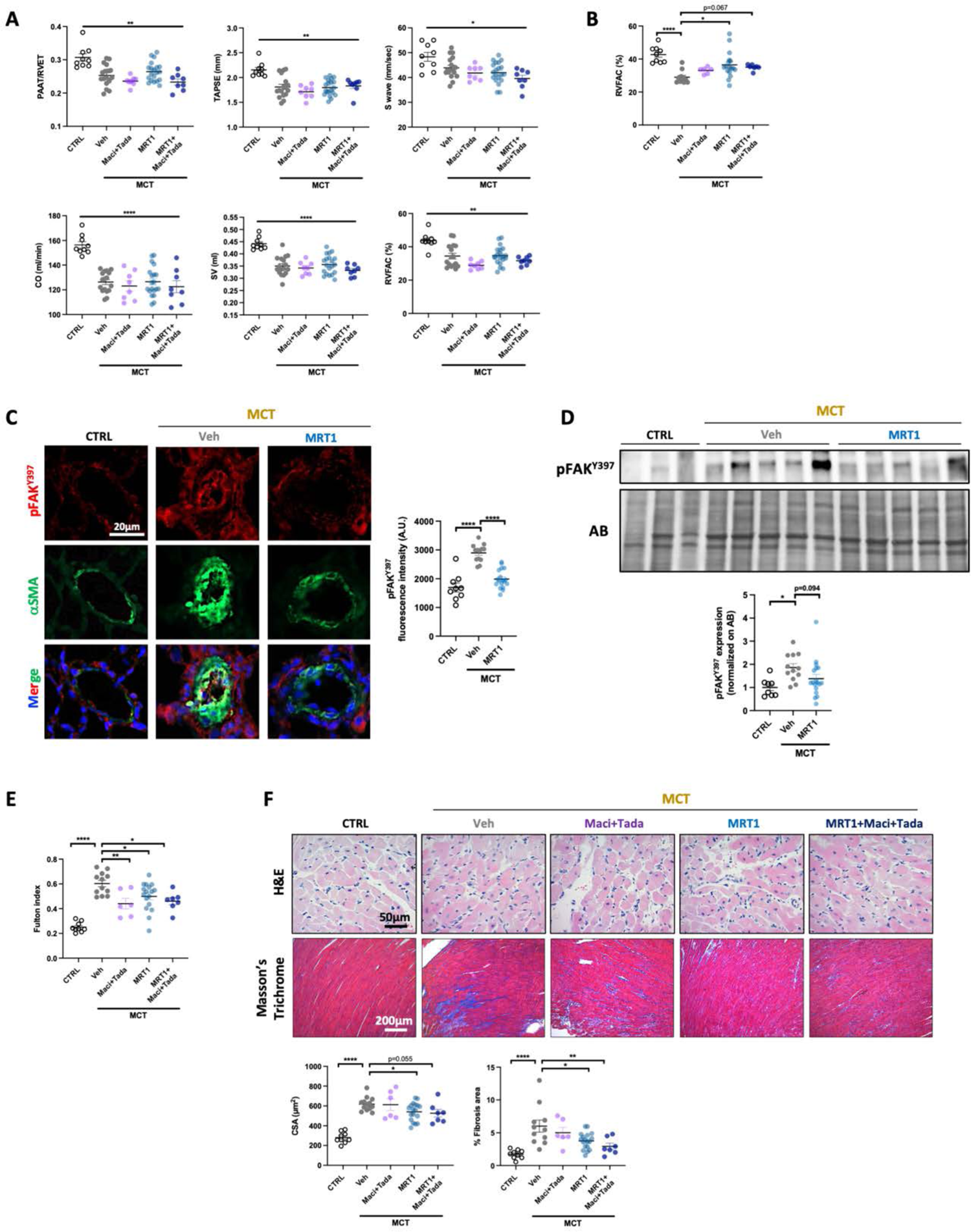
(related to Figure S9). MRT1 alone or combined with standard-of-care drugs improves structural and functional changes in the MCT rat model of PAH. (**A**) PAAT/ET, TAPSE, S wave, CO, SV, and RVFAC measured using echocardiography in control and MCT rats treated with vehicle, macitentan+tadalafil, MRT1 or MRT1+Macientant+tadalafil before treatment initiation (n=9 to 20; *p<0.05, **p<0.01, ***p<0.001, one-way ANOVA followed by Tukey’s post hoc analysis for S wave, CO and SV and Kruskal-Wallis’ test followed by Dunn’s post hoc analysis for PAAT/ET, TAPSE, and RVFAC; data represent mean ± SEM). (**B**) RVFAC measured using echocardiography in control and MCT rats treated with vehicle, macitentan+tadalafil, MRT1 or MRT1+macientant+tadalafil (n=9 to 17; **p<0.01, ****p<0.0001, Kruskal-Wallis’ test followed by Dunn’s post hoc analysis; data represent mean ± SEM). (**C**) Representative images, and corresponding quantification of phospho-FAK (Y397, red) expression in control and MCT rats treated with vehicle or MRT1. PASMCs were labeled with alpha-smooth muscle actin (aSMA, green) (n=9 to 18; ****p<0.0001, one-way ANOVA followed by Dunnett’s post hoc analysis; data represent mean ± SEM). Scale bars: 20 μm. (**D**) Representative Western blot and corresponding densitometric analyses of phospho-FAK (Y397) in dissected PAs from control and MCT rats treated with either MRT1 or its vehicle. Protein expression was normalized to amido black (AB) (n=8 to 18; *p<0.05, Kruskal-Wallis’ test followed by Dunn’s post hoc analysis; data represent mean ± SEM). (**E**) Assessment of RV hypertrophy by Fulton index (n=9 to 18; *p<0.05, **p<0.01, ****p<0.0001, one-way ANOVA followed by Tukey’s post hoc analysis; data represent mean ± SEM). (**F**) Representative images of RV sections stained with hematoxylin and eosin (H&E) or Masson’s Trichrome in control and MCT rats treated with vehicle, MRT1, macitentan+tadalafil, MRT1 or combination of MRT1, macitentan, and tadalafil. The measurement of cardiomyocyte cross-sectional area (CSA) and the quantification of the percentage of collagen area are presented below (n=9 to 18; *p<0.05, **p<0.01, ****p<0.0001, one-way ANOVA followed by Dunnett’s post hoc analysis for CSA or followed by Tukey’s post hoc analysis for Masson’s trichrome; data represent mean ± SEM). Scale bars: 50 μm (H&E) and 100 μm (Masson’s Trichrome).

**Figure S12.**
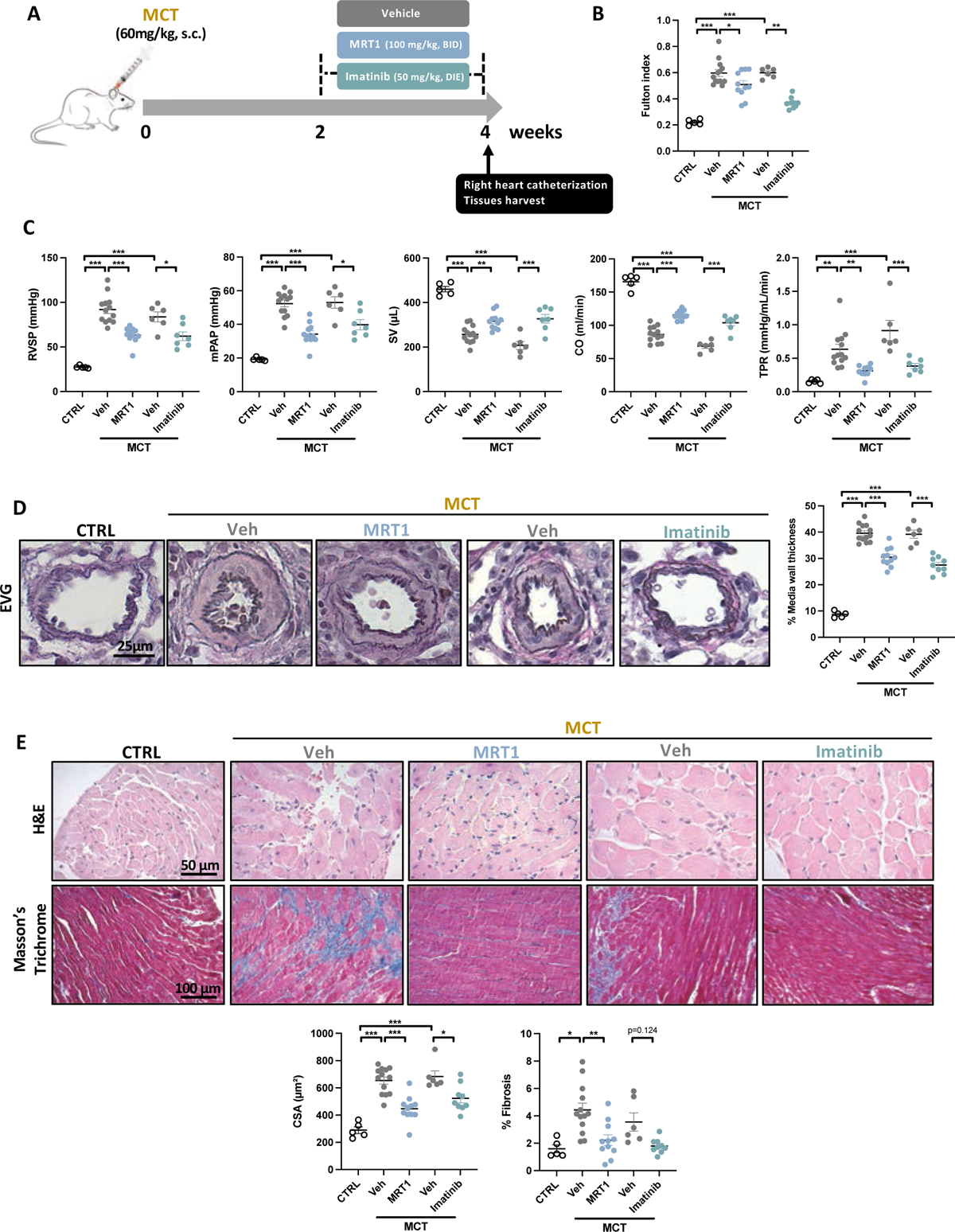
MRT1 and Imatinib demonstrate comparable efficacy in the MCT rat model. (**A**) Study design using the monocrotaline (MCT) rat model. (**B**) Assessment of RV hypertrophy by Fulton index (n=5 to 13; *p<0.05, **p<0.01, ***p<0.001, one-way ANOVA followed by Tukey’s post hoc analysis; data represent mean ± SEM). (**C**) Effect of MRT1 and imatinib on right ventricular systolic pressure (RVSP), mean pulmonary artery pressure (mPAP), stroke volume (SV), cardiac output (CO) and total pulmonary resistance (TPR), as assessed by right heart catheterization (n=5 to 13; *p<0.05, ***p<0.001, one-way ANOVA followed by Tukey’s post hoc analysis; data represent mean ± SEM). (**D**) Representative images of distal PAs stained with Elastica van Gieson (EVG) and quantification of vascular remodeling in control and MCT rats treated or not with MRT1 or imatinib (n=5 to 13; ***p<0.001, one-way ANOVA followed by Tukey’s post hoc analysis; data represent mean ± SEM). Scale bar: 25 μm. (**E**) Representative images of RV sections stained with Hematoxylin and eosin (H&E) or Masson’s Trichrome in control and MCT rats treated or not with MRT1 or imatinib. The measurement of cardiomyocyte cross-sectional area (CSA) and the quantification of the percentage of collagen area are shown (n=5 to 13; *p<0.05, **p<0.01, ****p<0.0001, one-way ANOVA followed by Tukey’s post hoc analysis; data represent mean ± SEM). Scale bars: 50 μm (H&E) and 100 μm (Masson’s Trichrome).

**Figure S13.**
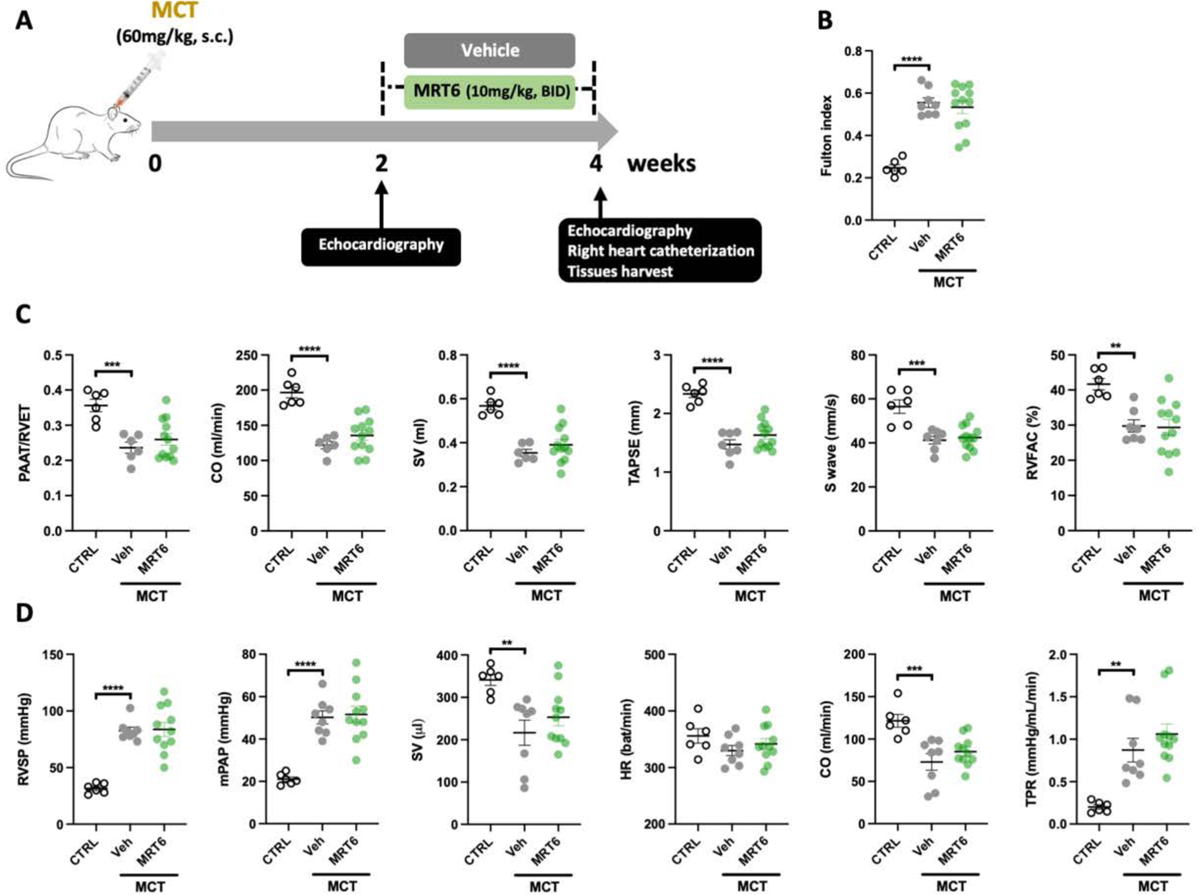
Inhibition of avpi/a8pi integrins using MRT6 did not influence the disease severity in monocrotaline-induced PH rats. (**A**) Study design using the monocrotaline (MCT) rat model. (**B**) Assessment of RV hypertrophy by Fulton index (n=6 to 12; ****p<0.0001, one-way ANOVA followed by Dunnett’s post hoc analysis; data represent mean ± SEM). (**C**) PAAT/ET, CO, SV, TAPSE, S wave and RVFAC measured by echocardiography in control and MCT rats treated or not with MRT6 (n=6 to 12; **p<0.01, ***p<0.001, ****p<0.0001, one-way ANOVA followed by Dunnett’s post hoc analysis; data represent mean ± SEM). (**D**) Effect of MRT6 on RVSP, mPAP, SV, HR, CO and TPR, as assessed by right heart catheterization (n=6 to 11; **p<0.01, ***p<0.001, ****p<0.0001, one-way ANOVA followed by Dunnett’s post hoc analysis for RVSP, mPAP, SV, HR and CO and Kruskal-Wallis’ test followed by Dunn’s post hoc analysis for TPR; data represent mean ± SEM).

**Figure S14.**
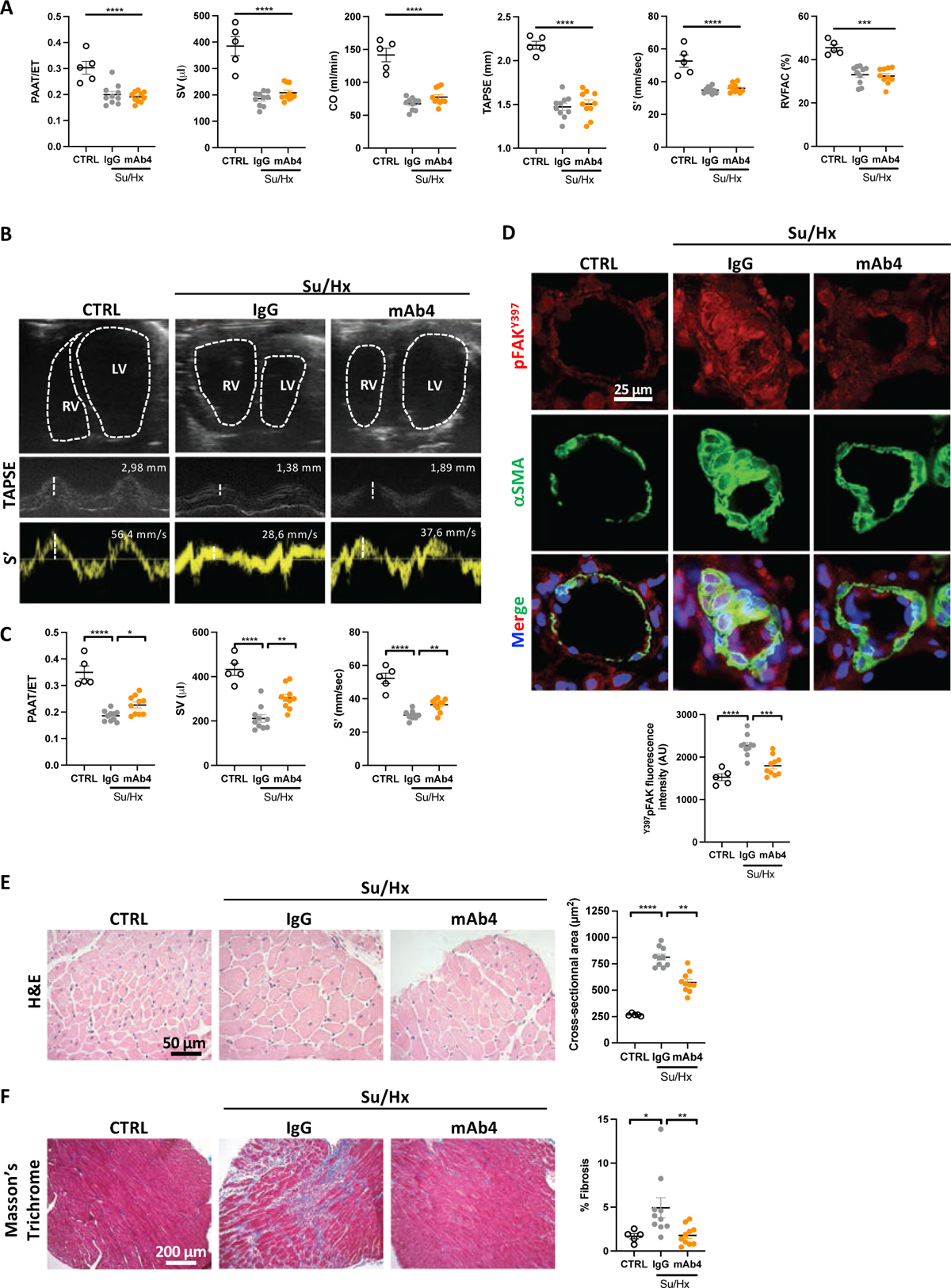
**(related to** Figure 6). A function-blocking anti-rat integrin a5pi *antibody* improves established PAH in Sugen/Hypoxia (Su/Hx)-challenged rats. (**A**) PAAT/ET, SV, CO, TAPSE, S’, and RVFAC measured using echocardiography in control and Su/Hx rats before treatment initiation (n=5 to 10; ****p<0.0001, one-way ANOVA followed by Dunnett’s post hoc analysis for PAAT/ET, CO, TAPSE, S’, and RVFAC and Kruskal-Wallis’ test followed by Dunn’s post hoc analysis for SV; data represent mean ± SEM). (**B**) Representative echocardiographic images of morphological change of the RV and LV, tricuspid annular plane systolic excursion (TAPSE) and S wave (S’) in control and Su/Hx rats treated with IgG or mAb4. (**C**) PAAT/ET, SV, and S’ measured using echocardiography in control and Su/Hx rats treated with IgG or mAb4 (n=5 to 10; *p<0.05, **p<0.01, ****p<0.0001, one-way ANOVA followed by Dunnett’s post hoc analysis for PAAT/ET and S’, Kruskal-Wallis’ test followed by Dunn’s post hoc analysis for SV; data represent mean ± SEM). (**D**) Representative images, and corresponding quantification of phospho-FAK (Y397, red) expression in control, Su/Hx+IgG, and Su/Hx+mAb4 rats. PASMCs were labeled with alpha-smooth muscle actin (aSMA, green) (n=5 to 10; **p<0.01, one-way ANOVA followed by Dunnett’s post hoc analysis; data represent mean ± SEM). Scale bar: 25 μm. (**E**) Representative images of Hematoxylin and eosin (H&E) stained RV sections and measurement of cardiomyocyte cross-sectional area (CSA) in control and Su/Hx rats treated with either mAb4 or IgG (n=5 to 10; **p<0.01, ****p<0.0001, one-way ANOVA followed by Dunnett’s post hoc analysis; data represent mean ± SEM). Scale bar: 50 μm. (**F**) Representative images of Masson’s Trichrome stained RV sections and quantification of the percentage of collagen area in control and Su/Hx rats treated with either mAb4 or IgG n=5 to 10; *p<0.05, **p<0.01, ****p<0.0001 Kruskal-Wallis’ test followed by Dunn’s post hoc analysis; data represent mean ± SEM). Scale bar: 200 pm.

**Figure S15.**
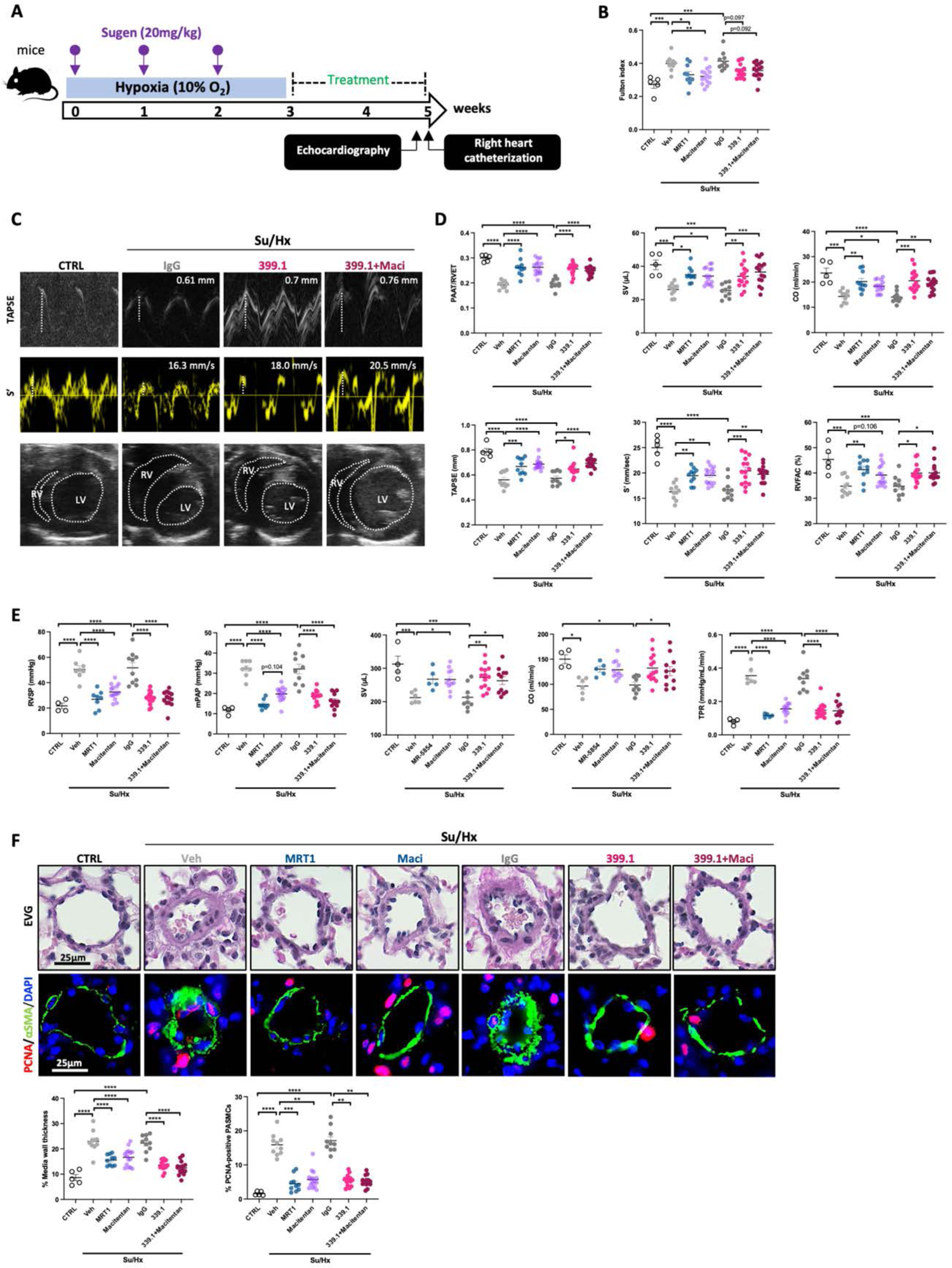
A function-blocking anti-mouse integrin a5pi *antibody* reverses PAH in Su/Hx-challenged mice. (**A**) Study design using the Sugen/Hypoxia (Su/Hx) mouse model. (**B**) Assessment of RV hypertrophy by Fulton index (n=5 to 15; *p<0.05, **p<0.01, ***p<0.001, ****p<0.0001, one-way ANOVA followed by Tukey’s post hoc analysis; data represent mean ± SEM). (**C**) Representative echocardiographic images of tricuspid annular plane systolic excursion (TAPSE), S wave (S’), and morphological change of the RV and LV in control and Su/Hx mice treated with IgG, 339.1 or combination of 339.1 and macitentan. (**D**) PAAT/ET, SV, CO, TAPSE, S’, and RV FAC measured using echocardiography in control and Su/Hx mice treated with vehicle, MRT1, macitentan, IgG, 339.1 or combination of 339.1 and macitentan (n=5 to 15; *p<0.05, **p<0.01, ***p<0.001, ****p<0.0001, one-way ANOVA followed by Tukey’s post hoc analysis; data represent mean ± SEM). (**E**) Effect of integrin a5pi inhibition on RVSP, mPAP, SV, CO and TPR, as assessed by right heart catheterization (n=4 to 14; *p<0.05, **p<0.01, ***p<0.001, ****p<0.0001, one-way ANOVA followed by Tukey’s post hoc analysis; data represent mean ± SEM). (**F**) Representative images of distal PAs stained with Elastica van Gieson (EVG) or co-labeled with PCNA (proliferative marker, red) and alpha-smooth muscle actin (aSMA, green). The quantifications of medial wall thickness and percentage of PCNA-positive smooth muscle cells are presented below (n=5 to 15; *p<0.05, **p<0.01, ***p<0.001, ****p<0.0001, one-way ANOVA followed by Tukey’s post hoc analysis for EVG and Kruskal-Wallis’ test followed by Dunn’s post hoc analysis for PCNA; data represent mean ± SEM). Scale bars, 25 pm.

**Figure S16.**
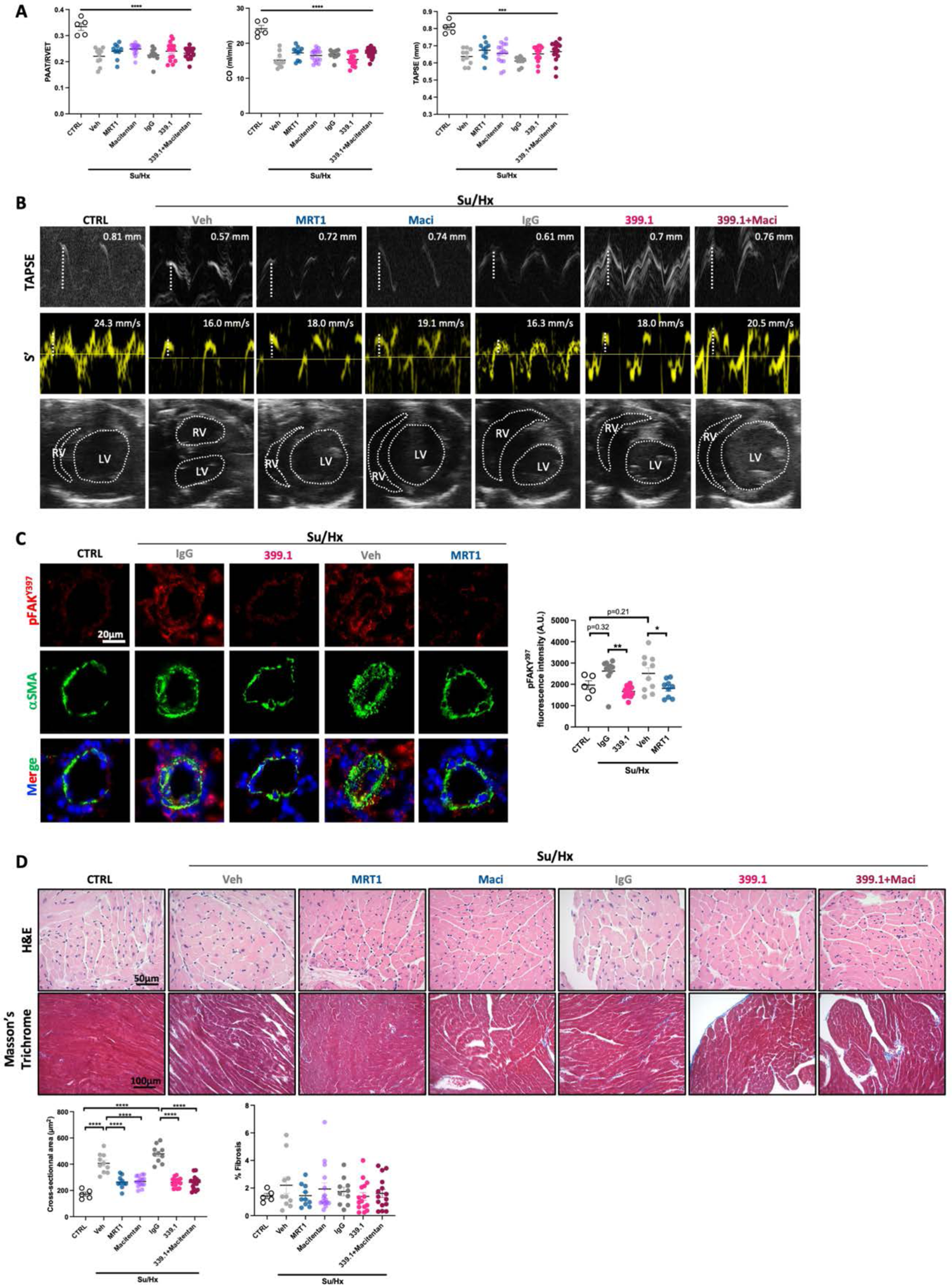
(related to Figure S15). A function-blocking anti-mouse integrin a5pi *antibody* reverses PAH in Sugen/Hypoxia (Su/Hx)-challenged mice. (**A**) PAAT/ET, CO, and TAPSE measured using echocardiography in control and Su/Hx mice before treatment initiation (n=5 to 15; ***p<0.001, one-way ANOVA followed by Tukey’s analysis; data represent mean ± SEM). (**B**) Representative echocardiographic images of tricuspid annular plane systolic excursion (TAPSE), S wave (S’), and the morphological change of RV and LV in control and Su/Hx mice treated with vehicle, MRT1, macitentan, IgG, 339.1 or combination of 339.1 and macitentan. (**C**) Representative images, and corresponding quantification of phospho-FAK (Y397, red) expression in control, Su/Hx+IgG, Su/Hx+339.1, Su/Hx+vehicle, and Su/Hx+MRT1 mice. PASMCs were labeled with alpha-smooth muscle actin (aSMA, green) (n=5 to 15; *p<0.05, one-way ANOVA followed by Tukey’s post hoc analysis; data represent mean ± SEM). Scale bar: 20 μm. (**D**) Representative images of RV sections stained with Hematoxylin and eosin (H&E) or Masson’s Trichrome in control and Su/Hx mice treated with vehicle, MRT1, macitentan, IgG, 339.1 or combination of 339.1 and macitentan. The measurement of cardiomyocyte cross-sectional area (CSA) and the quantification of the percentage of collagen area are presented below (n=5 to 10; ****p<0.0001. one-way ANOVA followed by Tukey’s analysis for H&E and Kruskal-Wallis’ test followed by Dunn’s post hoc analysis for Masson’s trichrome; data represent mean ± SEM). Scale bars: 50 μm (H&E) and 100 μm (Masson’s Trichrome).

## SUPPLEMENTARY TABLES

**Table S1.**
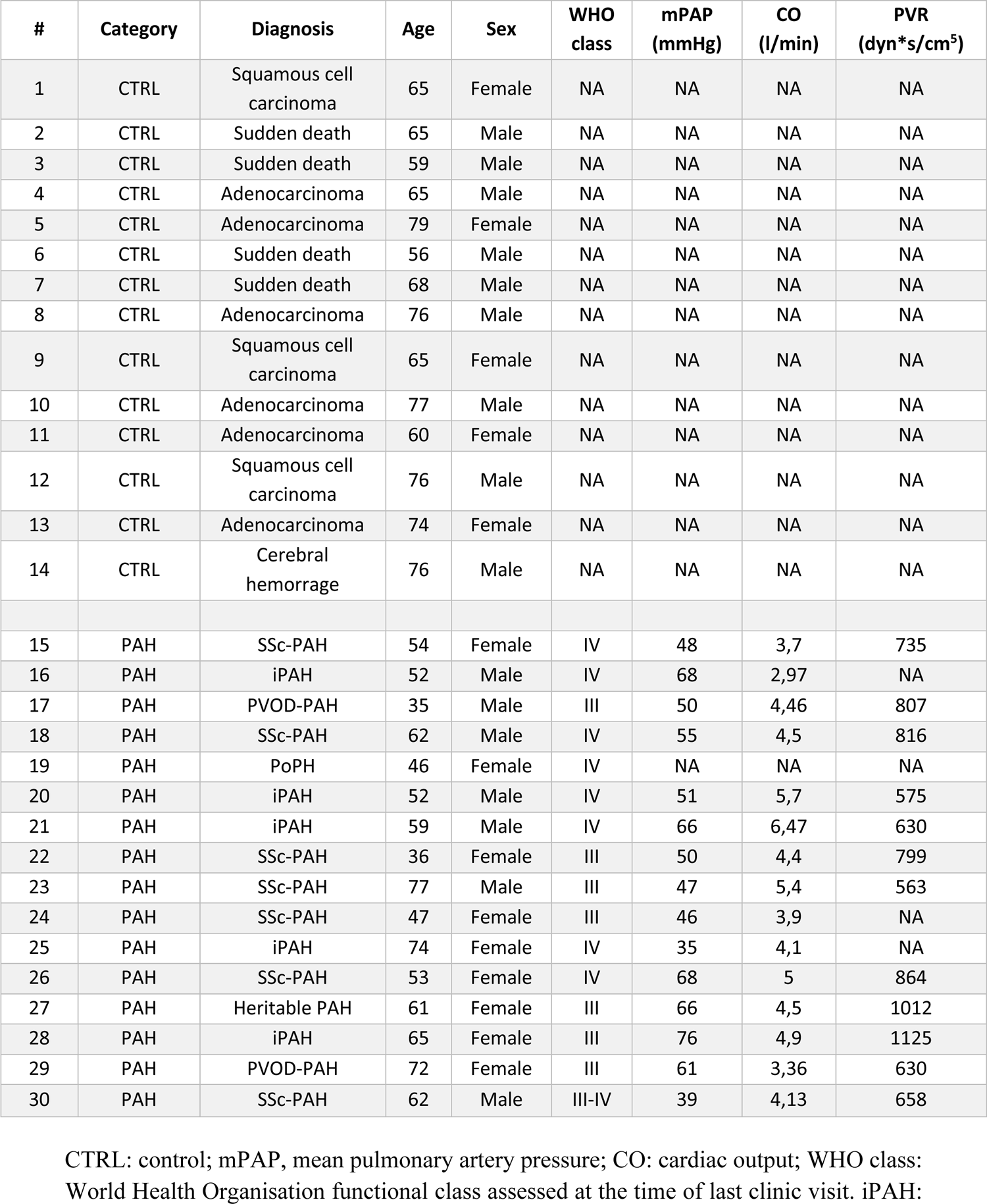
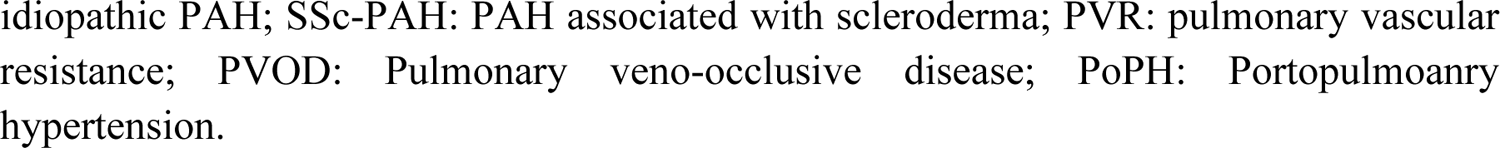
Detailed characteristics of human pulmonary arteries used in this study.

| # | Category | Diagnosis | Age | Sex | WHO class | mPAP (mmHg) | CO (l/min) | PVR (dyn*s/cm<sup>5</sup>) |
| --- | --- | --- | --- | --- | --- | --- | --- | --- |
| 1 | CTRL | Squamous cell carcinoma | 65 | Female | NA | NA | NA | NA |
| 2 | CTRL | Sudden death | 65 | Male | NA | NA | NA | NA |
| 3 | CTRL | Sudden death | 59 | Male | NA | NA | NA | NA |
| 4 | CTRL | Adenocarcinoma | 65 | Male | NA | NA | NA | NA |
| 5 | CTRL | Adenocarcinoma | 79 | Female | NA | NA | NA | NA |
| 6 | CTRL | Sudden death | 56 | Male | NA | NA | NA | NA |
| 7 | CTRL | Sudden death | 68 | Male | NA | NA | NA | NA |
| 8 | CTRL | Adenocarcinoma | 76 | Male | NA | NA | NA | NA |
| 9 | CTRL | Squamous cell carcinoma | 65 | Female | NA | NA | NA | NA |
| 10 | CTRL | Adenocarcinoma | 77 | Male | NA | NA | NA | NA |
| 11 | CTRL | Adenocarcinoma | 60 | Female | NA | NA | NA | NA |
| 12 | CTRL | Squamous cell carcinoma | 76 | Male | NA | NA | NA | NA |
| 13 | CTRL | Adenocarcinoma | 74 | Female | NA | NA | NA | NA |
| 14 | CTRL | Cerebral hemorrhage | 76 | Male | NA | NA | NA | NA |
| 15 | PAH | SSc-PAH | 54 | Female | IV | 48 | 3,7 | 735 |
| 16 | PAH | iPAH | 52 | Male | IV | 68 | 2,97 | NA |
| 17 | PAH | PVOD-PAH | 35 | Male | III | 50 | 4,46 | 807 |
| 18 | PAH | SSc-PAH | 62 | Male | IV | 55 | 4,5 | 816 |
| 19 | PAH | PoPH | 46 | Female | IV | NA | NA | NA |
| 20 | PAH | iPAH | 52 | Male | IV | 51 | 5,7 | 575 |
| 21 | PAH | iPAH | 59 | Male | IV | 66 | 6,47 | 630 |
| 22 | PAH | SSc-PAH | 36 | Female | III | 50 | 4,4 | 799 |
| 23 | PAH | SSc-PAH | 77 | Male | III | 47 | 5,4 | 563 |
| 24 | PAH | SSc-PAH | 47 | Female | III | 46 | 3,9 | NA |
| 25 | PAH | iPAH | 74 | Female | IV | 35 | 4,1 | NA |
| 26 | PAH | SSc-PAH | 53 | Female | IV | 68 | 5 | 864 |
| 27 | PAH | Heritable PAH | 61 | Female | III | 66 | 4,5 | 1012 |
| 28 | PAH | iPAH | 65 | Female | III | 76 | 4,9 | 1125 |
| 29 | PAH | PVOD-PAH | 72 | Female | III | 61 | 3,36 | 630 |
| 30 | PAH | SSc-PAH | 62 | Male | III-IV | 39 | 4,13 | 658 |

CTRL: control; mPAP, mean pulmonary artery pressure; CO: cardiac output; WHO class: World Health Organisation functional class assessed at the time of last clinic visit. iPAH:
idiopathic PAH; SSc-PAH: PAH associated with scleroderma; PVR: pulmonary vascular resistance; PVOD: Pulmonary veno-occlusive disease; PoPH: Portopulmonary hypertension.

**Table S2.** Detailed characteristics of human PASMCs used in this study.

| # | Category | Diagnosis | Age | Sex | WHO class | mPAP (mmHg) | CO (l/min) | PVR (dyn*s/cm<sup>5</sup>) |
| --- | --- | --- | --- | --- | --- | --- | --- | --- |
| 1 | CTRL | Normal | 35 | Female | NA | NA | NA | NA |
| 2 | CTRL | Normal | 56 | Female | NA | NA | NA | NA |
| 3 | CTRL | Normal | 53 | Male | NA | NA | NA | NA |
| 4 | CTRL | Normal | 45 | Male | NA | NA | NA | NA |
| 5 | CTRL | Normal | 50 | Female | NA | NA | NA | NA |
| 6 | CTRL | Normal | 40 | Male | NA | NA | NA | NA |
| 7 | PAH | iPAH | 39 | Male | Unknown | Unknown | Unknown | Unknown |
| 8 | PAH | iPAH | 45 | Male | Unknown | Unknown | Unknown | Unknown |
| 9 | PAH | SSc-PAH | 45 | Female | IV | 43 | 7,25 | 375 |
| 10 | PAH | iPAH | 65 | Male | III | 76 | 4,9 | 1125 |
| 11 | PAH | iPAH | 52 | Male | IV | 68 | 2,97 | Unknown |
| 12 | PAH | Heritable PAH | 61 | Female | III | 66 | 4,5 | 1012 |
| 13 | PAH | iPAH | 23 | Female | IV | Unknown | Unknown | 720 |
| 14 | PAH | Heritable PAH | 35 | Female | IV | 60 | 2,1 | Unknown |
| 15 | PAH | SSc-PAH | 62 | Male | IV | 55 | 4,5 | 816 |

CTRL: control; mPAP, mean pulmonary artery pressure; CO: cardiac output; WHO class: World Health Organisation functional class assessed at the time of last clinic visit. iPAH: idiopathic PAH; SSc-PAH: PAH associated with scleroderma; PVR: pulmonary vascular resistance.

**Table S3.** Detailed characteristics of human PAECs used in this study.

| # | Category | Diagnosis | Age | Sex | WHO class | mPAP (mmHg) | CO (l/min) | PVR (dyn*s/cm<sup>5</sup>) |
| --- | --- | --- | --- | --- | --- | --- | --- | --- |
| 1 | CTRL | Normal | 21 | Male | NA | NA | NA | NA |
| 2 | CTRL | Normal | 56 | Female | NA | NA | NA | NA |
| 3 | CTRL | Normal | 22 | Male | NA | NA | NA | NA |
| 4 | CTRL | Normal | 48 | Male | NA | NA | NA | NA |
| 5 | PAH | Heritable PAH | 40 | Female | III | 56 | 3,79 | Unknown |
| 6 | PAH | PVOD-PAH | 35 | Male | III | 50 | 4,46 | 807 |
| 7 | PAH | PVOD-PAH | 72 | Female | III | 61 | 3,36 | 630 |
| 8 | PAH | SSc-PAH | 45 | Female | IV | 43 | 7,25 | 375 |
| 9 | PAH | iPAH | 52 | Male | IV | 68 | 2,97 | Unknown |
| 10 | PAH | Heritable PAH | 61 | Female | III | 66 | 4,5 | 1012 |
| 11 | PAH | Heritable PAH | 32 | Female | IV | 56 | 5,9 | Unknown |
| 12 | PAH | Heritable PAH | 35 | Female | IV | 60 | 2,1 | Unknown |
| 13 | PAH | Heritable PAH | 26 | Female | IV | 99 | Unknown | Unknown |

CTRL: control; mPAP, mean pulmonary artery pressure; CO: cardiac output; WHO class: World Health Organisation functional class assessed at the time of last clinic visit. iPAH: idiopathic PAH; SSc-PAH: PAH associated with scleroderma; PVR: pulmonary vascular resistance; PVOD: Pulmonary veno-occlusive disease.

**Table S4.** Detailed characteristics of human lung tissues used in this study for the PCLS.

| # | Category | Diagnosis | Age | Sex | mPAP (mmHg) | CO (l/min) |
| --- | --- | --- | --- | --- | --- | --- |
| 1 | CTRL | Tumor resection | 69 | Female | NA | NA |
| 2 | CTRL | Tumor resection | 69 | Male | NA | NA |
| 3 | CTRL | Tumor resection | 67 | Female | NA | NA |
| 4 | CTRL | Tumor resection | 70 | Female | NA | NA |
| 5 | CTRL | Tumor resection | 64 | Female | NA | NA |
| 6 | PAH | SSc-PAH | 67 | Female | 43 | 3,07 |
| 7 | PAH | iPAH | 82 | Male | 47 | 5,20 |
| 8 | PAH | Heritable | 39 | Female | NA | 2,31 |
| 9 | IPF | IPF-PH | 68 | Male | 29* | 4,84 |
| 10 | IPF | IPF-PH | 65 | Male | 55** | NA |

CTRL: control; mPAP, mean pulmonary artery pressure; CO: cardiac output; SSc-PAH: PAH associated with scleroderma; iPAH: idiopathic PAH; IPF: idiopathic pulmonary fibrosis; IPF+PH: pulmonary hypertension associated with IPF
\* mPAP was measured by right heart catheterization
\*\* mPAP was estimated from echocardiography

**Table S5.**
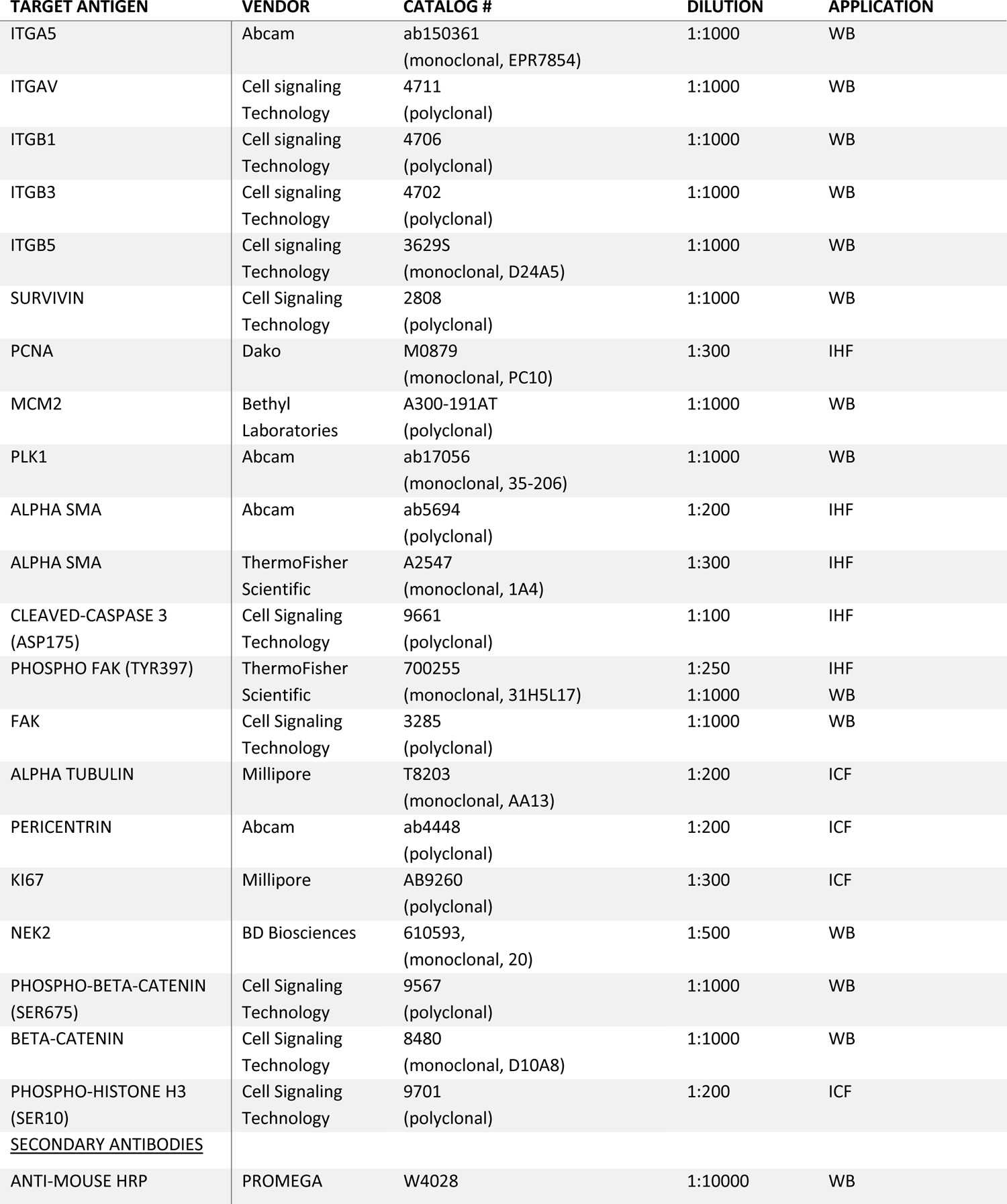
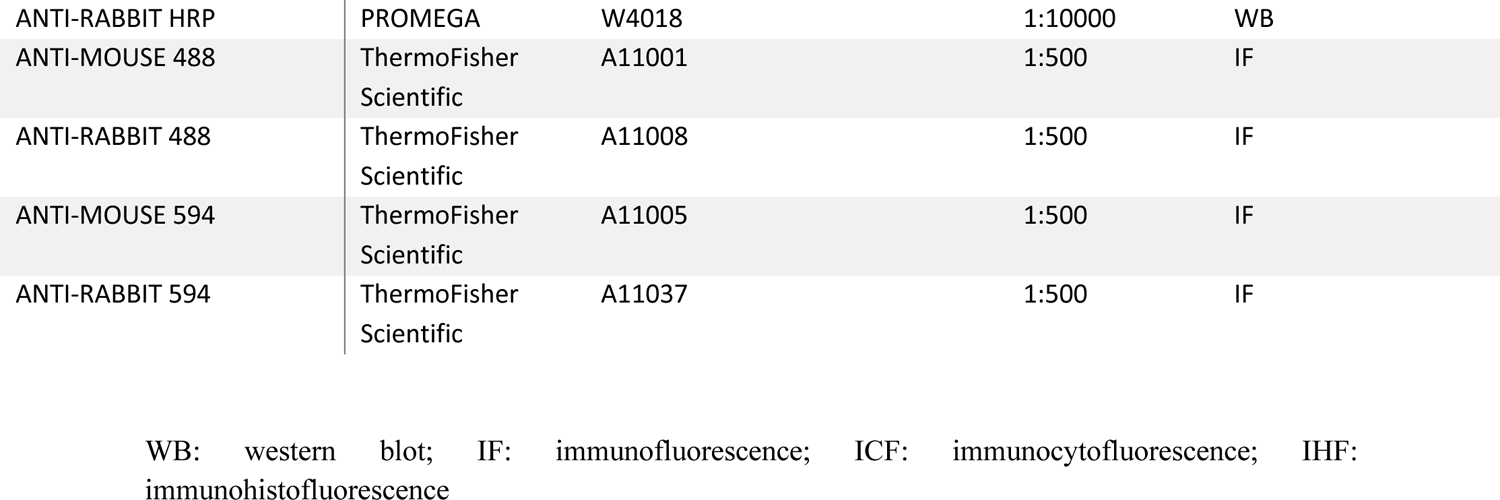
Antibodies used for Western blotting and immunofluorescence analysis.

**Table S6.**
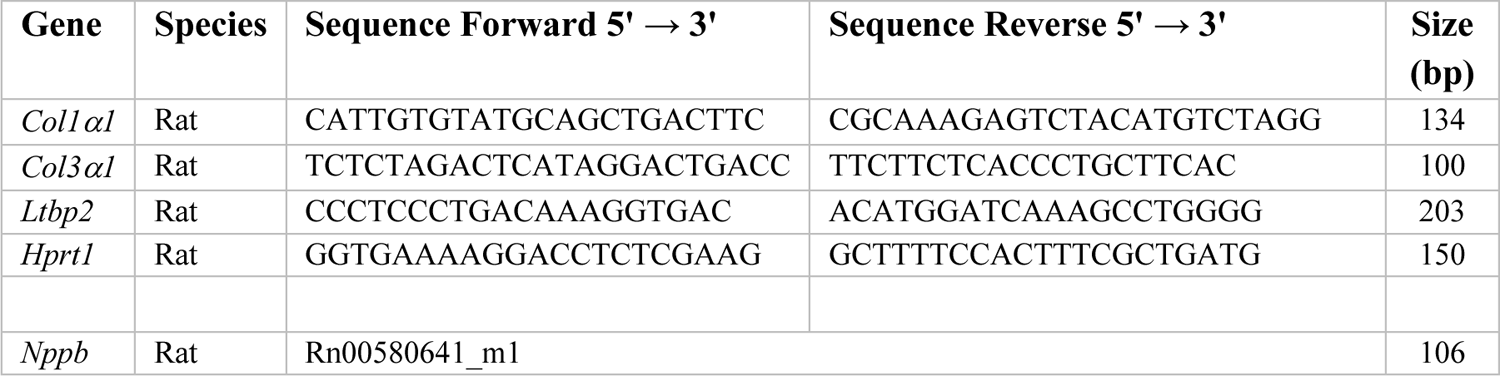
Primer sequences used for gene expression analysis.

**Table S7.**
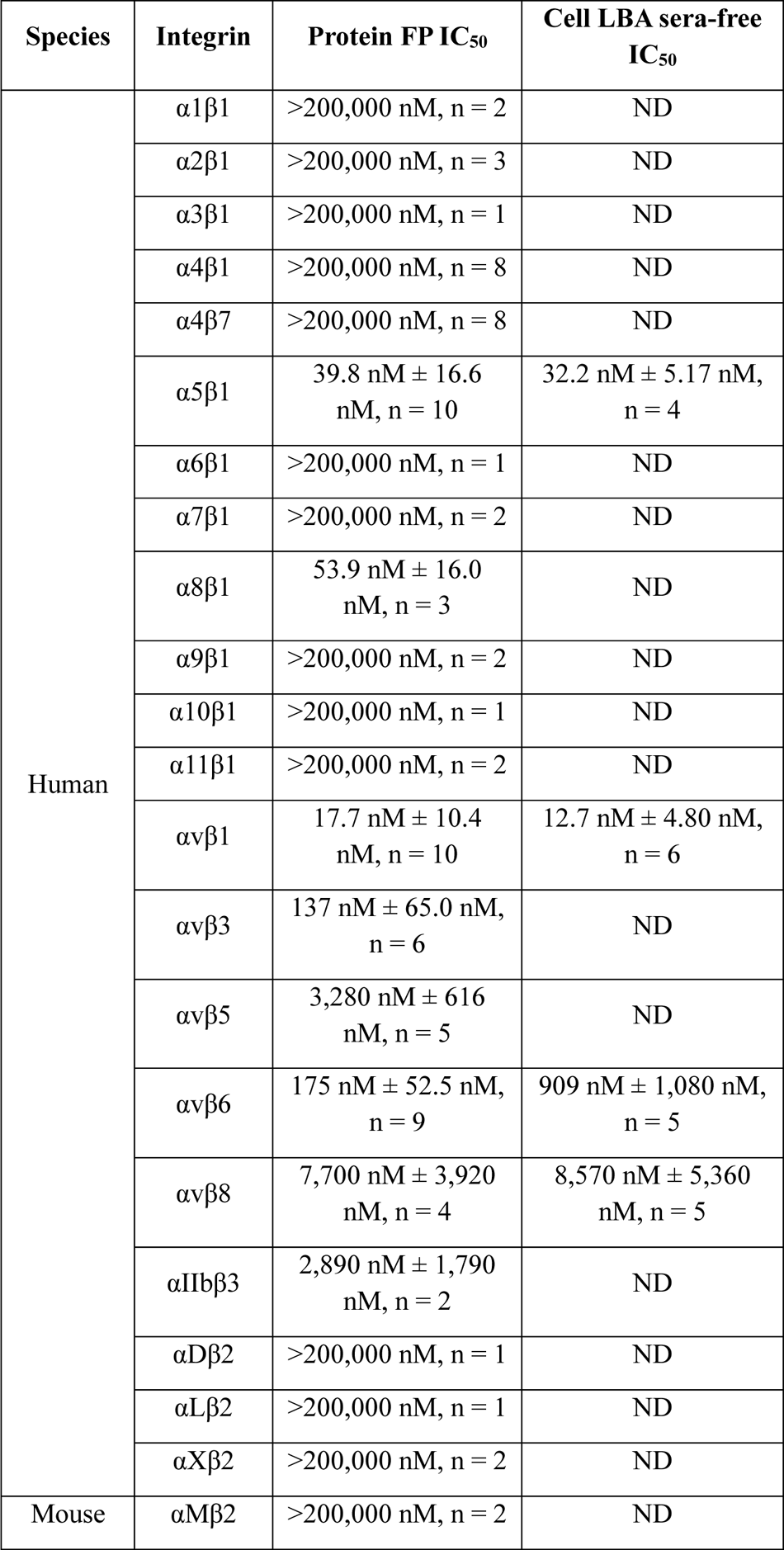
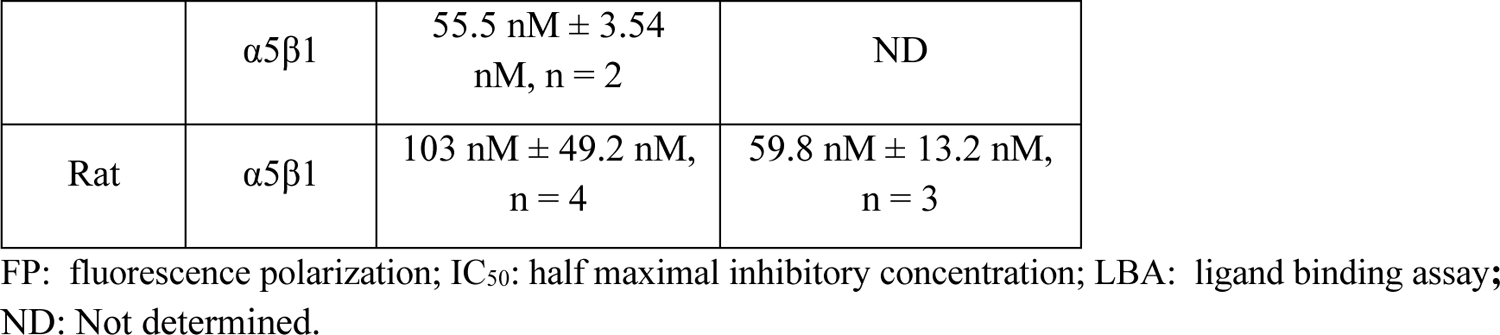
Activity of MRT1 (geomean ± standard deviation, number of assays)

| Species | Integrin | Protein FP IC<sub>50</sub> | Cell LBA sera-free IC<sub>50</sub> |
| --- | --- | --- | --- |
| Human | $\alpha 1\beta 1$ | >200,000 nM, n = 2 | ND |
| | $\alpha 2\beta 1$ | >200,000 nM, n = 3 | ND |
| | $\alpha 3\beta 1$ | >200,000 nM, n = 1 | ND |
| | $\alpha 4\beta 1$ | >200,000 nM, n = 8 | ND |
| | $\alpha 4\beta 7$ | >200,000 nM, n = 8 | ND |
| | $\alpha 5\beta 1$ | 39.8 nM ± 16.6 nM, n = 10 | 32.2 nM ± 5.17 nM, n = 4 |
| | $\alpha 6\beta 1$ | >200,000 nM, n = 1 | ND |
| | $\alpha 7\beta 1$ | >200,000 nM, n = 2 | ND |
| | $\alpha 8\beta 1$ | 53.9 nM ± 16.0 nM, n = 3 | ND |
| | $\alpha 9\beta 1$ | >200,000 nM, n = 2 | ND |
| | $\alpha 10\beta 1$ | >200,000 nM, n = 1 | ND |
| | $\alpha 11\beta 1$ | >200,000 nM, n = 2 | ND |
| | $\alpha v\beta 1$ | 17.7 nM ± 10.4 nM, n = 10 | 12.7 nM ± 4.80 nM, n = 6 |
| | $\alpha v\beta 3$ | 137 nM ± 65.0 nM, n = 6 | ND |
| | $\alpha v\beta 5$ | 3,280 nM ± 616 nM, n = 5 | ND |
| | $\alpha v\beta 6$ | 175 nM ± 52.5 nM, n = 9 | 909 nM ± 1,080 nM, n = 5 |
| | $\alpha v\beta 8$ | 7,700 nM ± 3,920 nM, n = 4 | 8,570 nM ± 5,360 nM, n = 5 |
| | $\alpha IIb\beta 3$ | 2,890 nM ± 1,790 nM, n = 2 | ND |
| | $\alpha D\beta 2$ | >200,000 nM, n = 1 | ND |
| | $\alpha L\beta 2$ | >200,000 nM, n = 1 | ND |
| | $\alpha X\beta 2$ | >200,000 nM, n = 2 | ND |
| Mouse | $\alpha M\beta 2$ | >200,000 nM, n = 2 | ND |
| | $\alpha 5\beta 1$ | 55.5 nM ± 3.54 nM, n = 2 | ND |
| Rat | $\alpha 5\beta 1$ | 103 nM ± 49.2 nM, n = 4 | 59.8 nM ± 13.2 nM, n = 3 |

FP: fluorescence polarization; IC<sub>50</sub>: half maximal inhibitory concentration; LBA: ligand binding assay; ND: Not determined.

**Table S8.**
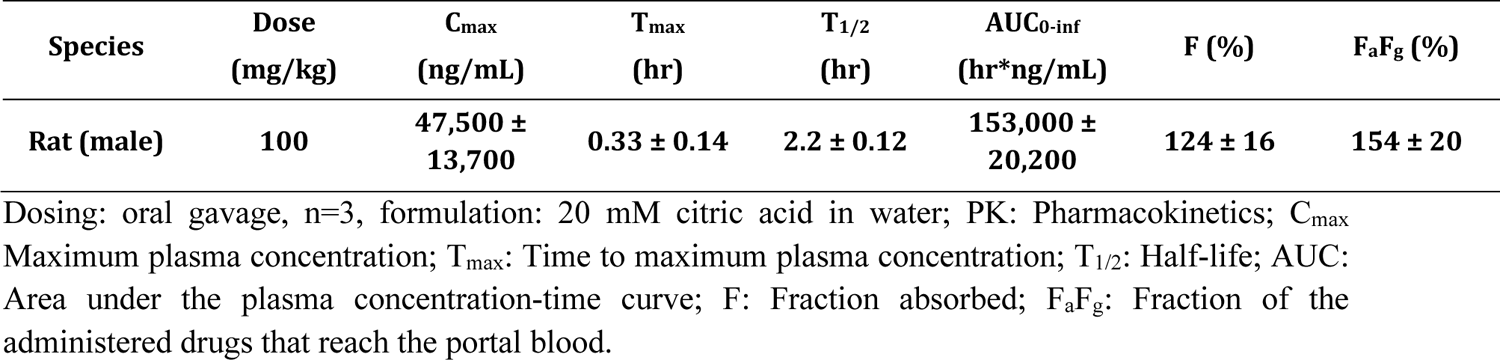
Preclinical oral PK of MRT1.

**Table S9.** MRT1 and mAb4 plasma and serum concentrations in the efficacy studies.

| Study Type | Test article | Dose* (mg/kg) | Terminal plasma/Serum concentration ± (ng/mL) |
| --- | --- | --- | --- |
| MCT | MRT1 | 100 | 1,410 ± 2,010 |
| SuHx | MRT1 | 100 | 151 ± 158 |
| | MRT1 + SoC | 100 | 401 ± 445 |
| SuHx | mAb4 | 10 | 177,000 ± 47,800<br>(1,180 ± 319 nM) |

\*Doses were selected to ensure IC<sub>90</sub> target coverage. MCT: monocrotaline; SoC: standard of care macitentan + tadalafil; Su/Hx: Sugen/Hypoxia

**Table S10.**
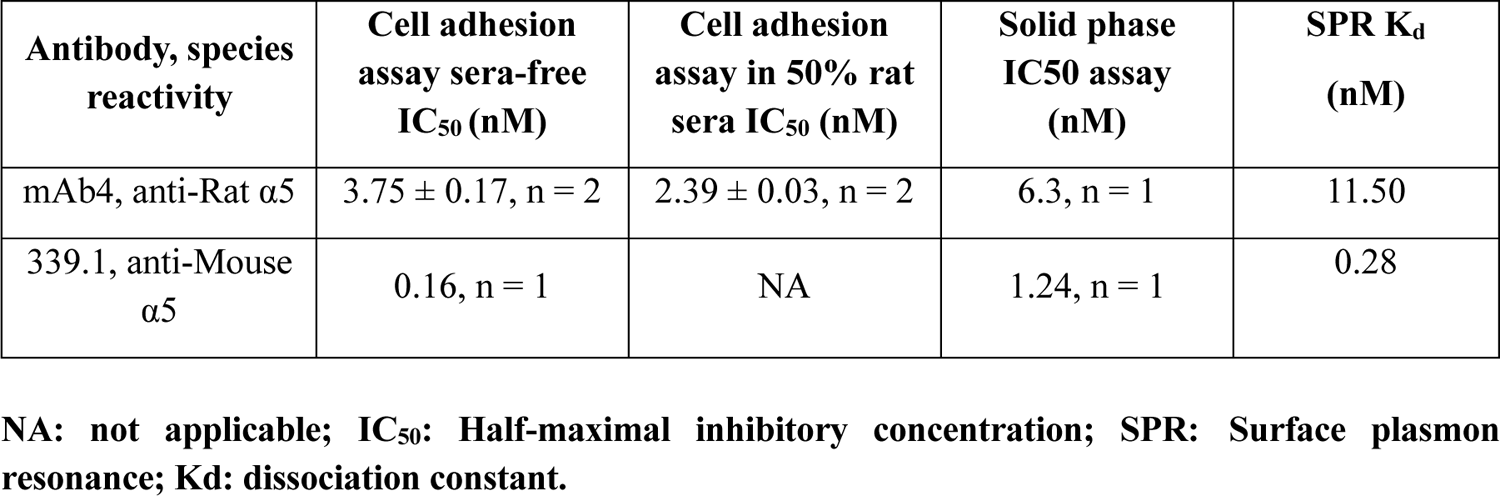
Activity of mAb4 and 339.1 antibodies in assays against rat and mouse α5β1, respectively (geomean ± standard deviation, number of assays)

| Antibody, species reactivity | Cell adhesion assay sera-free IC<sub>50</sub> (nM) | Cell adhesion assay in 50% rat sera IC<sub>50</sub> (nM) | Solid phase IC<sub>50</sub> assay (nM) | SPR K<sub>d</sub> (nM) |
| --- | --- | --- | --- | --- |
| mAb4, anti-Rat $\alpha 5$ | 3.75 ± 0.17, n = 2 | 2.39 ± 0.03, n = 2 | 6.3, n = 1 | 11.50 |
| 339.1, anti-Mouse $\alpha 5$ | 0.16, n = 1 | NA | 1.24, n = 1 | 0.28 |

NA: not applicable; IC<sub>50</sub>: Half-maximal inhibitory concentration; SPR: Surface plasmon resonance; K<sub>d</sub>: dissociation constant.

**Table S11.** Preclinical SC PK of mAb4.

| Species | Dose (mg/kg) | C<sub>max</sub> (ng/mL) | T<sub>max</sub> (hr) | T<sub>1/2</sub> (hr) | AUC<sub>0-inf</sub> (hr*ng/mL) | F (%) |
| --- | --- | --- | --- | --- | --- | --- |
| Rat | 10 | 37,900 ± 2,440 | 48 ± 0.0 | 19 ± 1.8 | 3,990,000 ± 657,000 | 52 ± 8.6 |

SC: Subcutaneous; PK: Pharmacokinetics; C<sub>max</sub>: Maximum plasma concentration; T<sub>max</sub>: Time to maximum plasma concentration; T<sub>1/2</sub>: Half-life; AUC: Area under the plasma concentration-time curve; F: Fraction absorbed.

**Table S12.** X-ray diffraction data and refinement.

| | $\alpha 5\beta 1$-MRT1 |
| --- | --- |
| PDB Code | TBD |
| **Data Collection** |  |
| Beamline | NSLSII |
| Wavelength (Å) | 0.9793 |
| Space group | P 2<sub>1</sub> 2<sub>1</sub> 2<sub>1</sub> |
| Cell Dimensions |  |
| a, b, c (Å) | 52.75, 116.23, 158.95 |
| $\alpha$, $\beta$, $\gamma$ (°) | 90, 90, 90 |
| Resolution (Å) | 58.12 - 1.699 (1.72 - 1.7)<sup>a</sup> |
| R<sub>meas</sub> | 0.1009 (3.435)<sup>a</sup> |
| Mean I/$\sigma$(I) | 8.38 (0.73)<sup>a</sup> |
| CC<sub>1/2</sub> | 0.995 (0.321)<sup>a</sup> |
| Completeness (%) | 99.81 (97.23)<sup>a</sup> |
| Multiplicity | 7.3 (6.1)<sup>a</sup> |
| **Refinement** |  |
| Number of reflections | 108184 |
| R<sub>work</sub>/R<sub>free</sub> | 0.1946/0.2159 |
| Number of atoms |  |
| Protein | 6807 |
| Ligands | 405 |
| Solvent | 501 |
| B-factors (Å<sup>2</sup>) |  |
| Protein | 45.18 |
| Ligands | 67.05 |
| Solvent | 48.36 |
| R.M.S. Deviations |  |
| Bond Lengths (Å) | 0.043 |
| Bond Angles (°) | 2.01 |

<sup>a</sup> Values in parentheses are for the highest resolution shell.
CC<sub>1/2</sub>, percentage of correlation between intensities from random half-dataset; PDB, Protein Data Bank;
R.M.S., root-mean-square; R<sub>meas</sub>, redundancy independent R-factor.

